# Adaptive peak tracking as explanation of sparse fossil data across fluctuating ancient environments

**DOI:** 10.1101/2024.10.30.621046

**Authors:** Rolf Ergon

## Abstract

Species that have existed over millions of years have done so because they have been able to track peaks in an adaptive landscape well enough to survive and reproduce. Such optima are defined by the mean phenotypic values that maximize mean fitness, and they are predominantly functions of the environment, for example the sea temperature. The mean phenotypic values over time will thus predominantly be determined by the environment over time, and the trait history may be found in the fossil record. Here, I simulate such a tracking system, using both a basic non-plastic selection model and a univariate intercept-slope reaction norm model. I show how both linear and nonlinear mean phenotype vs. environment functions can be found also from quite sparse and short time series from the fossil record, and I discuss how this methodology can be extended to multivariate systems. The simulations include cases with a constraint on the individual trait values and with other factors than environment influencing the position of the adaptive peak. The methodology is finally applied on a time series of mean phenotypic values in a record of bryozoan *Microporella agonistes* fossils spanning 2.3 million years, using the *∂*^18^*O* measure as proxy for sea water temperature. From as few as nine samples of mean phenotypic values found in the fossil record it was possible to identify a linear mean phenotype vs. environment function, and to predict the continuous mean phenotypic values as functions of time with prediction errors within or just outside the standard errors of the observations. Leave-one-out cross validation gave satisfactory results. It remains to verify predictions for longer time periods without known or investigated fossil data.

## 1 Introduction

Evolutionary time series in the fossil record provide information on phenotypic changes over millions of years. Such time series may consist of quite sparse samples, and they are traditionally modeled by various forms of random walk processes (Hunt, 2006; Voje, 2023), or by Ornstein-Uhlenbeck (OU) processes describing evolution of mean phenotypic values toward fixed or moving peaks in the adaptive landscape (Voje, 2023). Here, the adaptive landscape and adaptive peak metaphors assume that the mean population fitness is a function of possibly multivariate mean phenotypic values, see Pigliucci (2013) for the historical background and a clarifying discussion.

As an alternative to the traditional fossil time series modeling, I propose that adaptive peak tracking may be a useful modeling approach, and that this requires two types of landscape metaphors. First, I assume a fitness landscape where individual fitness is a function of a possibly multivariate set of individual phenotypic values. Second, I assume an adaptive landscape where the mean population fitness is a function of a possibly multivariate set of mean phenotypic values. For simplicity I assume that the position of the individual fitness peak is a function of a dominating environmental factor, and that the adaptive peak may have a somewhat different position owing to non-symmetrical individual fitness or trait distribution functions. I am thus not attempting to estimate the effect of various selective or other factors on the position of adaptive optima (Hansen, 1997), but I just assume that various selective factors over time keep the mean phenotypic values in the vicinity of the adaptive peak, and as an example I will use the relative size of feeding zooids (autozooids) in encrusting cheilostome bryozoans. Liow et al. (2017) studied communities of such organisms spanning more than 2 million years, and an interesting result was that six different species showed temporally coordinated changes in average zooid sizes, suggesting that the different species responded to a common external driver. Liow et al. pointed to sea temperature as a potential common driver, but they found that the available time series of changes in mean zooid size lacked sufficient length to rigorously test such a hypothesis. In my view, a study by means of an adaptive peak tracking model as introduced above is clarifying.

For an introduction, Fig. 1 shows data from Liow et al. (2024) (see a more detailed presentation in Section 4), where the *∂*^18^*O*(*t*) measure is a widely used proxy for seawater temperature (Lisiecki and Raymo, 2005). A natural question is now which type of model we need for explanation of the *log AZ area mean* data in panel B given the *∂*^18^*O*(*t*) data in panel A, and as already mentioned an adaptive peak tracking model is in my view a viable alternative.

**Figure 1.**
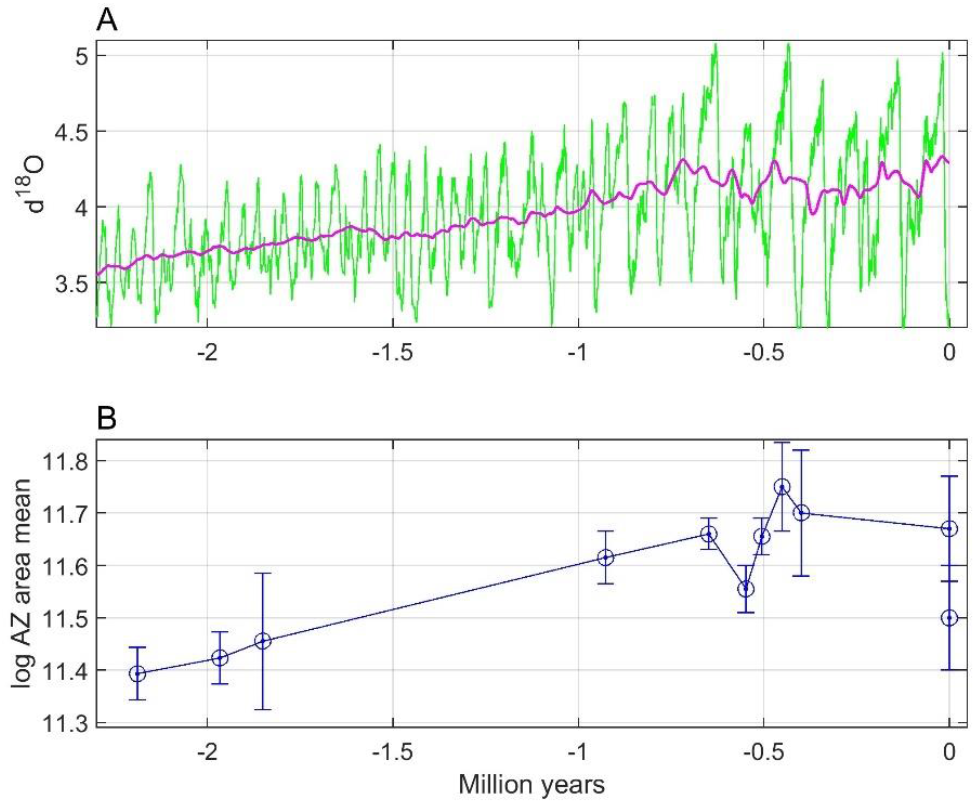
Raw *u*(*t*) = *∂*^18^*O*(*t*) data (panel A, green line) and a centered moving average signal *u*_*filt*_(*t*) computed from a sliding window of length 100 samples along the time axis (panel A, red line, see Section 4 for details), as well as *log AZ area mean* (panel B, circles and dots) with standard errors as in Fig. S2 in Liow at al. (2024). Present time is used as zero point. Note that the moving average signal cannot capture the sharp decline of the *∂*^18^*O*(*t*) value towards time zero.

A block diagram for the proposed adaptive peak tracking model is shown in Fig. 2, and as described above it is based on two types of landscape metaphors. The optimum phenotype that maximizes individual fitness is assumed to be a function *θ*_*i*_(*u, t*) = *f*(*u*(*t*)), where *u*(*t*) is the dominating environmental variable (in the bryozoan example the *∂*^18^*O*(*t*) measure). The optimum mean phenotypic value that maximizes mean fitness is assumed to be a function *θ*_*m*_(*u, t*) = *f*(*u*(*t*)), deviating from *θ*_*i*_(*u, t*) when either the individual fitness function or the trait distribution is non-symmetrical. When both the fitness function and the trait distribution are symmetrical, we will thus have *θ*_*m*_(*u, t*) = *θ*_*i*_(*u, t*) (see Section 2 for an example with Gaussian functions). The tracking process in Fig. 2 is attempting to keep the mean phenotypic value 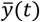 (in the bryozoan example the *log AZ area mean* value) in the vicinity of the optimal value *θ*_*m*_(*u, t*), such that the tracking error *e*(*t*) is kept small. The basic assumption is thus that the tracking process keeps the tracking error small, regardless of the details of this process.

**Figure 2.**
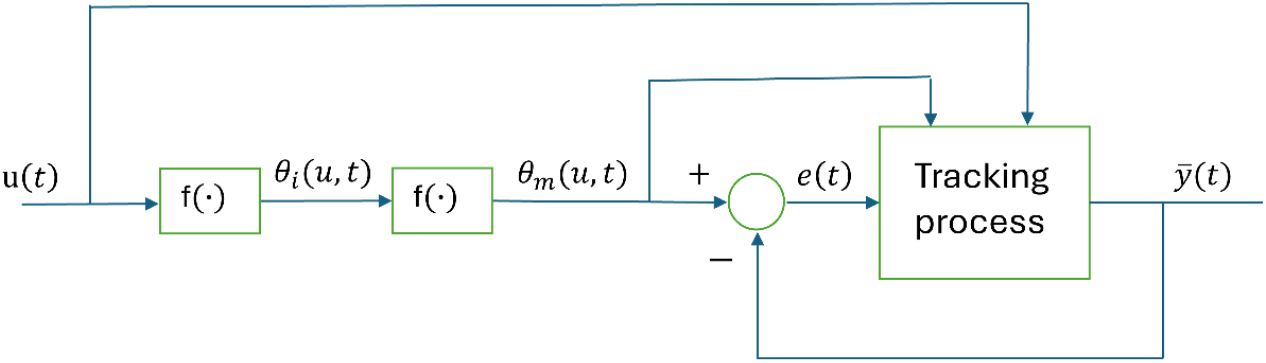
Block diagram for fitness peak tracking system, with the environmental variable *u*(*t*) as input and the mean phenotypic value 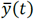 as output. The phenotypic values *θ*_*i*_(*u, t*) and *θ*_*m*_(*u, t*) that maximize individual and mean fitness, respectively, are together with the tracking error *e*(*t*) internal signals in the system. See text for explanations of blocks marked by *f*(·) symbols.

A fluctuating environment as shown in Fig. 1 corresponds to an adaptive landscape where the adaptive peak position fluctuates, and how well the peak is tracked depends on the properties of the evolutionary system. For a population with phenotypic plasticity the tracking error may be kept small at all times, while a population without plasticity may only be able to track a moving average as indicated in Fig. 1. However, a population without plasticity may also keep the tracking error small, provided that additive genetic variances are large. Sparse fossil data will in any case not be informative enough for predictions of strongly fluctuating mean phenotypic values, and a remaining possibility is therefore to predict moving averages over a window size that may be used as prediction parameter. In cases where the fossil phenotypic data are mean values from given time periods, a natural choice is also to use mean environmental data from these time periods as input to the tracking model (see Section 4 for example).

The assumption that the mean phenotypic value stays in the vicinity of the adaptive peak is supported by findings regarding multiple species and traits indicating that “fossil lineages have ascended adaptive peaks and remained at their summits as they shift position through time” (Bell, 2013). An important reference is here Estes and Arnold (2007), who based on a large data set found that “the underlying process causing phenotypic stasis is adaptation to an optimum that moves within an adaptive zone with stable boundaries”, which applied to the data in Fig. 1, panel B, and considering the standard errors given, means that *log AZ area mean* moves between approximately 11 and 12. Using a database over many contemporary populations, Estes and Arnold also found that phenotypic means are typically close to the adaptive peak (46% within 1 phenotypic standard deviation of the optimum, and 65% within 2 standard deviations). Estes and Arnold (2007) based their results on tests with a single displacement of the optimum, using an extensive set of data (Gingerich, 2001), but they acknowledged that the actual optimum might move many times with a small total net displacement. That is a good description of the optimal mean phenotype corresponding to the environmental data in Fig. 1, panel A, with large short-term variations and a rather limited long-time net variation. I thus assume that short-term environmental fluctuations as in, e.g., Fig. 1, panel A, result in variations in optima within a stable adaptive zone (Uyeda et al., 2011). However, contrary to Uyeda et al. I assume a causal tracking system and not a purely stochastic model. In real cases there will of course be stochastic variations in the environmental variable and in the position of the individual fitness peak, and additional stochastic factors that forces the adaptive peak to deviate from the individual fitness peak. Such stochastic variations are included in simulations, but further discussion of this topic is beyond the scope of this paper.

A persistent system excitation in the form of short-term environmental fluctuations as exemplified in Fig. 1, panel A, makes it unlikely that a tracking system as shown in Fig. 2 gets stuck in a local optimum or in a valley in the adaptive landscape. As illustrated in simulations it is more likely that the mean phenotypic values fluctuate around global optima, see however an example of slow adaptation in subsection 3.5 and a discussion in Section 5.

In simulations I assume that the tracking process block in Fig. 2 can be described by use of Lande’s selection theory for populations without plasticity (Lande, 1979), which was also used by Estes and Arnold (2007), but I also include examples with phenotypic plasticity (Lande, 2009). In real cases, however, other mechanisms such as, e.g., adaptive walks in response to mutations may also be involved (Ch. 27, Walsh and Lynch, 2018). It is as already mentioned essential to realize that although knowledge of the details of the tracking process may be interesting, such knowledge is not necessarily needed for finding continuous predictions 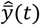 based on sparse measurements of mean phenotypic values 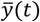.

Under the assumption that the mean trait values have remained centered around the mean fitness peaks as they have shifted position through time, such that the following error *e*(*t*) in Fig. 2 has remained small relative to the width of the underlying mean fitness function, the position of the mean fitness peak as function of the environment, *θ*_*m*_(*u, t*) = *f*(*u*(*t*)), can be reconstructed from fossil samples of 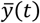, and provided that this function is sufficiently smooth this can be done also when the fossil samples are sparse and at irregular time intervals. For the bryozoan data in Fig. 1, it turns for example out that the mean trait value 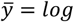 *AZ area mean* as function of the environment *u* = *∂*^18^*O* can be modeled by a straight line, from which follows continuous predictions of 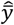 as function of time (see Section 4 for details).

The prediction method outlined above is supported by simulations in Section 3, with the theoretical background given in Section 2. The simulations are based on a reaction norm model where I use the *∂*^18^*O*(*t*) measure as environmental cue, and where the phenotypic plasticity may be set to zero. This gives a connection between the simulations and the real bryozoan data case in Section 4, in the sense that the same type of large but short-time fluctuations in the input signal is used. Since both *∂*^18^*O*(*t*) and 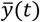 in the simulations have large variances, and since data from the fossil record often are mean values over both populations and time (Liow et al., 2024), I will in the simulations use moving average filtered versions of these variables. The simulations are intended to show that mean phenotype vs. environment functions can be predicted from sparse data also in cases where the population in shorter time periods have quite low mean fitness. The simulations also show that 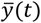 fluctuates around the optimal value *θ*_*m*_(*u, t*), as discussed above, although in some cases with a minor time lag. In order to show that *θ*_*m*_(*u, t*) may be different from *θ*_*i*_(*u, t*), a case with a constraint on individual trait values is included. In addition to theory necessary for the simulations, as given in Section 2, the theory is extended to cover multivariate cases.

In Section 4, the prediction method is applied on recently presented data from an already introduced field study of the bryozoan species *Microporella agonistes*, based on a fossil record over 2.3 million years (Liow et al., 2024). Parameters in a linear mean phenotype vs. environment function are here estimated from as few as 9 data points at irregular sampling times. This is done using mean values of the *∂*^18^*O*(*t*) measure in the time periods of collected fossil data, but also, and with better results, using moving average mean environmental data. Finally follows a summary and discussion in Section 5.

## 2 Theory

### 2.1 A basic univariate reaction norm case

For simplicity and for use in simulations I will initially assume that mean fitness is determined by a univariate phenotypic mean value 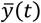, and that the adaptive landscape thus is two-dimensional. I will later extend some essential results to multivariate cases, and the simulations will also include examples where other factors than the physical environment are involved.

In the simulations I will make use of an individual intercept-slope reaction norm model with parameters *a*(*t*) and *b*(*t*) as given by

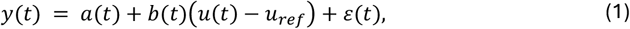

where *b*(*t*) and the variance *G*_*bb*_ = *var*(*b*(*t*)) in some cases are set to zero. Here, *u*(*t*) is the environmental cue, while *ε*(*t*) is assumed to be normally distributed with mean 0 and variance 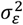 among individuals in every generation. In Eq. (1), *u*_*ref*_ is a reference environment defined as the environment at which the phenotypic variance is at a minimum (Lande, 2009). For a given population this will result in the mean reaction norm

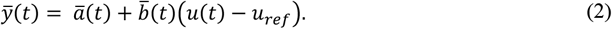

The optimal individual phenotypic value is in general a function of the environment,

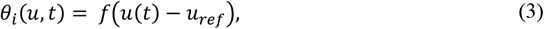

in the linear case *θ*_*i*_(*u, t*) = *α* + *β*(*u*(*t*) − *u*_*ref*_), where *α* and *β* are constant parameters.

In the simulations I will assume an individual fitness function

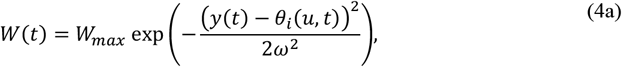

which assuming that *y*(*t*) is normally distributed gives the mean fitness in a population (Lande, 2009; Estes and Arnold, 2007)

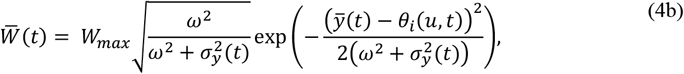

where 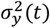 is the variance of *y*(*t*) according to Eq. (1). The mean fitness will thus be maximized when 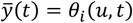. Note that this is an example with symmetrical individual fitness and trait distribution functions, such that optimal individual and mean phenotypic values are equal, i.e., *θ*_*m*_(*u, t*) = *θ*_*i*_(*u, t*). In Subsection 3.3 I will, however, also include a case where *θ*_*m*_(*u, t*) ≠ *θ*_*i*_(*u, t*).

Under the assumptions of non-overlapping generations and random mating in a large population the evolution of the mean reaction norm parameters is governed by (Lande, 1979)

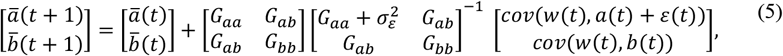

where *G*_*aa*_, *G*_*bb*_ and *G*_*ab*_ are variances and the covariance of the reaction norm parameters, and where 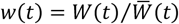 is the relative fitness. In a stationary stochastic environment, the optimal reaction norm slope is 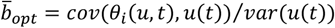 (Lande, 2009).

When the mean phenotypic trait in a population is tracking a moving optimum *θ*_*m*_(*u, t*) it will be given by 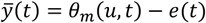, where *e*(*t*) is the tracking error (see block diagram in Fig. 2). This means that the parameter values 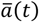 and 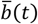 are forced to evolve such that 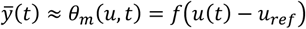, i.e., such that the mean fitness according to Eq. (4b) is kept close to the optimum and that the tracking error is small. In feedback control terminology it is thus the mean trait tracking error 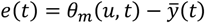 that via mean fitness is the driving force in the adaptive process. It follows from Eq. (5) that the tracking system in Fig. 2 has what in feedback control terminology is called integral action (Åström and Murray, 2008), which means that the tracking error in a stationary case with a fixed adaptive peak will asymptotically go to zero. As shown in simulations in Section 3 the parameters in the 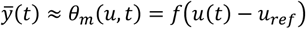 function can be estimated also from rather few fossil data points 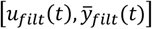 at irregular sampling times, where 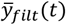 and *u*_*filt*_(*t*) are moving average filtered versions of often quite noisy signals. In the real data case in Section 4 the parameters in *θ*_*m*_(*u, t*) are also found from data points 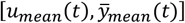 for the time periods of collected fossil data. Note that even though 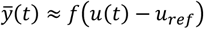, such a model is not a reaction norm, i.e., the reaction to a sudden change Δ*u* in *u*(*t*) will still be a sudden change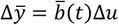.

The details in the tracking process in Fig. 2 as used in the simulations are described by Eqs. (2), (4) and (5), but also if the parameters in those equations were known they could not be used for computation of predictions 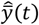 based on only a few fossil samples at irregular time points. The remaining possibility for prediction of 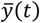 is to assume good enough tracking such that 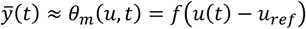, which as shown in the simulations as well as in the real data case makes it possible to estimate the parameters in the *θ*_*m*_(*u, t*) function also from few 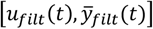 or 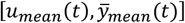 data points. Based on those estimates and known values of *u*(*t*) it is then possible to predict 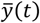.

### 2.2 Influence of other factors than the environment

Eqs. (2) and (3) assume that u(*t*) is the only factor that affects the mean phenotypic value 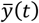 and the optimal mean phenotypic value *θ*_*m*_(*u, t*) = *θ*_*i*_(*u, t*), respectively. When other factors affect 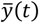, the tracking system in Fig. 2 will force the parameters 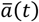 and 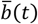 in Eq. (2) to evolve such that these factors are compensated for, and the result will be seen in the mean phenotype vs. environment function that can be found from data, and then most likely in the form of nonlinearities as shown in a simulation example in Subsection 3.3. When other factors affect *θ*_*m*_(*u, t*) it is again the 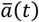 and 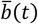 parameter values that must compensate, also then most likely in the form of nonlinearities as shown in Subsection 3.4. A linear mean phenotype vs. environment function thus indicates that no other factors than the environment affect the fitness vs. environment function, although there may of course be several other factors that compensates the effects of each other. And in cases where *θ*_*i*_(*u, t*) is a nonlinear function there may be factors that make *θ*_*m*_(*u, t*) linear.

### 2.3 Multivariate cases

When the mean phenotypic trait in a univariate population is tracking the optimal phenotypic value, the mean phenotypic trait can as shown above be described by 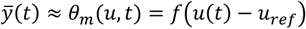, i.e., by the adaptive peak vs. environment function. Here, it is again important to note that we do not need to know the details of the tracking process in Fig. 2.

The univariate theory can be extended to multivariate cases, and although we in practice do not need to know the underlying fitness function, we may for a theoretical discussion use an example with two phenotypic traits and the bivariate fitness function (Ch. 4, Johnson & Wichern, 2008)

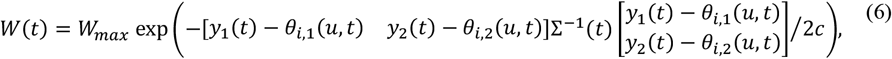

where Σ(t) = cov(*y*_1_(*t*), *y*_2_(*t*)) is the variance-covariance matrix and where *c* is a scaling factor controlling the ‘width’ of the fitness function. In order to keep the mean traits in the vicinity of the adaptive peak, they must in this case continuously evolve such that 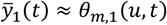 and 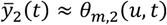, and we will thus find

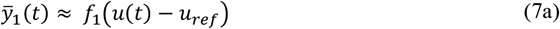

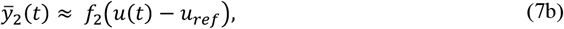

where the moving adaptive peak vs. environment functions may be linear or nonlinear, and where the parameters can be estimated from rather few fossil data points at irregular sampling times. Here, the essential point is that 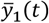 and 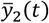 in this example are tracking a peak in a three-dimensional adaptive landscape, and that this tracking process can be decomposed into two univariate tracking processes. We may thus consider the projections of the bivariate fitness function onto orthogonal axes for the two phenotypic traits, and such a projection approach may be extended to systems with higher dimensions, as discussed in Goodnight (2013). Note that we here are only interested in the *θ*_*m*,1_(*u, t*) and *θ*_*m*,2_(*u, t*) functions that define the position of the adaptive peak, and that we do not need to know the shape of the adaptive landscape around the peak.

## 3 Simulations

### 3.1 Linear reaction norm example

A system with a linear mean reaction norm 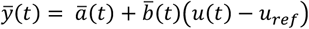 was simulated with use of the 2,115 available samples of *u*(*t*) = *∂*^18^*O*(*t*), and with population size 400. These samples go 5.3 million years back in time with various sampling intervals (Lisiecki and Raymo, 2005), but here I just assume 2,115 nonoverlapping generations (however, see Section 5 for a discussion on relevance for fossil data). At each new generation random individual samples around 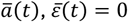 and 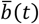 were drawn from normal populations with the given variances. The fitness function in Eq. (4a) had parameter values *W*_*max*_ = 1 and *ω*^2^ = 50. The optimal mean phenotypic value was in a first linear case chosen as *θ*_*m*_(*u, t*) = *θ*_*i*_(*u, t*) = *u*(*t*) − 2.9, where *θ*_*m*_(*u, t*) thus and for clarity of presentation is assumed to be a deterministic function, and where 2.9 is the mean value of *∂*^18^*O*(*t*) around *t* = 0 (however, see results with added white noise below).

As it is not realistic that *u*(*t*) and *θ*_*m*_(*u, t*) are purely deterministic signals, correlated white noise with variances 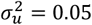 and 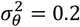, respectively, were added to *θ*_m_ (*u, t*) and *u*(*t*). The covariance was *σ*_*θ,u*_ = 0.025, see Lande (2009) for a possible explanation.

In a first case without phenotypic plasticity, the reaction norm parameter variances and covariance were chosen as 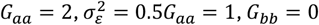 and *G* = 0, while the initial parameter values were 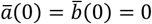. Since 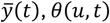 and *u*(*t*) = *∂*^18^*O*(*t*) have large variances, I used moving average filtered signals 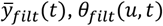 and *u*_*i,filt*_(*t*) in some of the plots. These were obtained by moving window filters with window size 100, in MATLAB notation 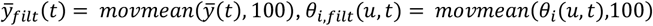 and *u*_*filt*_(*t*) = *movmean*(*u*(*t*), 100). Simulation results in the linear case without plasticity are shown in Fig. 3, panels A to E. Note that 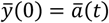 follows the main trend in *u*(*t*), and that the fluctuations are somewhat delayed compared to the fluctuations in *u*(*t*) (panel A).

**Figure 3.**
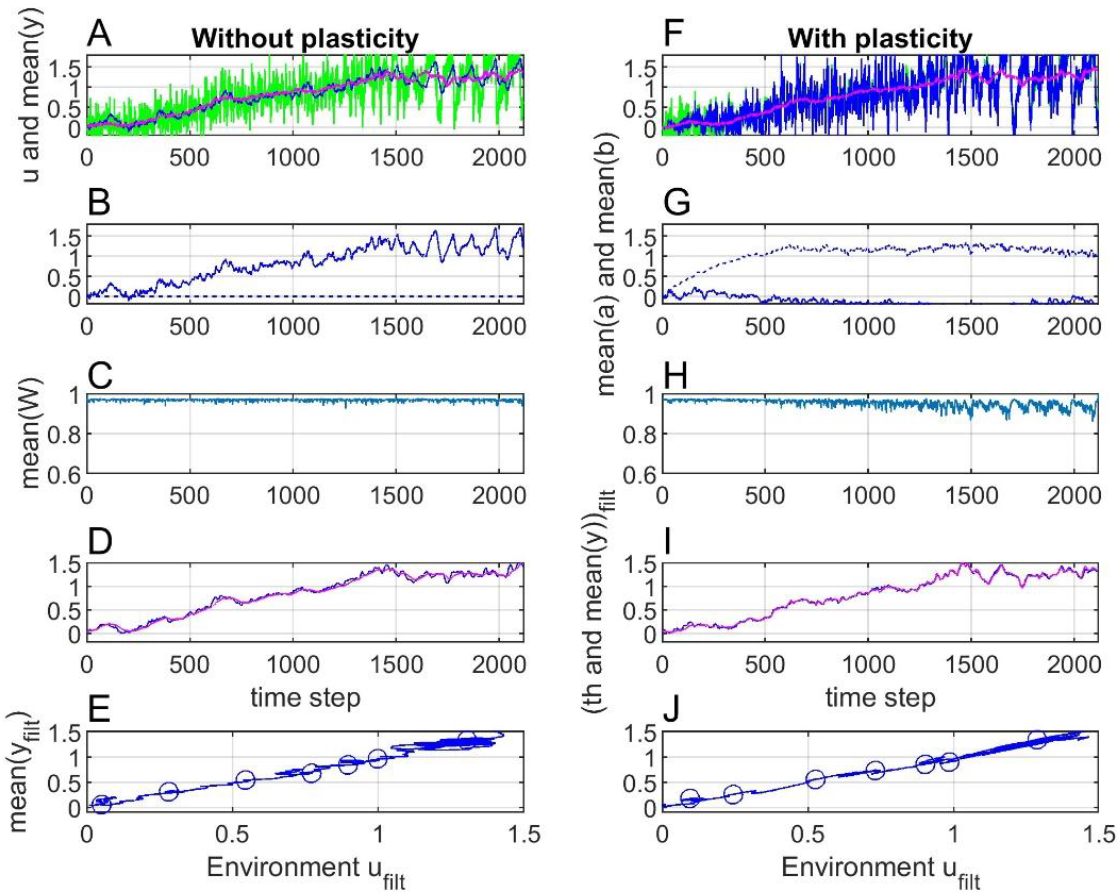
Simulation results with the linear *θ*_*m*_(*u, t*) = *θ*_*i*_(*u, t*) = *u*(*t*) − 2.9 function, without plasticity (panels A to E) and with plasticity (panels F to J). Panels A and F show *u*(*t*) (green lines) and 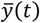 (blue lines), while the red line is *u*_*filt*_(*t*). Panels B and G show 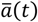 (solid lines) and 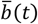 (dashed lines), while panels C and H show mean fitness 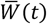. Panels D and I show *θ*_*i,filt*_(*u, t*) (blue lines) and 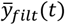 (red lines). Panels E and J show that 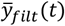 as function of *u*_*filt*_(*t*) in both cases is the same straight line, with values for samples 200, 400, 600, 800, 1000, 1200 and 1400 shown by circles.

In a second case with phenotypic plasticity, the reaction norm parameter variances and covariance were chosen as 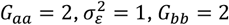 and *G*_*ab*_ = 0, while the initial parameter values were 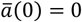 and 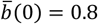. Simulation results in the linear case with plasticity are shown in Fig. 3, panels F to J. As follows from the reaction norm equations (1) and (2), the variance 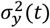 will increase with *u*(*t*) − *u*_*ref*_, and as a result the mean fitness will be reduced (panel H).

In both cases (without and with plasticity) 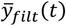 tracks *θ*_*m,filt*_(*u, t*) = *θ*_*i,filt*_(*u, t*) closely (panels D and I), such that the mean fitness is kept at a high level (panels C and H). This results in linear functions 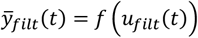 (panels E and J), with seven discrete values marked by circles. Using *u*_*ref*_ = *u*_*filt*_(0) = 2.9, these discrete values could be used for determination of the parameters 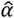 and 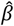 of this line by an ordinary least squares method, and thus to find predictions 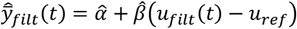, see examples in Section 4.

### 3.2 Nonlinear reaction norm example

Fig. 4 shows results as in Fig. 3, but with the nonlinear adaptive peak vs. environment function *θ*_*m*_(*u, t*) = *θ*_*i*_(*u, t*) = (*u*(*t*) − 2.9) + 4(*u*(*t*) − 2.9)^2^. Note that the mean fitness now is reduced (panels B and F), especially in the case without plasticity. The signal 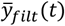 is also in this case tracking *θ*_*filt*_(*u, t*) well, although with a noticeable time lag (panels C and G). Here, 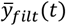 as function of *u*_*filt*_(*t*) is the same second order function in the different cases (panels D and H).

**Figure 4.**
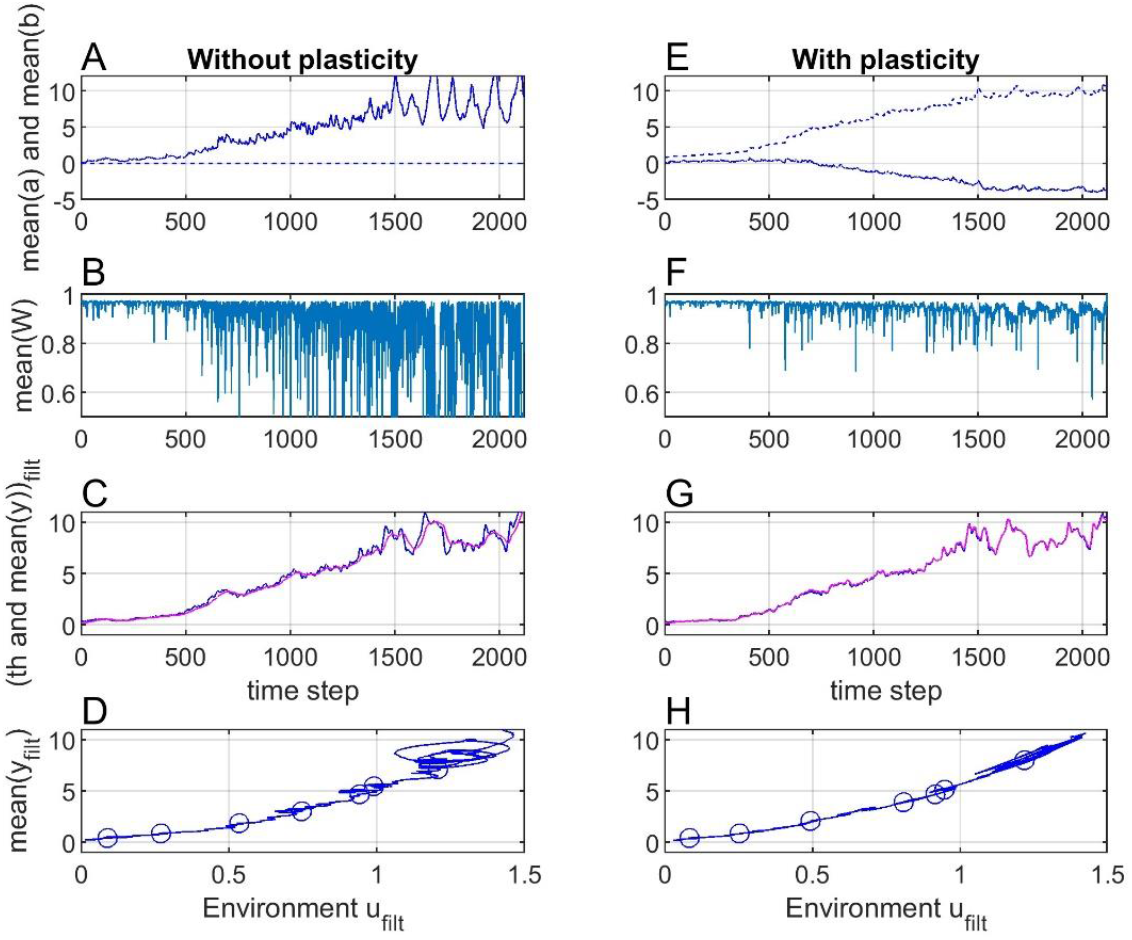
Simulation results as in Fig. 3, but with *θ*_*m*_(*u, t*) = *θ*_*i*_(*u, t*) = (*u*(*t*) − 2.9) + 4(*u*(*t*) − 2.9)^2^. The discrete values in panels D and H could here be used to determine the nonlinear function 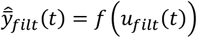 by means of any suitable fitting method.

### 3.3 Example with phenotypic constraint

With a constraint on the individual phenotypic values the distribution will no longer be normal, and the adaptive peak *θ*_*m*_(*u, t*) will therefore deviate from the individual fitness peak such that Eq. (4b) no longer is correct. The mean trait 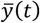 will, however, still track the adaptive peak that maximizes mean fitness. When there for example is an upper limit for the individual phenotypic trait values as shown in Fig. 5, the 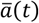 and 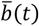 parameters will evolve such that 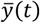 deviates from *θ*_*m*_(*u, t*) with results as shown in Fig. 6, panels A to D. The effect of the constraint is clearly seen in panel C in Fig. 6, in that 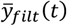 does not follow *θ*_*i,filt*_(*u, t*), but instead follows *θ*_*m,filt*_(*u, t*) (not shown) such that the mean fitness as shown in panel B is maximized. The effect is also seen by a comparison of panels A in Fig. 6 and panel G in Fig. 3, in that 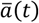 evolves into a large value, such that 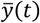 without constraints on *y*(*t*) would become larger than *θ*_*m*_(*u, t*). Note that the mean fitness is increased compared to the results in Fig. 3, panel H, because the variance 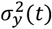 is reduced. Also note that the mean phenotype vs. environment functions in panel D now is nonlinear in a way that with knowledge of the biological system possibly could be interpreted in a meaningful way. Note that individual trait values will have a larger variance than the mean trait value, such that the constraint in Fig. 5 has an effect before 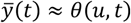 reaches the upper limit.

**Figure 5.**
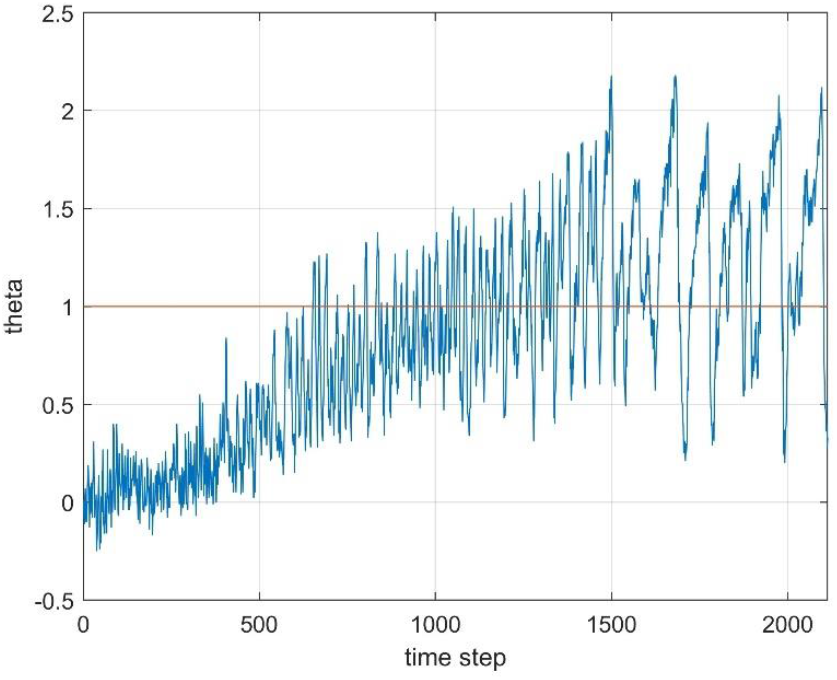
The linear function *θ*_*m*_(*u, t*) = *u*(*t*) − 2.9 as used for the results in Fig. 3 (blue line) and an upper constraint for the individual trait values (red line).

**Figure 6.**
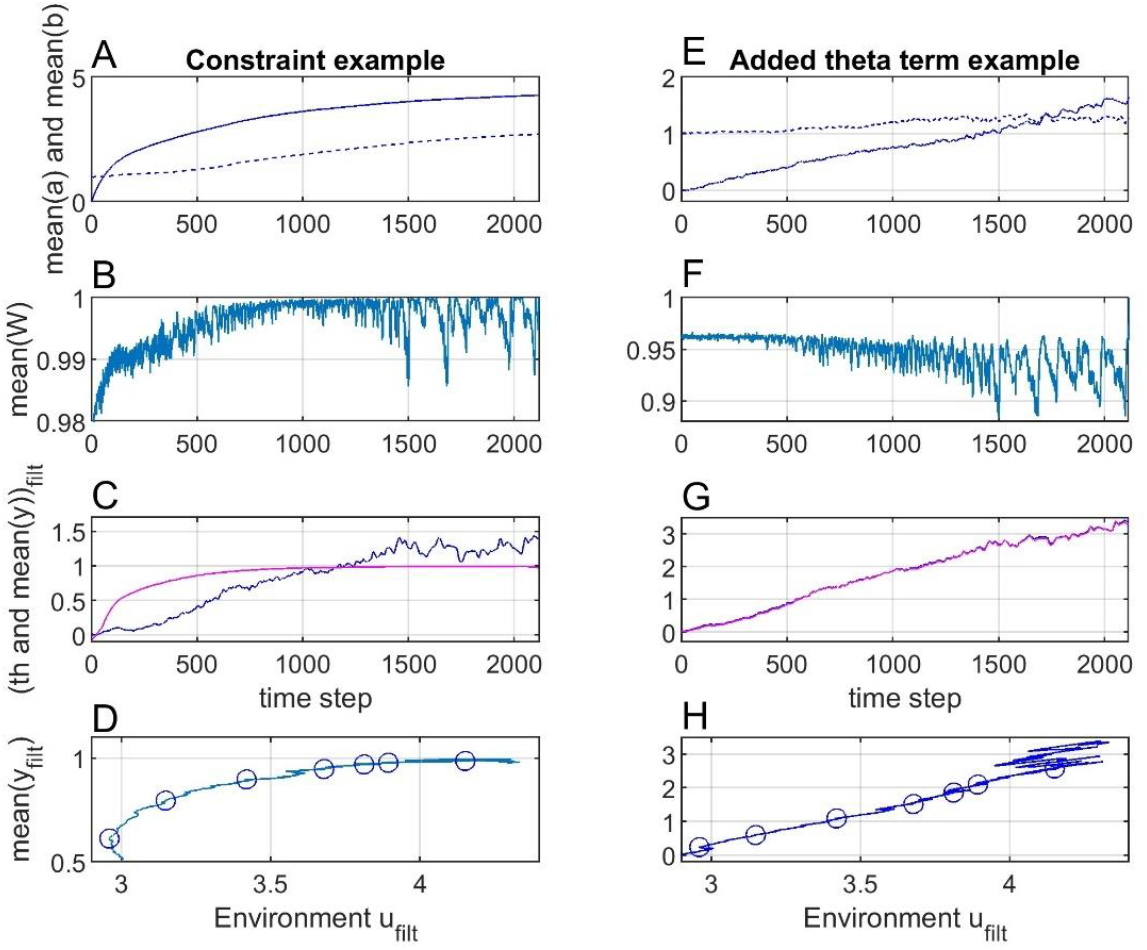
Panels A to D show simulation results as with plasticity in Fig. 3, but with a constraint on the individual phenotypic trait values as shown in Fig. 5. Panels E to H show simulation results as with plasticity in Fig. 3, but with *θ*_*m*_(*u, t*) = *θ*_*i*_(*u, t*) = *u*(*t*) − 2.9 + 2*t*/2115.

### 3.4 Example with additional term in *θ*(*u, t*)

The optimal value *θ*_*m*_(*u, t*) that maximizes mean fitness may not be a function of only the environment, and as an example Fig. 6, panels E to H, shows results as in Fig. 3, panels F to J, but with *θ*_*m*_(*u, t*) = *θ*_*i*_(*u, t*) = *u*(*t*) − 2.9 + 2*t*/2115. Note that panel H now shows a nonlinear function.

### 3.5 Example without plasticity, but with changed parameter values

Simulations as in Fig. 3, panels A to E, were repeated but with changed parameter values. First, *G*_*aa*_ and 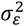 were reduced from 2 and 1 to 0.2 and 0.1, with results as in Fig. 7, panels A to E. As seen in panels A and F the mean phenotypic value 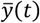 cannot now follow the adaptive peak *θ*_*m*_(*u, t*)), and as a result the mean phenotype vs. environment function in panel E becomes non-linear. However, owing to the broad fitness function with *ω*^2^ = 50, the mean fitness is still close to one.

**Figure 7.**
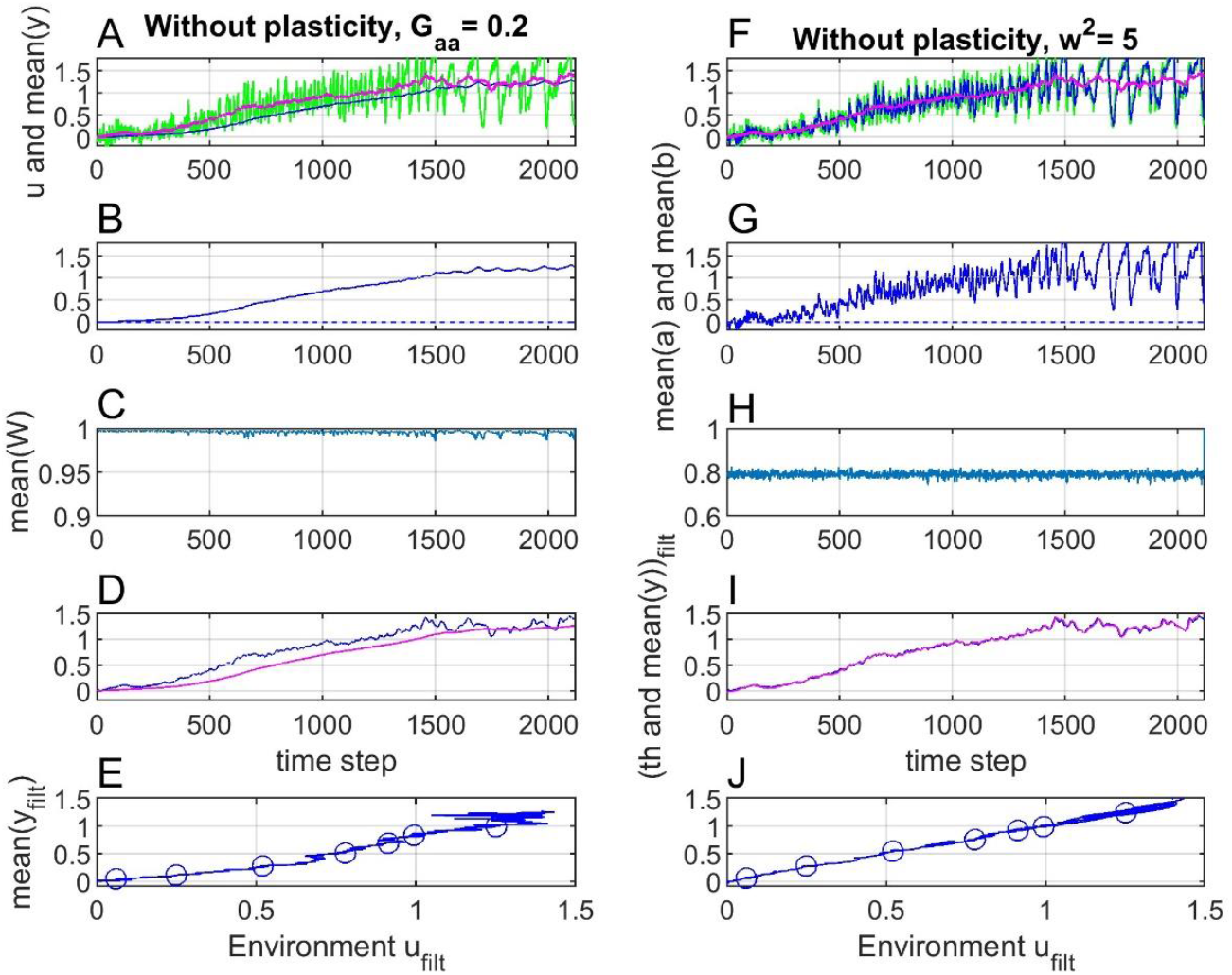
Panels A to E show simulation results as in Fig. 3 with plasticity, but with *G*_*aa*_ and 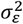 reduced by a factor 10. Panels F to J show simulation results as in Fig. 3 with plasticity, but with the width of the fitness function reduced from *ω*^2^ = 50 to *ω*^2^ = 5.

Second, with *G*_*aa*_ = 2 and 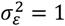 as in Fig. 3, the width of the fitness function was reduced by reducing *ω*^2^ from 50 to 5, with results as in Fig. 7, panels F to J. The mean phenotypic value still follows the adaptive peak but owing to the narrow fitness function the mean fitness is clearly reduced (panel H).

## 4 Real data case

Liow et al. (2024) recently presented results from a field study of the bryozoan species *Microporella agonistes*, based on a fossil record over 2.3 million years (see Fig. 1 for some results). The fossils were collected from colonies in New Zealand cliffs, and it is especially interesting that the same species still live in the waters near those cliffs. In a project running over several years, Liow and her coworkers collected 985 fossil colonies with a large total number of individuals and developed an artificial intelligence tool for data retrieval by use of pictures from a scanning electron microscope. This image analysis system can rapidly assess three well defined phenotypic traits; autozooid (AZ) area, ovicell (OV) area and autozooid shape, and as seen from time series in Fig. S2 in Liow et al. (2024) the two AZ and OV area mean trait values increase with time except for a drop at present time owing to an extreme *∂*^18^*O*(*t*) value.

Since the *log AZ area mean* and *log OV area mean* data in Fig. S2 are linearly dependent, I will here focus on the former. The last two data points in this time series are found from bryozoans from recent times with extreme *∂*^18^*O*(*t*) values. (Fig. S9 in Liow et al.,2024). As done in Fig. 6 in Liow at al. (2024) these data points are therefore excluded from the analysis (see Section 5 for a discussion). The fossil data used in the analysis are thus collected from nine periods back in time according to Table 1, which for the purpose of comparison with moving average mean data also includes the time window size for 100 samples. The reason for use of the natural logarithm of the AZ area is that it gives normally distributed data (Di Martino and Liow, 2021).

**Table 1.** Fossil collection time periods data.

| sample number | mean sample time point | window size | window size for 100 samples |
| --- | --- | --- | --- |
| 1 | −2.1895 | 0.203 | 0.25 |
| 2 | −1.9660 | 0.102 | 0.25 |
| 3 | −1.8505 | 0.049 | 0.25 |
| 4 | −0.9265 | 0.019 | 0.2 |
| 5 | −0.6485 | 0.055 | 0.2 |
| 6 | −0.5480 | 0.030 | 0.2 |
| 7 | −0.5055 | 0.055 | 0.1 |
| 8 | −0.4510 | 0.054 | 0.1 |
| 9 | −0.3990 | 0.050 | 0.1 |

Since the *log AZ area mean* values are given as mean values from samples within time windows as given in Table 1, it is natural to use mean values *u*_*mean*_(*t*) of *∂*^18^*O* in the same windows. Fig. 8 shows *u*(*t*) = *∂*^18^*O*(*t*) and *y*(*t*) = *log AZ area mean* (*t*), as well as predictions 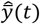 based on least squares fitting of a straight line *y* = *a* + *b*(*u* − 3.6) in a plot over *y*(*t*) as function of *u*_*mean*_(*t*). Note the rather large deviations from the straight line in panel C, but also that there are no indications of a non-linear function. The slope of the prediction line is 0.19, (as in Fig. 6 in Liow et al., 2024) and the mean squares prediction error is 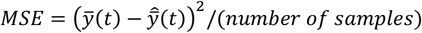 is *MSE* = 0.0089 (Table 2).

**Table 2.** Results with different prediction models.

| | Model with use of $u_{mean}$ as in Fig. 8 | Model with use of $u_{filt}$ as in Fig. 9 |
| --- | --- | --- |
| $b$ | 0.19 | 0.56 |
| $MSE$ | 0.0089 | 0.0014 |
| $b_{CV}$ | - | 0.56 |
| $MSE_{CV}$ | - | 0.0020 |

**Figure 8.**
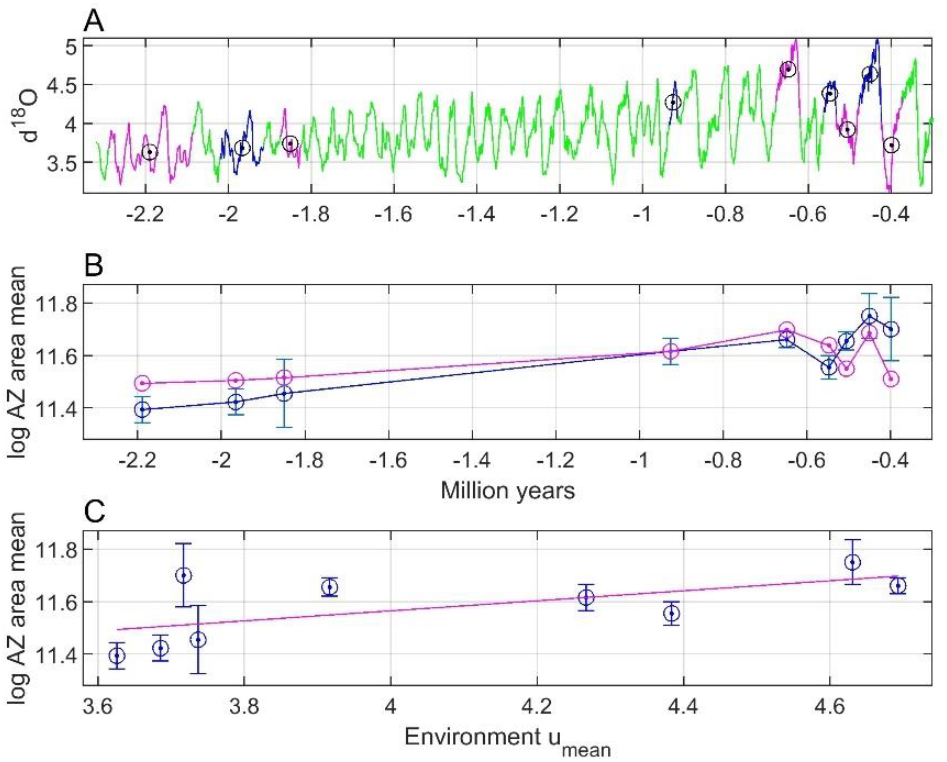
Panel A shows raw *∂*^18^*O*(*t*) data (green line) with values in fossil collection time windows marked by red and blue lines, and with mean values marked by circles with dots. Panel B shows *log AZ area mean* with error bars as found from Fig. S2 in Liow at al. (2024) (blue circles with dots and blue line), as well as predictions (red circles with dots and red line). Panel C shows prediction line (red) computed by the ordinary least squares method. See prediction results in Table 2.

Fig. 9 shows the corresponding results based on centered moving average filtering of *∂*^18^*O*(*t*). The optimal moving window size was 100 samples, resulting in prediction line slope 0.56 and prediction error *MSE*_*min*_ = 0.0014. Note that the deviations from the straight line in panel C are clearly smaller than in Fig. 8. Also note that it from the way the *∂*^18^*O*(*t*) data are organized in Lisiecki and Raymo (2005) follows that a fixed window size given by number of samples correspond to different time durations as shown in Table 1.

**Figure 9.**
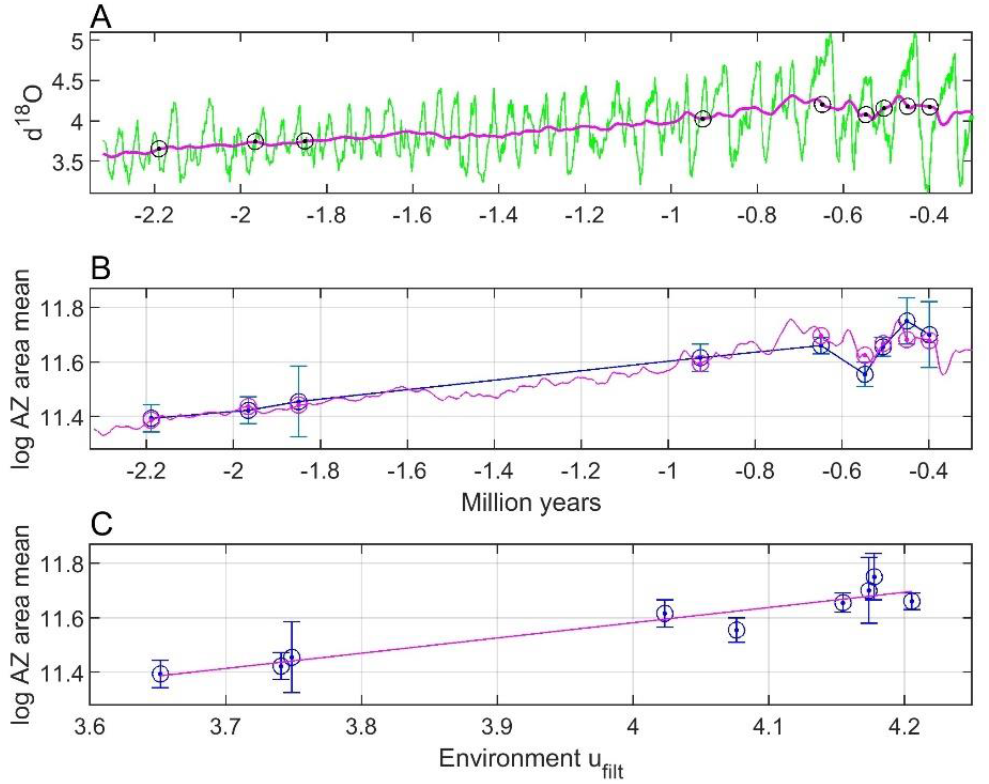
Results as in Fig. 8, but based on centered moving average filtering of *∂*^18^*O*(*t*) with window size 100 samples, and with continuous predictions of *log AZ mean area* values (red line in panel B). See prediction results in Table 2.

The prediction results in Fig. 9 were validated by leave-one-out cross validation, with results as in Fig. 10 and Table 2.

**Figure 10.**
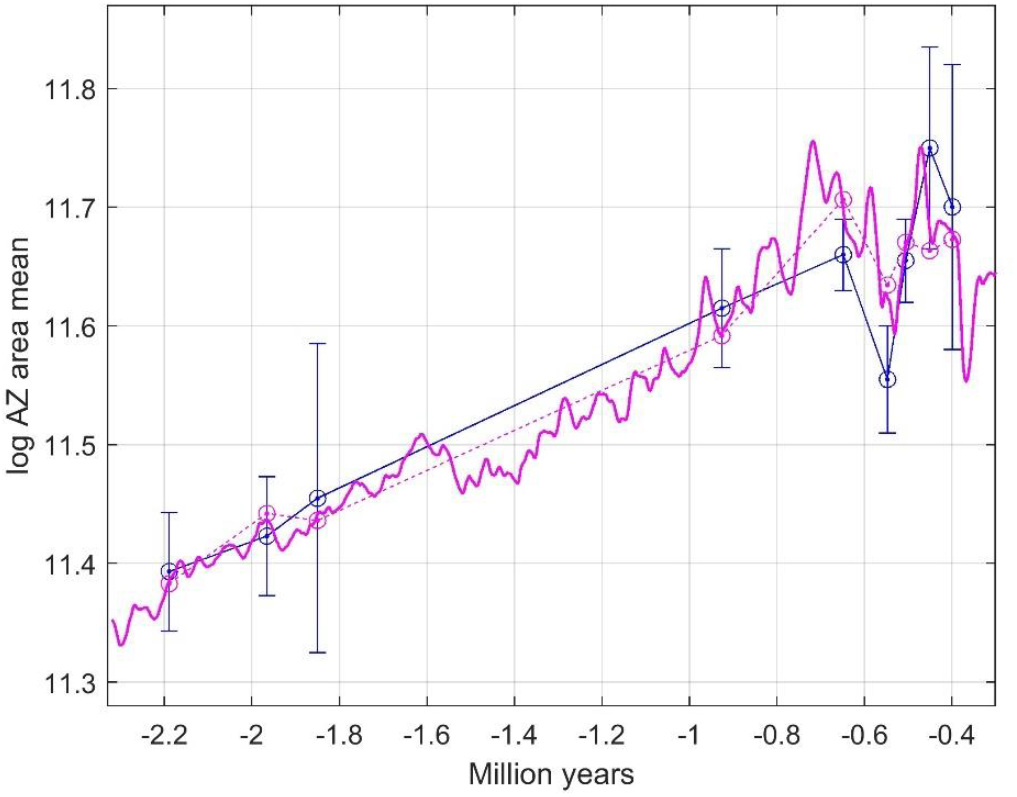
Results as in Fig. 9, panel B, and for leave-one-out cross validation. Solid red line shows prediction results, while dashed red line and red circled with dots show validation results for the left-out samples. See prediction and validation results in Table 2.

As shown in Figs. 1, 8, 9 and 10 the measurements of *log AZ area mean* have various standard errors. Repeated simulations with the model as used for Fig. 10, and with discrete values drawn from normal distributions with standard deviations equal to the standard errors, gave results as in Fig. 11. As can be seen in the upper panel there is ample space for nonlinear prediction functions, although the plot in Fig. 9, panel C, does not indicate a nonlinear function. From the histogram in the lower panel follows that the prediction line slope value *b* = 0.56 has a standard error of approximately SE = 0.10.

**Figure 11.**
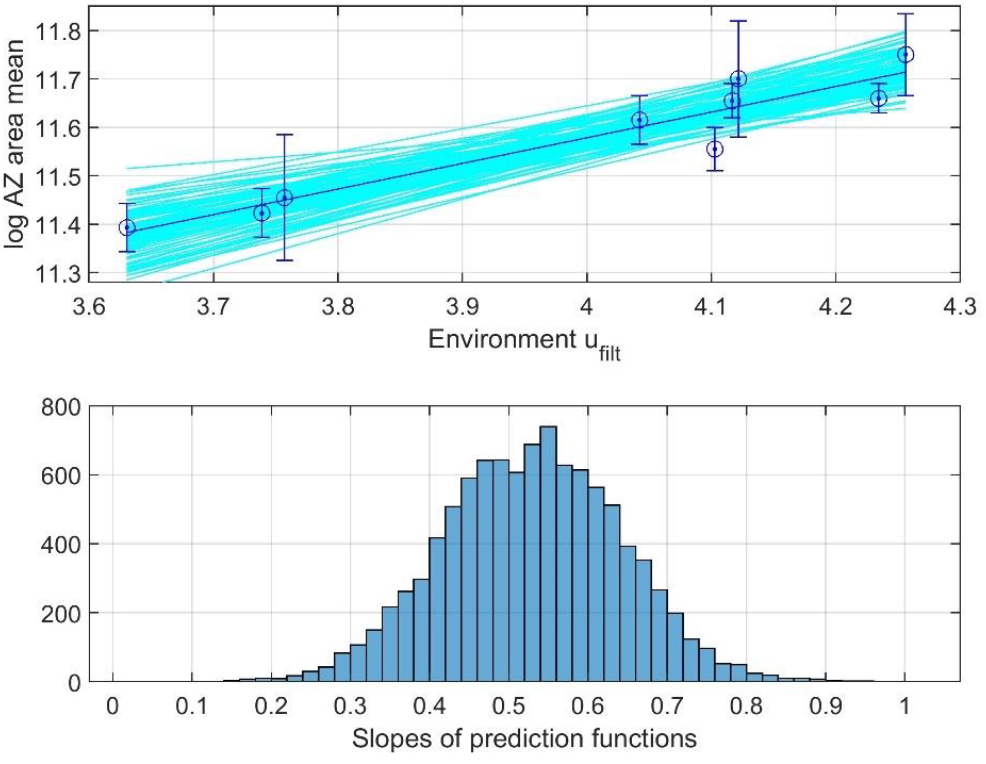
Upper panel shows linear prediction lines for *log AZ area mean* from 100 simulations with measurement values drawn from normal distributions of 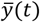 with the given standard errors (cyan lines), and for the mean prediction line corresponding to Fig. 10 (blue line with error bars). Lower panel shows histogram over slopes of fitted prediction lines as in upper panel, but for 10,000 simulations, indicating that the prediction line slope is *b* = 0.56 ± 0.10.

## 5 Summary and Discussion

### 5.1 Main points

In this paper I make use of two landscape metaphors. First, an individual fitness landscape where the peak of the individual fitness as function of individual phenotypic values moves with changes in the environment. Second, an adaptive landscape where the peak of mean fitness as function of mean phenotypic values moves with changes in the environment. The adaptive peak will essentially follow the individual fitness peak, but with deviations caused by non-symmetrical fitness and trait distribution functions and possibly also owing to other causes.

I also make some other fundamental assumptions. First, species that have existed for millions of years in a fluctuating environment have by means of various selection mechanisms been tracking a limited movement of an optimum in the adaptive landscape, and they have done so well enough to survive and reproduce. This assumption is supported by results in Estes and Arnold (2007) and other references in the introduction. Second, the location of the individual fitness peak is a simple continuous function of a dominating environmental variable, e.g., the temperature, such that the movement of the peak in the adaptive landscape is continuous without sudden jumps. Third, the dominating environmental variable fluctuates in a way that makes it unlikely that the mean phenotypic value gets stuck in local minima or valleys in the adaptive landscape.

Under the assumptions given, two main conclusions can be drawn from the results in this article: First, adaptive vs. environment functions, and thus approximate mean phenotype vs. environment functions can be found from sparse and short fossil data, provided that they are sufficiently spread over the time period of interest. As demonstrated in the real data example in Section 4, the environmental data used for prediction of mean phenotypic values may be means from time periods of fossil collection or a moving average mean. Second, prediction of mean phenotypic values does not require detailed knowledge of the tracking process in Fig. 2, i.e., neither parameter values in an underlying reaction norm or adaptive walk model nor the fitness functions need to be known. Also note that it is not necessary to find explicit mathematical adaptive peak vs. environment functions, predictions may instead be found by use of spline interpolation and similar methods.

Even if the position of the individual fitness peak is a linear or other simple function of the environment, the adaptive peak vs. environment function may be less simple. This is because non-symmetrical fitness functions and/or trait distributions may drive the adaptive peak away from the individual fitness peak. A closer look at the tracking process might thus be necessary for interpretations of nonlinear mean phenotype vs. environment results. A deeper understanding of the adaptive landscape may also be needed in cases with slow adaptation, where the mean phenotypic value can track the adaptive peak only with a large time lag.

### 5.2 Discussion of simulation results

The main purpose of the simulations is to show that the mean phenotypic values over time as functions of mean environment over time (panels E and J in Fig. 3 and panels D and H in Fig. 4) are determined by the adaptive peak vs. environment function, and thus independent of the details of the tracking process in Fig. 2. The variances of the reaction norm parameters are chosen such that there is a clear difference in the evolutionary responses between cases without and with plasticity. If these variances are increased by a factor ten (or a factor one hundred) this difference very much disappears in that 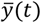 in both cases appear to follow the adaptive peak *θ*_*m*_(*t*) = *f*(*u*(*t*)) instantaneously, but the mean phenotypic values over time as functions of mean environment over time are essentially unchanged.

Figs. 3 and 4 show that although a population without plasticity may not be able to track rapid changes in environment it may track a moving average quite well, and that the resulting mean phenotype prediction functions, linear or non-linear, will be the same as for a population with plasticity. Note that the mean plasticity slope with a linear fitness peak vs. environment function approaches an optimal value (Fig. 3, panel F), such that the linear mean phenotype vs. environment function seen in panel J can be interpreted as the result of adaptive phenotypic plasticity. However, a comparison with results without plasticity (panel E) shows that this interpretation may be wrong.

Fig. 6, left panels, illustrates that non-symmetrical trait distributions result in non-linear mean phenotype vs. environment functions also in cases where the individual fitness peak position is a linear function of a dominating environmental factor. Fig. 6, right panels, illustrates that other factors than the environment may contribute to non-linearities in the mean phenotype vs. environment function.

Panel 7, left panels, illustrates a case with slow adaptation as in, e.g., Toljagic et al. (2018), where a mean trait value lags behind changes in the adaptive peak. The result is a non-linear mean phenotype vs. environment function, but also in this case mean phenotypic values over time can be predicted from a limited number of samples. Panel 7, right panels, shows that a narrow fitness function, as must be expected results in reduced mean fitness.

### 5.3 Discussion of the real data case

The time series in the real data case in Section 4 has samples from around 2 million years ago and from around 0.5 million years. Although it would have been nice with additional data from around 1.4 and 0.25 million years ago the available data indicate a linear adaptive peak vs. environment function. This is especially clear in Fig. 9, panel C, based on moving average means of the *∂*^18^*O* data. Considering the large standard errors in the *log AZ area mean* data it is in any case no clear indications of a very nonlinear function. Note, however, that the data from living or recent bryozoans are excluded from the analysis on the ground of extreme *∂*^18^*O* values. These values are around zero (Fig. S9 in Liow et al, 2024), and thus much lower than the last value in the *∂*^18^*O* time series in Fig. 1. With the linear function in Fig. 9, panel C, this would give log AZ area mean values around 9.3, which is much lower than the values in Fig. S2 in Liow et al. (2024). The reason for these differences may be that the individual fitness peak vs. environment function no longer is linear for extreme *∂*^18^*O* values, or that the adaptive peak vs. environment function is nonlinear owing to constraints as in Fig. 6, panels A to D. It also seems reasonable to assume that long existing species in the quite variable *∂*^18^*O* environment in general have adaptive peak vs. environment functions that are not too difficult to track, except for the last extreme environment, at least in the sense that the mean phenotypic value is fluctuating around a moving average as in Fig. 3, panel A.

The results in Fig. 9 show that the mean phenotypic values can be predicted quite well with use of a linear adaptive peak vs. environment function and moving average means of the *∂*^18^*O* data with window size 100 samples, although the predictions for samples number 5 and 6 fall outside of the error bars. With window size 120 only the prediction for sample number 8 falls outside of the error bar, but then the MSE value increases from 0.0014 to 0.0017. This indicates that each colony mean as used for computation of the population means in Fig. S6 in Liow et al. (2024) should be weighted according to the *∂*^18^*O* value for the colony. Also note that Fig. 9 gives continuous predictions, although the extensions to time periods before the first and beyond the last data point are speculative.

The prediction results using moving average means of the *∂*^18^*O* data were validated by leave-one-out cross validation, with results included in Fig. 10. Validation gave *MSE*_*CV*_ = 0.0020, which should be compared with *MSE*_*min*_ = 0.0014 for predictions. As shown in Fig. 10, the validation results are thus quite satisfactory in that all predictions of the left-out samples except two fall within the error bars.

Use of mean *∂*^18^*O* values for the different fossil formations worked less well (Fig. 8), indicating that the colony means from each formation (raw data points as shown in Fig. 2 in Liow et al., 2024) should, if possible, not all be considered as having the same *∂*^18^*O* value.

Note that although the predictions in Fig. 10 are quite good as compared with the error bars there are other possible errors that are not covered by these bars, such as fluctuations around the moving adaptive peak due to imperfect tracking, and random fluctuations of the adaptive peak itself.

It is of course an underlying assumption that the continuous mean trait predictions in Fig. 10 not only give values that can be compared to mean values from collected data, but that also mean values in time periods without a known or investigated fossil record are predicted. That remains to be verified, and it would be especially interesting if the pronounced variations around 1.5 million years ago could be reproduced.

Liow et al. (2024) report that plasticity plays a role in the real data case, although in a complex way owing to the clonal and colonial nature of the data. There is in fact a theoretical possibility that the apparently linear mean phenotype vs. environment function is caused by adaptive plasticity such that the mean phenotypic value tracks the fluctuations in sea temperature with very small errors. However, as pointed out above, a linear mean phenotype vs. environment function does not necessarily imply plasticity.

It is finally worth noting that just by looking at Fig. 1 it might be tempting to approximate the mean phenotypic values as a linear function of the moving average of the environment. The results in Fig. 10 could thus be found by an *ad hoc* least squares approach, without any consideration of the theoretical background. However, the discussion of the tracking model in Fig. 2 shows that the prediction results have a more general theoretical foundation.

## Supporting information

Raw data for simulations

MATLAB Code for all simulations

## Acknowledgement

I thank two anonymous reviewers for constructive criticism, and I thank University of South-Eastern Norway for support and funding.

## Author contribution

Rolf Ergon is the sole author of this article.

## Data availability statement

MATLAB code and data are archived on bioRxiv, doi.org/10.1101/2024.10.30.621046.

