## Supplementary material for "Adaptive peak tracking as explanation of sparse fossil data across fluctuating ancient environments": Raw data for simulations

**Raw data for MATLAB code**

for

**across fluctuating ancient environments**

Rolf Ergon

University of South-Eastern Norway

May 3, 2025

The columns in data dO18 below should be saved as x and u variables in a file O18.mat.

dO18 =

1.0e+03 *

[0000000000000 0.003230000000000

0.001000000000000 0.003230000000000

0.002000000000000 0.003180000000000

0.003000000000000 0.003290000000000

0.004000000000000 0.003300000000000

0.005000000000000 0.003260000000000

0.006000000000000 0.003330000000000

0.007000000000000 0.003370000000000

0.008000000000000 0.003420000000000

0.009000000000000 0.003380000000000

0.010000000000000 0.003520000000000

0.011000000000000 0.003600000000000

0.012000000000000 0.003920000000000

0.013000000000000 0.004060000000000

0.014000000000000 0.004280000000000

0.015000000000000 0.004490000000000

0.016000000000000 0.004750000000000

0.017000000000000 0.004880000000000

0.018000000000000 0.005020000000000

0.019000000000000 0.004960000000000

0.020000000000000 0.004990000000000

0.021000000000000 0.004910000000000

0.022000000000000 0.004880000000000

| | |
|---|---|
| 0.023000000000000 | 0.004860000000000 |
| 0.024000000000000 | 0.004810000000000 |
| 0.025000000000000 | 0.004820000000000 |
| 0.026000000000000 | 0.004670000000000 |
| 0.027000000000000 | 0.004750000000000 |
| 0.028000000000000 | 0.004750000000000 |
| 0.029000000000000 | 0.004730000000000 |
| 0.030000000000000 | 0.004620000000000 |
| 0.031000000000000 | 0.004620000000000 |
| 0.032000000000000 | 0.004600000000000 |
| 0.033000000000000 | 0.004580000000000 |
| 0.034000000000000 | 0.004590000000000 |
| 0.035000000000000 | 0.004540000000000 |
| 0.036000000000000 | 0.004450000000000 |
| 0.037000000000000 | 0.004460000000000 |
| 0.038000000000000 | 0.004410000000000 |
| 0.039000000000000 | 0.004550000000000 |
| 0.040000000000000 | 0.004550000000000 |
| 0.041000000000000 | 0.004510000000000 |
| 0.042000000000000 | 0.004500000000000 |
| 0.043000000000000 | 0.004460000000000 |
| 0.044000000000000 | 0.004480000000000 |
| 0.045000000000000 | 0.004330000000000 |
| 0.046000000000000 | 0.004360000000000 |
| 0.047000000000000 | 0.004380000000000 |
| 0.048000000000000 | 0.004450000000000 |
| 0.049000000000000 | 0.004460000000000 |
| 0.050000000000000 | 0.004340000000000 |
| 0.051000000000000 | 0.004330000000000 |
| 0.052000000000000 | 0.004310000000000 |
| 0.053000000000000 | 0.004440000000000 |
| 0.054000000000000 | 0.004380000000000 |
| 0.055000000000000 | 0.004310000000000 |
| 0.056000000000000 | 0.004350000000000 |
| 0.057000000000000 | 0.004430000000000 |
| 0.058000000000000 | 0.004490000000000 |
| 0.059000000000000 | 0.004430000000000 |
| 0.060000000000000 | 0.004600000000000 |

| | |
|---|---|
| 0.061000000000000 | 0.004510000000000 |
| 0.062000000000000 | 0.004600000000000 |
| 0.063000000000000 | 0.004530000000000 |
| 0.064000000000000 | 0.004360000000000 |
| 0.065000000000000 | 0.004480000000000 |
| 0.066000000000000 | 0.004570000000000 |
| 0.067000000000000 | 0.004440000000000 |
| 0.068000000000000 | 0.004420000000000 |
| 0.069000000000000 | 0.004470000000000 |
| 0.070000000000000 | 0.004320000000000 |
| 0.071000000000000 | 0.004220000000000 |
| 0.072000000000000 | 0.004210000000000 |
| 0.073000000000000 | 0.004230000000000 |
| 0.074000000000000 | 0.004030000000000 |
| 0.075000000000000 | 0.003950000000000 |
| 0.076000000000000 | 0.004060000000000 |
| 0.077000000000000 | 0.004080000000000 |
| 0.078000000000000 | 0.004050000000000 |
| 0.079000000000000 | 0.004050000000000 |
| 0.080000000000000 | 0.003900000000000 |
| 0.081000000000000 | 0.003820000000000 |
| 0.082000000000000 | 0.003800000000000 |
| 0.083000000000000 | 0.003830000000000 |
| 0.084000000000000 | 0.003820000000000 |
| 0.085000000000000 | 0.003950000000000 |
| 0.086000000000000 | 0.004060000000000 |
| 0.087000000000000 | 0.004180000000000 |
| 0.088000000000000 | 0.004110000000000 |
| 0.089000000000000 | 0.004080000000000 |
| 0.090000000000000 | 0.004060000000000 |
| 0.091000000000000 | 0.004030000000000 |
| 0.092000000000000 | 0.003980000000000 |
| 0.093000000000000 | 0.003900000000000 |
| 0.094000000000000 | 0.003840000000000 |
| 0.095000000000000 | 0.003770000000000 |
| 0.096000000000000 | 0.003750000000000 |
| 0.097000000000000 | 0.003830000000000 |
| 0.098000000000000 | 0.003900000000000 |

| | |
|---|---|
| 0.099000000000000 | 0.003780000000000 |
| 0.100000000000000 | 0.003810000000000 |
| 0.101000000000000 | 0.003920000000000 |
| 0.102000000000000 | 0.003860000000000 |
| 0.103000000000000 | 0.003880000000000 |
| 0.104000000000000 | 0.003920000000000 |
| 0.105000000000000 | 0.003850000000000 |
| 0.106000000000000 | 0.004000000000000 |
| 0.107000000000000 | 0.004040000000000 |
| 0.108000000000000 | 0.004110000000000 |
| 0.109000000000000 | 0.004120000000000 |
| 0.110000000000000 | 0.004040000000000 |
| 0.111000000000000 | 0.004020000000000 |
| 0.112000000000000 | 0.004030000000000 |
| 0.113000000000000 | 0.003930000000000 |
| 0.114000000000000 | 0.003810000000000 |
| 0.115000000000000 | 0.003710000000000 |
| 0.116000000000000 | 0.003580000000000 |
| 0.117000000000000 | 0.003540000000000 |
| 0.118000000000000 | 0.003440000000000 |
| 0.119000000000000 | 0.003300000000000 |
| 0.120000000000000 | 0.003270000000000 |
| 0.121000000000000 | 0.003260000000000 |
| 0.122000000000000 | 0.003180000000000 |
| 0.123000000000000 | 0.003100000000000 |
| 0.124000000000000 | 0.003270000000000 |
| 0.125000000000000 | 0.003140000000000 |
| 0.126000000000000 | 0.003160000000000 |
| 0.127000000000000 | 0.003370000000000 |
| 0.128000000000000 | 0.003710000000000 |
| 0.129000000000000 | 0.003900000000000 |
| 0.130000000000000 | 0.003670000000000 |
| 0.131000000000000 | 0.003810000000000 |
| 0.132000000000000 | 0.004200000000000 |
| 0.133000000000000 | 0.004410000000000 |
| 0.134000000000000 | 0.004700000000000 |
| 0.135000000000000 | 0.004860000000000 |
| 0.136000000000000 | 0.004820000000000 |

| | |
|---|---|
| 0.137000000000000 | 0.004800000000000 |
| 0.138000000000000 | 0.004890000000000 |
| 0.139000000000000 | 0.004870000000000 |
| 0.140000000000000 | 0.004980000000000 |
| 0.141000000000000 | 0.004810000000000 |
| 0.142000000000000 | 0.004750000000000 |
| 0.143000000000000 | 0.004780000000000 |
| 0.144000000000000 | 0.004820000000000 |
| 0.145000000000000 | 0.004740000000000 |
| 0.146000000000000 | 0.004770000000000 |
| 0.147000000000000 | 0.004820000000000 |
| 0.148000000000000 | 0.004710000000000 |
| 0.149000000000000 | 0.004750000000000 |
| 0.150000000000000 | 0.004750000000000 |
| 0.151000000000000 | 0.004660000000000 |
| 0.152000000000000 | 0.004640000000000 |
| 0.153000000000000 | 0.004620000000000 |
| 0.154000000000000 | 0.004660000000000 |
| 0.155000000000000 | 0.004510000000000 |
| 0.156000000000000 | 0.004780000000000 |
| 0.157000000000000 | 0.004740000000000 |
| 0.158000000000000 | 0.004690000000000 |
| 0.159000000000000 | 0.004660000000000 |
| 0.160000000000000 | 0.004690000000000 |
| 0.161000000000000 | 0.004650000000000 |
| 0.162000000000000 | 0.004660000000000 |
| 0.163000000000000 | 0.004700000000000 |
| 0.164000000000000 | 0.004680000000000 |
| 0.165000000000000 | 0.004680000000000 |
| 0.166000000000000 | 0.004480000000000 |
| 0.167000000000000 | 0.004410000000000 |
| 0.168000000000000 | 0.004480000000000 |
| 0.169000000000000 | 0.004500000000000 |
| 0.170000000000000 | 0.004500000000000 |
| 0.171000000000000 | 0.004530000000000 |
| 0.172000000000000 | 0.004460000000000 |
| 0.173000000000000 | 0.004380000000000 |
| 0.174000000000000 | 0.004280000000000 |

| | |
|---|---|
| 0.175000000000000 | 0.004320000000000 |
| 0.176000000000000 | 0.004390000000000 |
| 0.177000000000000 | 0.004460000000000 |
| 0.178000000000000 | 0.004440000000000 |
| 0.179000000000000 | 0.004480000000000 |
| 0.180000000000000 | 0.004390000000000 |
| 0.181000000000000 | 0.004460000000000 |
| 0.182000000000000 | 0.004480000000000 |
| 0.183000000000000 | 0.004370000000000 |
| 0.184000000000000 | 0.004400000000000 |
| 0.185000000000000 | 0.004590000000000 |
| 0.186000000000000 | 0.004470000000000 |
| 0.187000000000000 | 0.004400000000000 |
| 0.188000000000000 | 0.004460000000000 |
| 0.189000000000000 | 0.004390000000000 |
| 0.190000000000000 | 0.004280000000000 |
| 0.191000000000000 | 0.004130000000000 |
| 0.192000000000000 | 0.003760000000000 |
| 0.193000000000000 | 0.003920000000000 |
| 0.194000000000000 | 0.003840000000000 |
| 0.195000000000000 | 0.003910000000000 |
| 0.196000000000000 | 0.003820000000000 |
| 0.197000000000000 | 0.003740000000000 |
| 0.198000000000000 | 0.003820000000000 |
| 0.199000000000000 | 0.003570000000000 |
| 0.200000000000000 | 0.003530000000000 |
| 0.201000000000000 | 0.003530000000000 |
| 0.202000000000000 | 0.003630000000000 |
| 0.203000000000000 | 0.003610000000000 |
| 0.204000000000000 | 0.003780000000000 |
| 0.205000000000000 | 0.003820000000000 |
| 0.206000000000000 | 0.003770000000000 |
| 0.207000000000000 | 0.003780000000000 |
| 0.208000000000000 | 0.003710000000000 |
| 0.209000000000000 | 0.003640000000000 |
| 0.210000000000000 | 0.003570000000000 |
| 0.211000000000000 | 0.003600000000000 |
| 0.212000000000000 | 0.003610000000000 |

| | |
|---|---|
| 0.213000000000000 | 0.003650000000000 |
| 0.214000000000000 | 0.003560000000000 |
| 0.215000000000000 | 0.003540000000000 |
| 0.216000000000000 | 0.003530000000000 |
| 0.217000000000000 | 0.003480000000000 |
| 0.218000000000000 | 0.003720000000000 |
| 0.219000000000000 | 0.003940000000000 |
| 0.220000000000000 | 0.003980000000000 |
| 0.221000000000000 | 0.004230000000000 |
| 0.222000000000000 | 0.004380000000000 |
| 0.223000000000000 | 0.004440000000000 |
| 0.224000000000000 | 0.004390000000000 |
| 0.225000000000000 | 0.004280000000000 |
| 0.226000000000000 | 0.004290000000000 |
| 0.227000000000000 | 0.004190000000000 |
| 0.228000000000000 | 0.004190000000000 |
| 0.229000000000000 | 0.004020000000000 |
| 0.230000000000000 | 0.004300000000000 |
| 0.231000000000000 | 0.004220000000000 |
| 0.232000000000000 | 0.004170000000000 |
| 0.233000000000000 | 0.003970000000000 |
| 0.234000000000000 | 0.003840000000000 |
| 0.235000000000000 | 0.003710000000000 |
| 0.236000000000000 | 0.003660000000000 |
| 0.237000000000000 | 0.003470000000000 |
| 0.238000000000000 | 0.003510000000000 |
| 0.239000000000000 | 0.003440000000000 |
| 0.240000000000000 | 0.003440000000000 |
| 0.241000000000000 | 0.003540000000000 |
| 0.242000000000000 | 0.003680000000000 |
| 0.243000000000000 | 0.003780000000000 |
| 0.244000000000000 | 0.004050000000000 |
| 0.245000000000000 | 0.004170000000000 |
| 0.246000000000000 | 0.004380000000000 |
| 0.247000000000000 | 0.004310000000000 |
| 0.248000000000000 | 0.004380000000000 |
| 0.249000000000000 | 0.004360000000000 |
| 0.250000000000000 | 0.004500000000000 |

| | |
|---|---|
| 0.251000000000000 | 0.004580000000000 |
| 0.252000000000000 | 0.004630000000000 |
| 0.253000000000000 | 0.004520000000000 |
| 0.254000000000000 | 0.004570000000000 |
| 0.255000000000000 | 0.004370000000000 |
| 0.256000000000000 | 0.004430000000000 |
| 0.257000000000000 | 0.004550000000000 |
| 0.258000000000000 | 0.004510000000000 |
| 0.259000000000000 | 0.004520000000000 |
| 0.260000000000000 | 0.004460000000000 |
| 0.261000000000000 | 0.004520000000000 |
| 0.262000000000000 | 0.004450000000000 |
| 0.263000000000000 | 0.004460000000000 |
| 0.264000000000000 | 0.004420000000000 |
| 0.265000000000000 | 0.004440000000000 |
| 0.266000000000000 | 0.004500000000000 |
| 0.267000000000000 | 0.004500000000000 |
| 0.268000000000000 | 0.004330000000000 |
| 0.269000000000000 | 0.004520000000000 |
| 0.270000000000000 | 0.004510000000000 |
| 0.271000000000000 | 0.004500000000000 |
| 0.272000000000000 | 0.004440000000000 |
| 0.273000000000000 | 0.004440000000000 |
| 0.274000000000000 | 0.004380000000000 |
| 0.275000000000000 | 0.004380000000000 |
| 0.276000000000000 | 0.004300000000000 |
| 0.277000000000000 | 0.004440000000000 |
| 0.278000000000000 | 0.004310000000000 |
| 0.279000000000000 | 0.004230000000000 |
| 0.280000000000000 | 0.004140000000000 |
| 0.281000000000000 | 0.004210000000000 |
| 0.282000000000000 | 0.003980000000000 |
| 0.283000000000000 | 0.003870000000000 |
| 0.284000000000000 | 0.003860000000000 |
| 0.285000000000000 | 0.003840000000000 |
| 0.286000000000000 | 0.003820000000000 |
| 0.287000000000000 | 0.003890000000000 |
| 0.288000000000000 | 0.004020000000000 |

| | |
|---|---|
| 0.289000000000000 | 0.004060000000000 |
| 0.290000000000000 | 0.004100000000000 |
| 0.291000000000000 | 0.004110000000000 |
| 0.292000000000000 | 0.004270000000000 |
| 0.293000000000000 | 0.004220000000000 |
| 0.294000000000000 | 0.004330000000000 |
| 0.295000000000000 | 0.004310000000000 |
| 0.296000000000000 | 0.004230000000000 |
| 0.297000000000000 | 0.004300000000000 |
| 0.298000000000000 | 0.004170000000000 |
| 0.299000000000000 | 0.004230000000000 |
| 0.300000000000000 | 0.004020000000000 |
| 0.301000000000000 | 0.004060000000000 |
| 0.302000000000000 | 0.004060000000000 |
| 0.303000000000000 | 0.004000000000000 |
| 0.304000000000000 | 0.004070000000000 |
| 0.305000000000000 | 0.004070000000000 |
| 0.306000000000000 | 0.004050000000000 |
| 0.307000000000000 | 0.004000000000000 |
| 0.308000000000000 | 0.003850000000000 |
| 0.309000000000000 | 0.003860000000000 |
| 0.310000000000000 | 0.003800000000000 |
| 0.311000000000000 | 0.003700000000000 |
| 0.312000000000000 | 0.003790000000000 |
| 0.313000000000000 | 0.003690000000000 |
| 0.314000000000000 | 0.003770000000000 |
| 0.315000000000000 | 0.003620000000000 |
| 0.316000000000000 | 0.003690000000000 |
| 0.317000000000000 | 0.003890000000000 |
| 0.318000000000000 | 0.003760000000000 |
| 0.319000000000000 | 0.003580000000000 |
| 0.320000000000000 | 0.003660000000000 |
| 0.321000000000000 | 0.003530000000000 |
| 0.322000000000000 | 0.003530000000000 |
| 0.323000000000000 | 0.003470000000000 |
| 0.324000000000000 | 0.003210000000000 |
| 0.325000000000000 | 0.003310000000000 |
| 0.326000000000000 | 0.003230000000000 |

| | |
|---|---|
| 0.327000000000000 | 0.003220000000000 |
| 0.328000000000000 | 0.003230000000000 |
| 0.329000000000000 | 0.003190000000000 |
| 0.330000000000000 | 0.003300000000000 |
| 0.331000000000000 | 0.003290000000000 |
| 0.332000000000000 | 0.003390000000000 |
| 0.333000000000000 | 0.003380000000000 |
| 0.334000000000000 | 0.003510000000000 |
| 0.335000000000000 | 0.003750000000000 |
| 0.336000000000000 | 0.004080000000000 |
| 0.337000000000000 | 0.004230000000000 |
| 0.338000000000000 | 0.004170000000000 |
| 0.339000000000000 | 0.004420000000000 |
| 0.340000000000000 | 0.004600000000000 |
| 0.341000000000000 | 0.004840000000000 |
| 0.342000000000000 | 0.004830000000000 |
| 0.343000000000000 | 0.004800000000000 |
| 0.344000000000000 | 0.004760000000000 |
| 0.345000000000000 | 0.004790000000000 |
| 0.346000000000000 | 0.004680000000000 |
| 0.347000000000000 | 0.004530000000000 |
| 0.348000000000000 | 0.004490000000000 |
| 0.349000000000000 | 0.004620000000000 |
| 0.350000000000000 | 0.004670000000000 |
| 0.351000000000000 | 0.004640000000000 |
| 0.352000000000000 | 0.004470000000000 |
| 0.353000000000000 | 0.004470000000000 |
| 0.354000000000000 | 0.004400000000000 |
| 0.355000000000000 | 0.004590000000000 |
| 0.356000000000000 | 0.004460000000000 |
| 0.357000000000000 | 0.004550000000000 |
| 0.358000000000000 | 0.004580000000000 |
| 0.359000000000000 | 0.004490000000000 |
| 0.360000000000000 | 0.004540000000000 |
| 0.361000000000000 | 0.004500000000000 |
| 0.362000000000000 | 0.004500000000000 |
| 0.363000000000000 | 0.004530000000000 |
| 0.364000000000000 | 0.004360000000000 |

| | |
|---|---|
| 0.365000000000000 | 0.004460000000000 |
| 0.366000000000000 | 0.004320000000000 |
| 0.367000000000000 | 0.004370000000000 |
| 0.368000000000000 | 0.004400000000000 |
| 0.369000000000000 | 0.004380000000000 |
| 0.370000000000000 | 0.004360000000000 |
| 0.371000000000000 | 0.004270000000000 |
| 0.372000000000000 | 0.004420000000000 |
| 0.373000000000000 | 0.004290000000000 |
| 0.374000000000000 | 0.004340000000000 |
| 0.375000000000000 | 0.004200000000000 |
| 0.376000000000000 | 0.004210000000000 |
| 0.377000000000000 | 0.004200000000000 |
| 0.378000000000000 | 0.004190000000000 |
| 0.379000000000000 | 0.004170000000000 |
| 0.380000000000000 | 0.004190000000000 |
| 0.381000000000000 | 0.004150000000000 |
| 0.382000000000000 | 0.004050000000000 |
| 0.383000000000000 | 0.004120000000000 |
| 0.384000000000000 | 0.004080000000000 |
| 0.385000000000000 | 0.004140000000000 |
| 0.386000000000000 | 0.004020000000000 |
| 0.387000000000000 | 0.003990000000000 |
| 0.388000000000000 | 0.003920000000000 |
| 0.389000000000000 | 0.003890000000000 |
| 0.390000000000000 | 0.003960000000000 |
| 0.391000000000000 | 0.004000000000000 |
| 0.392000000000000 | 0.003970000000000 |
| 0.393000000000000 | 0.003860000000000 |
| 0.394000000000000 | 0.003920000000000 |
| 0.395000000000000 | 0.003760000000000 |
| 0.396000000000000 | 0.003720000000000 |
| 0.397000000000000 | 0.003470000000000 |
| 0.398000000000000 | 0.003540000000000 |
| 0.399000000000000 | 0.003480000000000 |
| 0.400000000000000 | 0.003320000000000 |
| 0.401000000000000 | 0.003200000000000 |
| 0.402000000000000 | 0.003170000000000 |

| | |
|---|---|
| 0.403000000000000 | 0.003150000000000 |
| 0.404000000000000 | 0.003200000000000 |
| 0.405000000000000 | 0.003110000000000 |
| 0.406000000000000 | 0.003190000000000 |
| 0.407000000000000 | 0.003210000000000 |
| 0.408000000000000 | 0.003170000000000 |
| 0.409000000000000 | 0.003220000000000 |
| 0.410000000000000 | 0.003150000000000 |
| 0.411000000000000 | 0.003290000000000 |
| 0.412000000000000 | 0.003310000000000 |
| 0.413000000000000 | 0.003380000000000 |
| 0.414000000000000 | 0.003410000000000 |
| 0.415000000000000 | 0.003410000000000 |
| 0.416000000000000 | 0.003510000000000 |
| 0.417000000000000 | 0.003570000000000 |
| 0.418000000000000 | 0.003750000000000 |
| 0.419000000000000 | 0.003810000000000 |
| 0.420000000000000 | 0.003770000000000 |
| 0.421000000000000 | 0.003850000000000 |
| 0.422000000000000 | 0.003940000000000 |
| 0.423000000000000 | 0.004020000000000 |
| 0.424000000000000 | 0.003920000000000 |
| 0.425000000000000 | 0.004300000000000 |
| 0.426000000000000 | 0.004500000000000 |
| 0.427000000000000 | 0.004300000000000 |
| 0.428000000000000 | 0.004720000000000 |
| 0.429000000000000 | 0.004910000000000 |
| 0.430000000000000 | 0.004980000000000 |
| 0.431000000000000 | 0.005050000000000 |
| 0.432000000000000 | 0.005050000000000 |
| 0.433000000000000 | 0.005080000000000 |
| 0.434000000000000 | 0.005080000000000 |
| 0.435000000000000 | 0.005020000000000 |
| 0.436000000000000 | 0.005070000000000 |
| 0.437000000000000 | 0.004900000000000 |
| 0.438000000000000 | 0.004790000000000 |
| 0.439000000000000 | 0.004870000000000 |
| 0.440000000000000 | 0.004760000000000 |

| | |
|---|---|
| 0.441000000000000 | 0.004950000000000 |
| 0.442000000000000 | 0.004960000000000 |
| 0.443000000000000 | 0.004910000000000 |
| 0.444000000000000 | 0.004820000000000 |
| 0.445000000000000 | 0.004910000000000 |
| 0.446000000000000 | 0.004820000000000 |
| 0.447000000000000 | 0.004510000000000 |
| 0.448000000000000 | 0.004770000000000 |
| 0.449000000000000 | 0.004730000000000 |
| 0.450000000000000 | 0.004750000000000 |
| 0.451000000000000 | 0.004580000000000 |
| 0.452000000000000 | 0.004730000000000 |
| 0.453000000000000 | 0.004560000000000 |
| 0.454000000000000 | 0.004600000000000 |
| 0.455000000000000 | 0.004410000000000 |
| 0.456000000000000 | 0.004620000000000 |
| 0.457000000000000 | 0.004630000000000 |
| 0.458000000000000 | 0.004550000000000 |
| 0.459000000000000 | 0.004630000000000 |
| 0.460000000000000 | 0.004560000000000 |
| 0.461000000000000 | 0.004590000000000 |
| 0.462000000000000 | 0.004610000000000 |
| 0.463000000000000 | 0.004610000000000 |
| 0.464000000000000 | 0.004440000000000 |
| 0.465000000000000 | 0.004570000000000 |
| 0.466000000000000 | 0.004460000000000 |
| 0.467000000000000 | 0.004440000000000 |
| 0.468000000000000 | 0.004380000000000 |
| 0.469000000000000 | 0.004510000000000 |
| 0.470000000000000 | 0.004330000000000 |
| 0.471000000000000 | 0.004250000000000 |
| 0.472000000000000 | 0.004320000000000 |
| 0.473000000000000 | 0.004360000000000 |
| 0.474000000000000 | 0.004390000000000 |
| 0.475000000000000 | 0.004350000000000 |
| 0.476000000000000 | 0.004300000000000 |
| 0.477000000000000 | 0.004270000000000 |
| 0.478000000000000 | 0.004220000000000 |

| | |
|---|---|
| 0.479000000000000 | 0.004120000000000 |
| 0.480000000000000 | 0.004120000000000 |
| 0.481000000000000 | 0.004170000000000 |
| 0.482000000000000 | 0.004010000000000 |
| 0.483000000000000 | 0.003970000000000 |
| 0.484000000000000 | 0.003790000000000 |
| 0.485000000000000 | 0.003810000000000 |
| 0.486000000000000 | 0.003760000000000 |
| 0.487000000000000 | 0.003720000000000 |
| 0.488000000000000 | 0.003670000000000 |
| 0.489000000000000 | 0.003600000000000 |
| 0.490000000000000 | 0.003540000000000 |
| 0.491000000000000 | 0.003470000000000 |
| 0.492000000000000 | 0.003630000000000 |
| 0.493000000000000 | 0.003720000000000 |
| 0.494000000000000 | 0.003830000000000 |
| 0.495000000000000 | 0.003830000000000 |
| 0.496000000000000 | 0.003830000000000 |
| 0.497000000000000 | 0.003680000000000 |
| 0.498000000000000 | 0.003820000000000 |
| 0.499000000000000 | 0.003750000000000 |
| 0.500000000000000 | 0.003840000000000 |
| 0.501000000000000 | 0.003780000000000 |
| 0.502000000000000 | 0.003750000000000 |
| 0.503000000000000 | 0.003810000000000 |
| 0.504000000000000 | 0.003910000000000 |
| 0.505000000000000 | 0.003860000000000 |
| 0.506000000000000 | 0.004010000000000 |
| 0.507000000000000 | 0.003930000000000 |
| 0.508000000000000 | 0.004160000000000 |
| 0.509000000000000 | 0.004060000000000 |
| 0.510000000000000 | 0.004140000000000 |
| 0.511000000000000 | 0.004060000000000 |
| 0.512000000000000 | 0.004220000000000 |
| 0.513000000000000 | 0.004250000000000 |
| 0.514000000000000 | 0.004100000000000 |
| 0.515000000000000 | 0.004100000000000 |
| 0.516000000000000 | 0.004010000000000 |

| | |
|---|---|
| 0.517000000000000 | 0.003960000000000 |
| 0.518000000000000 | 0.004070000000000 |
| 0.519000000000000 | 0.003960000000000 |
| 0.520000000000000 | 0.003950000000000 |
| 0.521000000000000 | 0.003910000000000 |
| 0.522000000000000 | 0.003880000000000 |
| 0.523000000000000 | 0.003930000000000 |
| 0.524000000000000 | 0.003830000000000 |
| 0.525000000000000 | 0.003920000000000 |
| 0.526000000000000 | 0.003970000000000 |
| 0.527000000000000 | 0.003960000000000 |
| 0.528000000000000 | 0.003920000000000 |
| 0.529000000000000 | 0.003990000000000 |
| 0.530000000000000 | 0.003970000000000 |
| 0.531000000000000 | 0.003920000000000 |
| 0.532000000000000 | 0.004010000000000 |
| 0.533000000000000 | 0.004110000000000 |
| 0.534000000000000 | 0.004270000000000 |
| 0.535000000000000 | 0.004290000000000 |
| 0.536000000000000 | 0.004550000000000 |
| 0.537000000000000 | 0.004430000000000 |
| 0.538000000000000 | 0.004540000000000 |
| 0.539000000000000 | 0.004480000000000 |
| 0.540000000000000 | 0.004530000000000 |
| 0.541000000000000 | 0.004500000000000 |
| 0.542000000000000 | 0.004450000000000 |
| 0.543000000000000 | 0.004450000000000 |
| 0.544000000000000 | 0.004460000000000 |
| 0.545000000000000 | 0.004430000000000 |
| 0.546000000000000 | 0.004510000000000 |
| 0.547000000000000 | 0.004530000000000 |
| 0.548000000000000 | 0.004550000000000 |
| 0.549000000000000 | 0.004530000000000 |
| 0.550000000000000 | 0.004500000000000 |
| 0.551000000000000 | 0.004450000000000 |
| 0.552000000000000 | 0.004520000000000 |
| 0.553000000000000 | 0.004460000000000 |
| 0.554000000000000 | 0.004290000000000 |

| | |
|---|---|
| 0.555000000000000 | 0.004250000000000 |
| 0.556000000000000 | 0.004390000000000 |
| 0.557000000000000 | 0.004350000000000 |
| 0.558000000000000 | 0.004250000000000 |
| 0.559000000000000 | 0.004170000000000 |
| 0.560000000000000 | 0.004140000000000 |
| 0.561000000000000 | 0.004210000000000 |
| 0.562000000000000 | 0.004140000000000 |
| 0.563000000000000 | 0.004170000000000 |
| 0.564000000000000 | 0.004010000000000 |
| 0.565000000000000 | 0.003960000000000 |
| 0.566000000000000 | 0.003860000000000 |
| 0.567000000000000 | 0.003940000000000 |
| 0.568000000000000 | 0.004000000000000 |
| 0.569000000000000 | 0.003970000000000 |
| 0.570000000000000 | 0.003910000000000 |
| 0.571000000000000 | 0.003890000000000 |
| 0.572000000000000 | 0.003590000000000 |
| 0.573000000000000 | 0.003650000000000 |
| 0.574000000000000 | 0.003730000000000 |
| 0.575000000000000 | 0.003390000000000 |
| 0.576000000000000 | 0.003460000000000 |
| 0.577000000000000 | 0.003480000000000 |
| 0.578000000000000 | 0.003500000000000 |
| 0.579000000000000 | 0.003570000000000 |
| 0.580000000000000 | 0.003690000000000 |
| 0.581000000000000 | 0.004000000000000 |
| 0.582000000000000 | 0.004130000000000 |
| 0.583000000000000 | 0.004290000000000 |
| 0.584000000000000 | 0.004310000000000 |
| 0.585000000000000 | 0.004330000000000 |
| 0.586000000000000 | 0.004180000000000 |
| 0.587000000000000 | 0.004110000000000 |
| 0.588000000000000 | 0.004190000000000 |
| 0.589000000000000 | 0.004070000000000 |
| 0.590000000000000 | 0.004090000000000 |
| 0.591000000000000 | 0.004070000000000 |
| 0.592000000000000 | 0.004020000000000 |

| | |
|---|---|
| 0.593000000000000 | 0.003990000000000 |
| 0.594000000000000 | 0.003900000000000 |
| 0.595000000000000 | 0.003880000000000 |
| 0.596000000000000 | 0.003890000000000 |
| 0.597000000000000 | 0.003800000000000 |
| 0.598000000000000 | 0.003990000000000 |
| 0.599000000000000 | 0.004020000000000 |
| 0.600000000000000 | 0.004070000000000 |
| 0.602000000000000 | 0.003970000000000 |
| 0.604000000000000 | 0.003770000000000 |
| 0.606000000000000 | 0.003800000000000 |
| 0.608000000000000 | 0.003800000000000 |
| 0.610000000000000 | 0.003490000000000 |
| 0.612000000000000 | 0.003520000000000 |
| 0.614000000000000 | 0.003530000000000 |
| 0.616000000000000 | 0.003660000000000 |
| 0.618000000000000 | 0.003810000000000 |
| 0.620000000000000 | 0.004090000000000 |
| 0.622000000000000 | 0.004310000000000 |
| 0.624000000000000 | 0.004630000000000 |
| 0.626000000000000 | 0.004900000000000 |
| 0.628000000000000 | 0.005010000000000 |
| 0.630000000000000 | 0.005080000000000 |
| 0.632000000000000 | 0.005050000000000 |
| 0.634000000000000 | 0.004930000000000 |
| 0.636000000000000 | 0.005000000000000 |
| 0.638000000000000 | 0.004800000000000 |
| 0.640000000000000 | 0.005010000000000 |
| 0.642000000000000 | 0.004750000000000 |
| 0.644000000000000 | 0.004680000000000 |
| 0.646000000000000 | 0.004720000000000 |
| 0.648000000000000 | 0.004750000000000 |
| 0.650000000000000 | 0.004710000000000 |
| 0.652000000000000 | 0.004790000000000 |
| 0.654000000000000 | 0.004770000000000 |
| 0.656000000000000 | 0.004570000000000 |
| 0.658000000000000 | 0.004580000000000 |
| 0.660000000000000 | 0.004640000000000 |

| | |
|---|---|
| 0.662000000000000 | 0.004800000000000 |
| 0.664000000000000 | 0.004580000000000 |
| 0.666000000000000 | 0.004500000000000 |
| 0.668000000000000 | 0.004650000000000 |
| 0.670000000000000 | 0.004420000000000 |
| 0.672000000000000 | 0.004520000000000 |
| 0.674000000000000 | 0.004460000000000 |
| 0.676000000000000 | 0.004400000000000 |
| 0.678000000000000 | 0.004260000000000 |
| 0.680000000000000 | 0.004140000000000 |
| 0.682000000000000 | 0.004150000000000 |
| 0.684000000000000 | 0.004030000000000 |
| 0.686000000000000 | 0.004130000000000 |
| 0.688000000000000 | 0.003950000000000 |
| 0.690000000000000 | 0.003700000000000 |
| 0.692000000000000 | 0.003710000000000 |
| 0.694000000000000 | 0.003640000000000 |
| 0.696000000000000 | 0.003500000000000 |
| 0.698000000000000 | 0.003570000000000 |
| 0.700000000000000 | 0.003650000000000 |
| 0.702000000000000 | 0.003760000000000 |
| 0.704000000000000 | 0.003840000000000 |
| 0.706000000000000 | 0.003930000000000 |
| 0.708000000000000 | 0.004030000000000 |
| 0.710000000000000 | 0.003980000000000 |
| 0.712000000000000 | 0.004060000000000 |
| 0.714000000000000 | 0.004290000000000 |
| 0.716000000000000 | 0.004610000000000 |
| 0.718000000000000 | 0.004750000000000 |
| 0.720000000000000 | 0.004660000000000 |
| 0.722000000000000 | 0.004650000000000 |
| 0.724000000000000 | 0.004440000000000 |
| 0.726000000000000 | 0.004320000000000 |
| 0.728000000000000 | 0.004160000000000 |
| 0.730000000000000 | 0.004160000000000 |
| 0.732000000000000 | 0.004020000000000 |
| 0.734000000000000 | 0.004150000000000 |
| 0.736000000000000 | 0.004190000000000 |

| | |
|---|---|
| 0.738000000000000 | 0.004210000000000 |
| 0.740000000000000 | 0.004110000000000 |
| 0.742000000000000 | 0.004170000000000 |
| 0.744000000000000 | 0.004480000000000 |
| 0.746000000000000 | 0.004670000000000 |
| 0.748000000000000 | 0.004590000000000 |
| 0.750000000000000 | 0.004600000000000 |
| 0.752000000000000 | 0.004480000000000 |
| 0.754000000000000 | 0.004550000000000 |
| 0.756000000000000 | 0.004590000000000 |
| 0.758000000000000 | 0.004260000000000 |
| 0.760000000000000 | 0.004330000000000 |
| 0.762000000000000 | 0.004110000000000 |
| 0.764000000000000 | 0.004280000000000 |
| 0.766000000000000 | 0.004110000000000 |
| 0.768000000000000 | 0.004070000000000 |
| 0.770000000000000 | 0.003990000000000 |
| 0.772000000000000 | 0.003870000000000 |
| 0.774000000000000 | 0.003760000000000 |
| 0.776000000000000 | 0.003640000000000 |
| 0.778000000000000 | 0.003560000000000 |
| 0.780000000000000 | 0.003480000000000 |
| 0.782000000000000 | 0.003540000000000 |
| 0.784000000000000 | 0.003550000000000 |
| 0.786000000000000 | 0.003680000000000 |
| 0.788000000000000 | 0.003700000000000 |
| 0.790000000000000 | 0.004160000000000 |
| 0.792000000000000 | 0.004270000000000 |
| 0.794000000000000 | 0.004740000000000 |
| 0.796000000000000 | 0.004670000000000 |
| 0.798000000000000 | 0.004680000000000 |
| 0.800000000000000 | 0.004680000000000 |
| 0.802000000000000 | 0.004730000000000 |
| 0.804000000000000 | 0.004550000000000 |
| 0.806000000000000 | 0.004530000000000 |
| 0.808000000000000 | 0.004490000000000 |
| 0.810000000000000 | 0.004310000000000 |
| 0.812000000000000 | 0.004380000000000 |

| | |
|---|---|
| 0.814000000000000 | 0.004120000000000 |
| 0.816000000000000 | 0.004080000000000 |
| 0.818000000000000 | 0.003900000000000 |
| 0.820000000000000 | 0.003960000000000 |
| 0.822000000000000 | 0.003900000000000 |
| 0.824000000000000 | 0.003920000000000 |
| 0.826000000000000 | 0.004110000000000 |
| 0.828000000000000 | 0.003950000000000 |
| 0.830000000000000 | 0.004040000000000 |
| 0.832000000000000 | 0.004070000000000 |
| 0.834000000000000 | 0.003950000000000 |
| 0.836000000000000 | 0.003820000000000 |
| 0.838000000000000 | 0.003990000000000 |
| 0.840000000000000 | 0.004020000000000 |
| 0.842000000000000 | 0.003810000000000 |
| 0.844000000000000 | 0.003670000000000 |
| 0.846000000000000 | 0.003580000000000 |
| 0.848000000000000 | 0.003630000000000 |
| 0.850000000000000 | 0.003500000000000 |
| 0.852000000000000 | 0.003650000000000 |
| 0.854000000000000 | 0.003630000000000 |
| 0.856000000000000 | 0.003460000000000 |
| 0.858000000000000 | 0.003420000000000 |
| 0.860000000000000 | 0.003450000000000 |
| 0.862000000000000 | 0.003680000000000 |
| 0.864000000000000 | 0.003580000000000 |
| 0.866000000000000 | 0.004060000000000 |
| 0.868000000000000 | 0.004320000000000 |
| 0.870000000000000 | 0.004510000000000 |
| 0.872000000000000 | 0.004670000000000 |
| 0.874000000000000 | 0.004680000000000 |
| 0.876000000000000 | 0.004690000000000 |
| 0.878000000000000 | 0.004590000000000 |
| 0.880000000000000 | 0.004680000000000 |
| 0.882000000000000 | 0.004420000000000 |
| 0.884000000000000 | 0.004470000000000 |
| 0.886000000000000 | 0.004470000000000 |
| 0.888000000000000 | 0.004450000000000 |

| | |
|---|---|
| 0.890000000000000 | 0.004390000000000 |
| 0.892000000000000 | 0.004320000000000 |
| 0.894000000000000 | 0.004410000000000 |
| 0.896000000000000 | 0.004400000000000 |
| 0.898000000000000 | 0.004330000000000 |
| 0.900000000000000 | 0.004290000000000 |
| 0.902000000000000 | 0.004170000000000 |
| 0.904000000000000 | 0.004140000000000 |
| 0.906000000000000 | 0.004120000000000 |
| 0.908000000000000 | 0.003990000000000 |
| 0.910000000000000 | 0.003990000000000 |
| 0.912000000000000 | 0.004090000000000 |
| 0.914000000000000 | 0.004030000000000 |
| 0.916000000000000 | 0.004060000000000 |
| 0.918000000000000 | 0.004370000000000 |
| 0.920000000000000 | 0.004510000000000 |
| 0.922000000000000 | 0.004550000000000 |
| 0.924000000000000 | 0.004430000000000 |
| 0.926000000000000 | 0.004380000000000 |
| 0.928000000000000 | 0.004260000000000 |
| 0.930000000000000 | 0.004100000000000 |
| 0.932000000000000 | 0.004070000000000 |
| 0.934000000000000 | 0.003980000000000 |
| 0.936000000000000 | 0.004020000000000 |
| 0.938000000000000 | 0.003860000000000 |
| 0.940000000000000 | 0.003890000000000 |
| 0.942000000000000 | 0.003690000000000 |
| 0.944000000000000 | 0.003570000000000 |
| 0.946000000000000 | 0.003630000000000 |
| 0.948000000000000 | 0.003480000000000 |
| 0.950000000000000 | 0.003440000000000 |
| 0.952000000000000 | 0.003300000000000 |
| 0.954000000000000 | 0.003390000000000 |
| 0.956000000000000 | 0.003330000000000 |
| 0.958000000000000 | 0.003730000000000 |
| 0.960000000000000 | 0.004160000000000 |
| 0.962000000000000 | 0.004420000000000 |
| 0.964000000000000 | 0.004580000000000 |

| | |
|---|---|
| 0.966000000000000 | 0.004300000000000 |
| 0.968000000000000 | 0.004210000000000 |
| 0.970000000000000 | 0.004090000000000 |
| 0.972000000000000 | 0.003980000000000 |
| 0.974000000000000 | 0.003870000000000 |
| 0.976000000000000 | 0.003770000000000 |
| 0.978000000000000 | 0.003720000000000 |
| 0.980000000000000 | 0.003760000000000 |
| 0.982000000000000 | 0.003990000000000 |
| 0.984000000000000 | 0.004150000000000 |
| 0.986000000000000 | 0.004240000000000 |
| 0.988000000000000 | 0.004210000000000 |
| 0.990000000000000 | 0.004110000000000 |
| 0.992000000000000 | 0.004060000000000 |
| 0.994000000000000 | 0.003940000000000 |
| 0.996000000000000 | 0.003840000000000 |
| 0.998000000000000 | 0.003840000000000 |
| 1.000000000000000 | 0.003940000000000 |
| 1.002000000000000 | 0.004150000000000 |
| 1.004000000000000 | 0.004280000000000 |
| 1.006000000000000 | 0.004220000000000 |
| 1.008000000000000 | 0.004220000000000 |
| 1.010000000000000 | 0.004120000000000 |
| 1.012000000000000 | 0.004130000000000 |
| 1.014000000000000 | 0.003990000000000 |
| 1.016000000000000 | 0.003730000000000 |
| 1.018000000000000 | 0.003780000000000 |
| 1.020000000000000 | 0.003830000000000 |
| 1.022000000000000 | 0.003570000000000 |
| 1.024000000000000 | 0.003640000000000 |
| 1.026000000000000 | 0.003870000000000 |
| 1.028000000000000 | 0.003840000000000 |
| 1.030000000000000 | 0.003950000000000 |
| 1.032000000000000 | 0.004080000000000 |
| 1.034000000000000 | 0.004360000000000 |
| 1.036000000000000 | 0.004330000000000 |
| 1.038000000000000 | 0.004540000000000 |
| 1.040000000000000 | 0.004330000000000 |

| | |
|---|---|
| 1.042000000000000 | 0.004230000000000 |
| 1.044000000000000 | 0.004380000000000 |
| 1.046000000000000 | 0.004220000000000 |
| 1.048000000000000 | 0.004290000000000 |
| 1.050000000000000 | 0.004180000000000 |
| 1.052000000000000 | 0.004270000000000 |
| 1.054000000000000 | 0.004250000000000 |
| 1.056000000000000 | 0.004120000000000 |
| 1.058000000000000 | 0.004230000000000 |
| 1.060000000000000 | 0.004080000000000 |
| 1.062000000000000 | 0.003940000000000 |
| 1.064000000000000 | 0.003660000000000 |
| 1.066000000000000 | 0.003480000000000 |
| 1.068000000000000 | 0.003330000000000 |
| 1.070000000000000 | 0.003340000000000 |
| 1.072000000000000 | 0.003210000000000 |
| 1.074000000000000 | 0.003420000000000 |
| 1.076000000000000 | 0.003480000000000 |
| 1.078000000000000 | 0.003540000000000 |
| 1.080000000000000 | 0.003610000000000 |
| 1.082000000000000 | 0.003810000000000 |
| 1.084000000000000 | 0.003700000000000 |
| 1.086000000000000 | 0.003890000000000 |
| 1.088000000000000 | 0.003910000000000 |
| 1.090000000000000 | 0.003950000000000 |
| 1.092000000000000 | 0.004010000000000 |
| 1.094000000000000 | 0.004000000000000 |
| 1.096000000000000 | 0.004030000000000 |
| 1.098000000000000 | 0.004290000000000 |
| 1.100000000000000 | 0.004080000000000 |
| 1.102000000000000 | 0.004030000000000 |
| 1.104000000000000 | 0.003930000000000 |
| 1.106000000000000 | 0.003830000000000 |
| 1.108000000000000 | 0.003670000000000 |
| 1.110000000000000 | 0.003850000000000 |
| 1.112000000000000 | 0.003840000000000 |
| 1.114000000000000 | 0.004120000000000 |
| 1.116000000000000 | 0.004140000000000 |

| | |
|---|---|
| 1.118000000000000 | 0.004200000000000 |
| 1.120000000000000 | 0.004090000000000 |
| 1.122000000000000 | 0.004470000000000 |
| 1.124000000000000 | 0.004340000000000 |
| 1.126000000000000 | 0.004500000000000 |
| 1.128000000000000 | 0.004380000000000 |
| 1.130000000000000 | 0.004320000000000 |
| 1.132000000000000 | 0.004050000000000 |
| 1.134000000000000 | 0.004220000000000 |
| 1.136000000000000 | 0.004240000000000 |
| 1.138000000000000 | 0.004020000000000 |
| 1.140000000000000 | 0.004020000000000 |
| 1.142000000000000 | 0.003970000000000 |
| 1.144000000000000 | 0.003990000000000 |
| 1.146000000000000 | 0.003860000000000 |
| 1.148000000000000 | 0.004020000000000 |
| 1.150000000000000 | 0.003820000000000 |
| 1.152000000000000 | 0.003750000000000 |
| 1.154000000000000 | 0.003760000000000 |
| 1.156000000000000 | 0.003790000000000 |
| 1.158000000000000 | 0.003680000000000 |
| 1.160000000000000 | 0.003600000000000 |
| 1.162000000000000 | 0.003600000000000 |
| 1.164000000000000 | 0.003660000000000 |
| 1.166000000000000 | 0.003530000000000 |
| 1.168000000000000 | 0.003560000000000 |
| 1.170000000000000 | 0.003600000000000 |
| 1.172000000000000 | 0.003730000000000 |
| 1.174000000000000 | 0.003760000000000 |
| 1.176000000000000 | 0.003740000000000 |
| 1.178000000000000 | 0.003800000000000 |
| 1.180000000000000 | 0.003600000000000 |
| 1.182000000000000 | 0.003840000000000 |
| 1.184000000000000 | 0.003650000000000 |
| 1.186000000000000 | 0.003770000000000 |
| 1.188000000000000 | 0.003670000000000 |
| 1.190000000000000 | 0.004050000000000 |
| 1.192000000000000 | 0.004140000000000 |

| | |
|---|---|
| 1.194000000000000 | 0.004270000000000 |
| 1.196000000000000 | 0.004280000000000 |
| 1.198000000000000 | 0.004430000000000 |
| 1.200000000000000 | 0.004420000000000 |
| 1.202000000000000 | 0.004200000000000 |
| 1.204000000000000 | 0.004340000000000 |
| 1.206000000000000 | 0.004160000000000 |
| 1.208000000000000 | 0.004320000000000 |
| 1.210000000000000 | 0.004200000000000 |
| 1.212000000000000 | 0.004080000000000 |
| 1.214000000000000 | 0.004100000000000 |
| 1.216000000000000 | 0.003820000000000 |
| 1.218000000000000 | 0.003730000000000 |
| 1.220000000000000 | 0.003970000000000 |
| 1.222000000000000 | 0.003940000000000 |
| 1.224000000000000 | 0.003740000000000 |
| 1.226000000000000 | 0.003630000000000 |
| 1.228000000000000 | 0.003550000000000 |
| 1.230000000000000 | 0.003420000000000 |
| 1.232000000000000 | 0.003410000000000 |
| 1.234000000000000 | 0.003340000000000 |
| 1.236000000000000 | 0.003380000000000 |
| 1.238000000000000 | 0.003350000000000 |
| 1.240000000000000 | 0.003330000000000 |
| 1.242000000000000 | 0.003470000000000 |
| 1.244000000000000 | 0.003850000000000 |
| 1.246000000000000 | 0.004100000000000 |
| 1.248000000000000 | 0.004360000000000 |
| 1.250000000000000 | 0.004310000000000 |
| 1.252000000000000 | 0.004250000000000 |
| 1.254000000000000 | 0.004270000000000 |
| 1.256000000000000 | 0.004110000000000 |
| 1.258000000000000 | 0.003900000000000 |
| 1.260000000000000 | 0.003870000000000 |
| 1.262000000000000 | 0.004070000000000 |
| 1.264000000000000 | 0.003960000000000 |
| 1.266000000000000 | 0.003910000000000 |
| 1.268000000000000 | 0.003910000000000 |

| | |
|---|---|
| 1.270000000000000 | 0.003840000000000 |
| 1.272000000000000 | 0.003810000000000 |
| 1.274000000000000 | 0.003840000000000 |
| 1.276000000000000 | 0.003710000000000 |
| 1.278000000000000 | 0.003670000000000 |
| 1.280000000000000 | 0.003560000000000 |
| 1.282000000000000 | 0.003650000000000 |
| 1.284000000000000 | 0.003840000000000 |
| 1.286000000000000 | 0.003970000000000 |
| 1.288000000000000 | 0.004260000000000 |
| 1.290000000000000 | 0.004350000000000 |
| 1.292000000000000 | 0.004140000000000 |
| 1.294000000000000 | 0.003940000000000 |
| 1.296000000000000 | 0.004200000000000 |
| 1.298000000000000 | 0.004150000000000 |
| 1.300000000000000 | 0.004070000000000 |
| 1.302000000000000 | 0.004060000000000 |
| 1.304000000000000 | 0.003970000000000 |
| 1.306000000000000 | 0.003900000000000 |
| 1.308000000000000 | 0.003800000000000 |
| 1.310000000000000 | 0.003650000000000 |
| 1.312000000000000 | 0.003690000000000 |
| 1.314000000000000 | 0.003710000000000 |
| 1.316000000000000 | 0.003590000000000 |
| 1.318000000000000 | 0.003730000000000 |
| 1.320000000000000 | 0.003950000000000 |
| 1.322000000000000 | 0.003940000000000 |
| 1.324000000000000 | 0.003990000000000 |
| 1.326000000000000 | 0.004000000000000 |
| 1.328000000000000 | 0.004090000000000 |
| 1.330000000000000 | 0.004060000000000 |
| 1.332000000000000 | 0.004020000000000 |
| 1.334000000000000 | 0.004190000000000 |
| 1.336000000000000 | 0.004190000000000 |
| 1.338000000000000 | 0.004100000000000 |
| 1.340000000000000 | 0.004190000000000 |
| 1.342000000000000 | 0.004070000000000 |
| 1.344000000000000 | 0.003880000000000 |

| | |
|---|---|
| 1.346000000000000 | 0.003860000000000 |
| 1.348000000000000 | 0.003670000000000 |
| 1.350000000000000 | 0.003590000000000 |
| 1.352000000000000 | 0.003490000000000 |
| 1.354000000000000 | 0.003410000000000 |
| 1.356000000000000 | 0.003500000000000 |
| 1.358000000000000 | 0.003550000000000 |
| 1.360000000000000 | 0.003630000000000 |
| 1.362000000000000 | 0.003800000000000 |
| 1.364000000000000 | 0.003980000000000 |
| 1.366000000000000 | 0.003990000000000 |
| 1.368000000000000 | 0.004200000000000 |
| 1.370000000000000 | 0.004040000000000 |
| 1.372000000000000 | 0.004260000000000 |
| 1.374000000000000 | 0.004210000000000 |
| 1.376000000000000 | 0.004080000000000 |
| 1.378000000000000 | 0.004200000000000 |
| 1.380000000000000 | 0.004050000000000 |
| 1.382000000000000 | 0.003960000000000 |
| 1.384000000000000 | 0.003820000000000 |
| 1.386000000000000 | 0.003720000000000 |
| 1.388000000000000 | 0.003610000000000 |
| 1.390000000000000 | 0.003730000000000 |
| 1.392000000000000 | 0.003590000000000 |
| 1.394000000000000 | 0.003700000000000 |
| 1.396000000000000 | 0.003540000000000 |
| 1.398000000000000 | 0.003490000000000 |
| 1.400000000000000 | 0.003560000000000 |
| 1.402000000000000 | 0.003630000000000 |
| 1.404000000000000 | 0.003740000000000 |
| 1.406000000000000 | 0.003970000000000 |
| 1.408000000000000 | 0.003970000000000 |
| 1.410000000000000 | 0.004200000000000 |
| 1.412000000000000 | 0.004400000000000 |
| 1.414000000000000 | 0.004220000000000 |
| 1.416000000000000 | 0.003960000000000 |
| 1.418000000000000 | 0.003840000000000 |
| 1.420000000000000 | 0.003960000000000 |

| | |
|---|---|
| 1.422000000000000 | 0.003830000000000 |
| 1.424000000000000 | 0.003720000000000 |
| 1.426000000000000 | 0.003720000000000 |
| 1.428000000000000 | 0.003610000000000 |
| 1.430000000000000 | 0.003520000000000 |
| 1.432000000000000 | 0.003380000000000 |
| 1.434000000000000 | 0.003380000000000 |
| 1.436000000000000 | 0.003240000000000 |
| 1.438000000000000 | 0.003240000000000 |
| 1.440000000000000 | 0.003280000000000 |
| 1.442000000000000 | 0.003340000000000 |
| 1.444000000000000 | 0.003330000000000 |
| 1.446000000000000 | 0.003360000000000 |
| 1.448000000000000 | 0.003350000000000 |
| 1.450000000000000 | 0.003490000000000 |
| 1.452000000000000 | 0.003800000000000 |
| 1.454000000000000 | 0.004200000000000 |
| 1.456000000000000 | 0.004310000000000 |
| 1.458000000000000 | 0.004240000000000 |
| 1.460000000000000 | 0.004100000000000 |
| 1.462000000000000 | 0.004070000000000 |
| 1.464000000000000 | 0.004220000000000 |
| 1.466000000000000 | 0.003990000000000 |
| 1.468000000000000 | 0.003870000000000 |
| 1.470000000000000 | 0.003620000000000 |
| 1.472000000000000 | 0.003450000000000 |
| 1.474000000000000 | 0.003380000000000 |
| 1.476000000000000 | 0.003310000000000 |
| 1.478000000000000 | 0.003420000000000 |
| 1.480000000000000 | 0.003390000000000 |
| 1.482000000000000 | 0.003440000000000 |
| 1.484000000000000 | 0.003570000000000 |
| 1.486000000000000 | 0.003420000000000 |
| 1.488000000000000 | 0.003420000000000 |
| 1.490000000000000 | 0.003510000000000 |
| 1.492000000000000 | 0.003770000000000 |
| 1.494000000000000 | 0.004050000000000 |
| 1.496000000000000 | 0.004360000000000 |

| | |
|---|---|
| 1.498000000000000 | 0.004240000000000 |
| 1.500000000000000 | 0.004210000000000 |
| 1.502500000000000 | 0.004290000000000 |
| 1.505000000000000 | 0.004160000000000 |
| 1.507500000000000 | 0.004150000000000 |
| 1.510000000000000 | 0.004020000000000 |
| 1.512500000000000 | 0.003740000000000 |
| 1.515000000000000 | 0.003690000000000 |
| 1.517500000000000 | 0.003790000000000 |
| 1.520000000000000 | 0.003640000000000 |
| 1.522500000000000 | 0.003670000000000 |
| 1.525000000000000 | 0.003700000000000 |
| 1.527500000000000 | 0.003890000000000 |
| 1.530000000000000 | 0.003960000000000 |
| 1.532500000000000 | 0.004200000000000 |
| 1.535000000000000 | 0.004410000000000 |
| 1.537500000000000 | 0.004350000000000 |
| 1.540000000000000 | 0.004380000000000 |
| 1.542500000000000 | 0.004300000000000 |
| 1.545000000000000 | 0.004070000000000 |
| 1.547500000000000 | 0.003970000000000 |
| 1.550000000000000 | 0.003820000000000 |
| 1.552500000000000 | 0.003680000000000 |
| 1.555000000000000 | 0.003720000000000 |
| 1.557500000000000 | 0.003790000000000 |
| 1.560000000000000 | 0.003760000000000 |
| 1.562500000000000 | 0.003580000000000 |
| 1.565000000000000 | 0.003560000000000 |
| 1.567500000000000 | 0.003730000000000 |
| 1.570000000000000 | 0.003850000000000 |
| 1.572500000000000 | 0.004170000000000 |
| 1.575000000000000 | 0.004240000000000 |
| 1.577500000000000 | 0.004180000000000 |
| 1.580000000000000 | 0.003990000000000 |
| 1.582500000000000 | 0.003990000000000 |
| 1.585000000000000 | 0.003800000000000 |
| 1.587500000000000 | 0.003660000000000 |
| 1.590000000000000 | 0.003720000000000 |

| | |
|---|---|
| 1.592500000000000 | 0.003860000000000 |
| 1.595000000000000 | 0.003710000000000 |
| 1.597500000000000 | 0.003610000000000 |
| 1.600000000000000 | 0.003530000000000 |
| 1.602500000000000 | 0.003370000000000 |
| 1.605000000000000 | 0.003550000000000 |
| 1.607500000000000 | 0.003660000000000 |
| 1.610000000000000 | 0.003660000000000 |
| 1.612500000000000 | 0.003880000000000 |
| 1.615000000000000 | 0.003840000000000 |
| 1.617500000000000 | 0.003900000000000 |
| 1.620000000000000 | 0.003650000000000 |
| 1.622500000000000 | 0.003680000000000 |
| 1.625000000000000 | 0.003760000000000 |
| 1.627500000000000 | 0.004010000000000 |
| 1.630000000000000 | 0.003550000000000 |
| 1.632500000000000 | 0.003570000000000 |
| 1.635000000000000 | 0.003560000000000 |
| 1.637500000000000 | 0.003600000000000 |
| 1.640000000000000 | 0.003680000000000 |
| 1.642500000000000 | 0.003880000000000 |
| 1.645000000000000 | 0.004050000000000 |
| 1.647500000000000 | 0.004170000000000 |
| 1.650000000000000 | 0.004180000000000 |
| 1.652500000000000 | 0.004250000000000 |
| 1.655000000000000 | 0.004280000000000 |
| 1.657500000000000 | 0.004110000000000 |
| 1.660000000000000 | 0.004130000000000 |
| 1.662500000000000 | 0.004080000000000 |
| 1.665000000000000 | 0.004130000000000 |
| 1.667500000000000 | 0.003910000000000 |
| 1.670000000000000 | 0.003880000000000 |
| 1.672500000000000 | 0.003860000000000 |
| 1.675000000000000 | 0.003550000000000 |
| 1.677500000000000 | 0.003720000000000 |
| 1.680000000000000 | 0.003650000000000 |
| 1.682500000000000 | 0.003570000000000 |
| 1.685000000000000 | 0.003590000000000 |

| | |
|---|---|
| 1.687500000000000 | 0.003740000000000 |
| 1.690000000000000 | 0.003720000000000 |
| 1.692500000000000 | 0.003750000000000 |
| 1.695000000000000 | 0.003820000000000 |
| 1.697500000000000 | 0.003770000000000 |
| 1.700000000000000 | 0.004090000000000 |
| 1.702500000000000 | 0.004150000000000 |
| 1.705000000000000 | 0.004120000000000 |
| 1.707500000000000 | 0.004170000000000 |
| 1.710000000000000 | 0.003930000000000 |
| 1.712500000000000 | 0.003860000000000 |
| 1.715000000000000 | 0.003830000000000 |
| 1.717500000000000 | 0.003570000000000 |
| 1.720000000000000 | 0.003520000000000 |
| 1.722500000000000 | 0.003720000000000 |
| 1.725000000000000 | 0.003790000000000 |
| 1.727500000000000 | 0.003800000000000 |
| 1.730000000000000 | 0.003740000000000 |
| 1.732500000000000 | 0.003860000000000 |
| 1.735000000000000 | 0.003600000000000 |
| 1.737500000000000 | 0.003600000000000 |
| 1.740000000000000 | 0.003710000000000 |
| 1.742500000000000 | 0.003690000000000 |
| 1.745000000000000 | 0.004130000000000 |
| 1.747500000000000 | 0.004210000000000 |
| 1.750000000000000 | 0.004160000000000 |
| 1.752500000000000 | 0.003850000000000 |
| 1.755000000000000 | 0.003740000000000 |
| 1.757500000000000 | 0.003830000000000 |
| 1.760000000000000 | 0.003530000000000 |
| 1.762500000000000 | 0.003630000000000 |
| 1.765000000000000 | 0.003490000000000 |
| 1.767500000000000 | 0.003520000000000 |
| 1.770000000000000 | 0.003480000000000 |
| 1.772500000000000 | 0.003380000000000 |
| 1.775000000000000 | 0.003530000000000 |
| 1.777500000000000 | 0.003570000000000 |
| 1.780000000000000 | 0.003650000000000 |

| | |
|---|---|
| 1.782500000000000 | 0.003800000000000 |
| 1.785000000000000 | 0.003770000000000 |
| 1.787500000000000 | 0.003850000000000 |
| 1.790000000000000 | 0.003990000000000 |
| 1.792500000000000 | 0.004060000000000 |
| 1.795000000000000 | 0.004090000000000 |
| 1.797500000000000 | 0.003890000000000 |
| 1.800000000000000 | 0.003950000000000 |
| 1.802500000000000 | 0.003800000000000 |
| 1.805000000000000 | 0.003680000000000 |
| 1.807500000000000 | 0.003660000000000 |
| 1.810000000000000 | 0.003520000000000 |
| 1.812500000000000 | 0.003680000000000 |
| 1.815000000000000 | 0.003610000000000 |
| 1.817500000000000 | 0.003980000000000 |
| 1.820000000000000 | 0.003820000000000 |
| 1.822500000000000 | 0.003730000000000 |
| 1.825000000000000 | 0.003720000000000 |
| 1.827500000000000 | 0.003650000000000 |
| 1.830000000000000 | 0.003420000000000 |
| 1.832500000000000 | 0.003630000000000 |
| 1.835000000000000 | 0.003680000000000 |
| 1.837500000000000 | 0.003570000000000 |
| 1.840000000000000 | 0.003570000000000 |
| 1.842500000000000 | 0.003570000000000 |
| 1.845000000000000 | 0.003610000000000 |
| 1.847500000000000 | 0.003800000000000 |
| 1.850000000000000 | 0.003620000000000 |
| 1.852500000000000 | 0.003610000000000 |
| 1.855000000000000 | 0.003560000000000 |
| 1.857500000000000 | 0.003640000000000 |
| 1.860000000000000 | 0.003890000000000 |
| 1.862500000000000 | 0.004190000000000 |
| 1.865000000000000 | 0.004160000000000 |
| 1.867500000000000 | 0.003970000000000 |
| 1.870000000000000 | 0.003950000000000 |
| 1.872500000000000 | 0.003830000000000 |
| 1.875000000000000 | 0.003840000000000 |

| | |
|---|---|
| 1.877500000000000 | 0.003730000000000 |
| 1.880000000000000 | 0.003770000000000 |
| 1.882500000000000 | 0.003690000000000 |
| 1.885000000000000 | 0.003750000000000 |
| 1.887500000000000 | 0.003590000000000 |
| 1.890000000000000 | 0.003680000000000 |
| 1.892500000000000 | 0.003610000000000 |
| 1.895000000000000 | 0.003740000000000 |
| 1.897500000000000 | 0.003690000000000 |
| 1.900000000000000 | 0.004020000000000 |
| 1.902500000000000 | 0.003930000000000 |
| 1.905000000000000 | 0.004010000000000 |
| 1.907500000000000 | 0.003830000000000 |
| 1.910000000000000 | 0.003810000000000 |
| 1.912500000000000 | 0.003810000000000 |
| 1.915000000000000 | 0.003700000000000 |
| 1.917500000000000 | 0.003640000000000 |
| 1.920000000000000 | 0.003700000000000 |
| 1.922500000000000 | 0.003650000000000 |
| 1.925000000000000 | 0.003500000000000 |
| 1.927500000000000 | 0.003480000000000 |
| 1.930000000000000 | 0.003510000000000 |
| 1.932500000000000 | 0.003450000000000 |
| 1.935000000000000 | 0.003520000000000 |
| 1.937500000000000 | 0.003580000000000 |
| 1.940000000000000 | 0.003510000000000 |
| 1.942500000000000 | 0.004020000000000 |
| 1.945000000000000 | 0.004030000000000 |
| 1.947500000000000 | 0.004170000000000 |
| 1.950000000000000 | 0.004100000000000 |
| 1.952500000000000 | 0.004020000000000 |
| 1.955000000000000 | 0.003890000000000 |
| 1.957500000000000 | 0.003770000000000 |
| 1.960000000000000 | 0.003860000000000 |
| 1.962500000000000 | 0.003900000000000 |
| 1.965000000000000 | 0.003740000000000 |
| 1.967500000000000 | 0.003610000000000 |
| 1.970000000000000 | 0.003750000000000 |

| | |
|---|---|
| 1.972500000000000 | 0.003500000000000 |
| 1.975000000000000 | 0.003550000000000 |
| 1.977500000000000 | 0.003450000000000 |
| 1.980000000000000 | 0.003330000000000 |
| 1.982500000000000 | 0.003380000000000 |
| 1.985000000000000 | 0.003360000000000 |
| 1.987500000000000 | 0.003520000000000 |
| 1.990000000000000 | 0.003650000000000 |
| 1.992500000000000 | 0.003680000000000 |
| 1.995000000000000 | 0.003580000000000 |
| 1.997500000000000 | 0.003690000000000 |
| 2.000000000000000 | 0.003850000000000 |
| 2.002500000000000 | 0.003810000000000 |
| 2.005000000000000 | 0.003920000000000 |
| 2.007500000000000 | 0.003890000000000 |
| 2.010000000000000 | 0.003550000000000 |
| 2.012500000000000 | 0.003630000000000 |
| 2.015000000000000 | 0.003810000000000 |
| 2.017500000000000 | 0.003540000000000 |
| 2.020000000000000 | 0.003570000000000 |
| 2.022500000000000 | 0.003460000000000 |
| 2.025000000000000 | 0.003240000000000 |
| 2.027500000000000 | 0.003440000000000 |
| 2.030000000000000 | 0.003340000000000 |
| 2.032500000000000 | 0.003360000000000 |
| 2.035000000000000 | 0.003560000000000 |
| 2.037500000000000 | 0.003630000000000 |
| 2.040000000000000 | 0.003670000000000 |
| 2.042500000000000 | 0.003620000000000 |
| 2.045000000000000 | 0.003920000000000 |
| 2.047500000000000 | 0.003780000000000 |
| 2.050000000000000 | 0.003710000000000 |
| 2.052500000000000 | 0.003620000000000 |
| 2.055000000000000 | 0.003600000000000 |
| 2.057500000000000 | 0.003720000000000 |
| 2.060000000000000 | 0.003740000000000 |
| 2.062500000000000 | 0.003880000000000 |
| 2.065000000000000 | 0.004180000000000 |

| | |
|---|---|
| 2.067500000000000 | 0.004180000000000 |
| 2.070000000000000 | 0.004210000000000 |
| 2.072500000000000 | 0.004280000000000 |
| 2.075000000000000 | 0.004160000000000 |
| 2.077500000000000 | 0.004150000000000 |
| 2.080000000000000 | 0.003960000000000 |
| 2.082500000000000 | 0.003990000000000 |
| 2.085000000000000 | 0.003850000000000 |
| 2.087500000000000 | 0.003880000000000 |
| 2.090000000000000 | 0.003700000000000 |
| 2.092500000000000 | 0.003630000000000 |
| 2.095000000000000 | 0.003490000000000 |
| 2.097500000000000 | 0.003460000000000 |
| 2.100000000000000 | 0.003440000000000 |
| 2.102500000000000 | 0.003550000000000 |
| 2.105000000000000 | 0.003660000000000 |
| 2.107500000000000 | 0.003630000000000 |
| 2.110000000000000 | 0.003510000000000 |
| 2.112500000000000 | 0.003500000000000 |
| 2.115000000000000 | 0.003500000000000 |
| 2.117500000000000 | 0.003720000000000 |
| 2.120000000000000 | 0.003680000000000 |
| 2.122500000000000 | 0.003680000000000 |
| 2.125000000000000 | 0.003500000000000 |
| 2.127500000000000 | 0.003470000000000 |
| 2.130000000000000 | 0.003320000000000 |
| 2.132500000000000 | 0.003290000000000 |
| 2.135000000000000 | 0.003290000000000 |
| 2.137500000000000 | 0.003340000000000 |
| 2.140000000000000 | 0.003230000000000 |
| 2.142500000000000 | 0.003440000000000 |
| 2.145000000000000 | 0.003600000000000 |
| 2.147500000000000 | 0.004000000000000 |
| 2.150000000000000 | 0.004150000000000 |
| 2.152500000000000 | 0.004220000000000 |
| 2.155000000000000 | 0.004230000000000 |
| 2.157500000000000 | 0.004100000000000 |
| 2.160000000000000 | 0.004150000000000 |

| | |
|---|---|
| 2.162500000000000 | 0.003940000000000 |
| 2.165000000000000 | 0.003880000000000 |
| 2.167500000000000 | 0.003770000000000 |
| 2.170000000000000 | 0.003690000000000 |
| 2.172500000000000 | 0.003670000000000 |
| 2.175000000000000 | 0.003560000000000 |
| 2.177500000000000 | 0.003500000000000 |
| 2.180000000000000 | 0.003360000000000 |
| 2.182500000000000 | 0.003410000000000 |
| 2.185000000000000 | 0.003420000000000 |
| 2.187500000000000 | 0.003380000000000 |
| 2.190000000000000 | 0.003560000000000 |
| 2.192500000000000 | 0.003680000000000 |
| 2.195000000000000 | 0.003770000000000 |
| 2.197500000000000 | 0.003830000000000 |
| 2.200000000000000 | 0.003860000000000 |
| 2.202500000000000 | 0.003830000000000 |
| 2.205000000000000 | 0.003740000000000 |
| 2.207500000000000 | 0.003600000000000 |
| 2.210000000000000 | 0.003590000000000 |
| 2.212500000000000 | 0.003460000000000 |
| 2.215000000000000 | 0.003590000000000 |
| 2.217500000000000 | 0.003610000000000 |
| 2.220000000000000 | 0.003570000000000 |
| 2.222500000000000 | 0.003490000000000 |
| 2.225000000000000 | 0.003390000000000 |
| 2.227500000000000 | 0.003420000000000 |
| 2.230000000000000 | 0.003460000000000 |
| 2.232500000000000 | 0.003490000000000 |
| 2.235000000000000 | 0.003560000000000 |
| 2.237500000000000 | 0.003800000000000 |
| 2.240000000000000 | 0.004010000000000 |
| 2.242500000000000 | 0.003870000000000 |
| 2.245000000000000 | 0.003800000000000 |
| 2.247500000000000 | 0.003720000000000 |
| 2.250000000000000 | 0.003610000000000 |
| 2.252500000000000 | 0.003450000000000 |
| 2.255000000000000 | 0.003390000000000 |

| | |
|---|---|
| 2.257500000000000 | 0.003320000000000 |
| 2.260000000000000 | 0.003210000000000 |
| 2.262500000000000 | 0.003300000000000 |
| 2.265000000000000 | 0.003450000000000 |
| 2.267500000000000 | 0.003370000000000 |
| 2.270000000000000 | 0.003480000000000 |
| 2.272500000000000 | 0.003500000000000 |
| 2.275000000000000 | 0.003870000000000 |
| 2.277500000000000 | 0.003800000000000 |
| 2.280000000000000 | 0.003840000000000 |
| 2.282500000000000 | 0.003910000000000 |
| 2.285000000000000 | 0.003800000000000 |
| 2.287500000000000 | 0.003690000000000 |
| 2.290000000000000 | 0.003680000000000 |
| 2.292500000000000 | 0.003460000000000 |
| 2.295000000000000 | 0.003450000000000 |
| 2.297500000000000 | 0.003270000000000 |
| 2.300000000000000 | 0.003280000000000 |
| 2.302500000000000 | 0.003360000000000 |
| 2.305000000000000 | 0.003350000000000 |
| 2.307500000000000 | 0.003430000000000 |
| 2.310000000000000 | 0.003610000000000 |
| 2.312500000000000 | 0.003700000000000 |
| 2.315000000000000 | 0.003740000000000 |
| 2.317500000000000 | 0.003750000000000 |
| 2.320000000000000 | 0.003760000000000 |
| 2.322500000000000 | 0.003600000000000 |
| 2.325000000000000 | 0.003520000000000 |
| 2.327500000000000 | 0.003640000000000 |
| 2.330000000000000 | 0.003640000000000 |
| 2.332500000000000 | 0.003520000000000 |
| 2.335000000000000 | 0.003390000000000 |
| 2.337500000000000 | 0.003290000000000 |
| 2.340000000000000 | 0.003200000000000 |
| 2.342500000000000 | 0.003330000000000 |
| 2.345000000000000 | 0.003370000000000 |
| 2.347500000000000 | 0.003250000000000 |
| 2.350000000000000 | 0.003580000000000 |

| | |
|---|---|
| 2.352500000000000 | 0.003720000000000 |
| 2.355000000000000 | 0.003800000000000 |
| 2.357500000000000 | 0.003960000000000 |
| 2.360000000000000 | 0.003920000000000 |
| 2.362500000000000 | 0.003760000000000 |
| 2.365000000000000 | 0.003870000000000 |
| 2.367500000000000 | 0.003630000000000 |
| 2.370000000000000 | 0.003720000000000 |
| 2.372500000000000 | 0.003630000000000 |
| 2.375000000000000 | 0.003390000000000 |
| 2.377500000000000 | 0.003380000000000 |
| 2.380000000000000 | 0.003370000000000 |
| 2.382500000000000 | 0.003240000000000 |
| 2.385000000000000 | 0.003330000000000 |
| 2.387500000000000 | 0.003480000000000 |
| 2.390000000000000 | 0.003540000000000 |
| 2.392500000000000 | 0.003600000000000 |
| 2.395000000000000 | 0.003410000000000 |
| 2.397500000000000 | 0.003460000000000 |
| 2.400000000000000 | 0.003480000000000 |
| 2.402500000000000 | 0.003670000000000 |
| 2.405000000000000 | 0.003690000000000 |
| 2.407500000000000 | 0.003410000000000 |
| 2.410000000000000 | 0.003290000000000 |
| 2.412500000000000 | 0.003370000000000 |
| 2.415000000000000 | 0.003370000000000 |
| 2.417500000000000 | 0.003330000000000 |
| 2.420000000000000 | 0.003350000000000 |
| 2.422500000000000 | 0.003530000000000 |
| 2.425000000000000 | 0.003490000000000 |
| 2.427500000000000 | 0.003660000000000 |
| 2.430000000000000 | 0.003930000000000 |
| 2.432500000000000 | 0.004170000000000 |
| 2.435000000000000 | 0.004090000000000 |
| 2.437500000000000 | 0.004140000000000 |
| 2.440000000000000 | 0.004010000000000 |
| 2.442500000000000 | 0.004000000000000 |
| 2.445000000000000 | 0.003950000000000 |

| | |
|---|---|
| 2.447500000000000 | 0.003910000000000 |
| 2.450000000000000 | 0.003730000000000 |
| 2.452500000000000 | 0.003690000000000 |
| 2.455000000000000 | 0.003450000000000 |
| 2.457500000000000 | 0.003390000000000 |
| 2.460000000000000 | 0.003390000000000 |
| 2.462500000000000 | 0.003210000000000 |
| 2.465000000000000 | 0.003250000000000 |
| 2.467500000000000 | 0.003310000000000 |
| 2.470000000000000 | 0.003320000000000 |
| 2.472500000000000 | 0.003430000000000 |
| 2.475000000000000 | 0.003570000000000 |
| 2.477500000000000 | 0.003700000000000 |
| 2.480000000000000 | 0.003700000000000 |
| 2.482500000000000 | 0.003960000000000 |
| 2.485000000000000 | 0.003990000000000 |
| 2.487500000000000 | 0.004160000000000 |
| 2.490000000000000 | 0.003830000000000 |
| 2.492500000000000 | 0.003650000000000 |
| 2.495000000000000 | 0.003620000000000 |
| 2.497500000000000 | 0.003430000000000 |
| 2.500000000000000 | 0.003180000000000 |
| 2.502500000000000 | 0.003350000000000 |
| 2.505000000000000 | 0.003480000000000 |
| 2.507500000000000 | 0.003420000000000 |
| 2.510000000000000 | 0.003650000000000 |
| 2.512500000000000 | 0.003760000000000 |
| 2.515000000000000 | 0.003890000000000 |
| 2.517500000000000 | 0.004120000000000 |
| 2.520000000000000 | 0.004130000000000 |
| 2.522500000000000 | 0.004060000000000 |
| 2.525000000000000 | 0.004130000000000 |
| 2.527500000000000 | 0.004030000000000 |
| 2.530000000000000 | 0.003890000000000 |
| 2.532500000000000 | 0.003930000000000 |
| 2.535000000000000 | 0.003730000000000 |
| 2.537500000000000 | 0.003740000000000 |
| 2.540000000000000 | 0.003630000000000 |

| | |
|---|---|
| 2.542500000000000 | 0.003350000000000 |
| 2.545000000000000 | 0.003270000000000 |
| 2.547500000000000 | 0.003180000000000 |
| 2.550000000000000 | 0.003210000000000 |
| 2.552500000000000 | 0.003280000000000 |
| 2.555000000000000 | 0.003430000000000 |
| 2.557500000000000 | 0.003390000000000 |
| 2.560000000000000 | 0.003370000000000 |
| 2.562500000000000 | 0.003380000000000 |
| 2.565000000000000 | 0.003630000000000 |
| 2.567500000000000 | 0.003530000000000 |
| 2.570000000000000 | 0.003470000000000 |
| 2.572500000000000 | 0.003470000000000 |
| 2.575000000000000 | 0.003410000000000 |
| 2.577500000000000 | 0.003320000000000 |
| 2.580000000000000 | 0.003230000000000 |
| 2.582500000000000 | 0.003260000000000 |
| 2.585000000000000 | 0.003160000000000 |
| 2.587500000000000 | 0.003270000000000 |
| 2.590000000000000 | 0.003430000000000 |
| 2.592500000000000 | 0.003450000000000 |
| 2.595000000000000 | 0.003470000000000 |
| 2.597500000000000 | 0.003740000000000 |
| 2.600000000000000 | 0.003900000000000 |
| 2.602500000000000 | 0.003860000000000 |
| 2.605000000000000 | 0.003810000000000 |
| 2.607500000000000 | 0.003850000000000 |
| 2.610000000000000 | 0.003620000000000 |
| 2.612500000000000 | 0.003640000000000 |
| 2.615000000000000 | 0.003440000000000 |
| 2.617500000000000 | 0.003450000000000 |
| 2.620000000000000 | 0.003270000000000 |
| 2.622500000000000 | 0.003240000000000 |
| 2.625000000000000 | 0.003310000000000 |
| 2.627500000000000 | 0.003260000000000 |
| 2.630000000000000 | 0.003230000000000 |
| 2.632500000000000 | 0.003440000000000 |
| 2.635000000000000 | 0.003450000000000 |

| | |
|---|---|
| 2.637500000000000 | 0.003480000000000 |
| 2.640000000000000 | 0.003660000000000 |
| 2.642500000000000 | 0.003720000000000 |
| 2.645000000000000 | 0.003840000000000 |
| 2.647500000000000 | 0.003690000000000 |
| 2.650000000000000 | 0.003510000000000 |
| 2.652500000000000 | 0.003410000000000 |
| 2.655000000000000 | 0.003350000000000 |
| 2.657500000000000 | 0.003140000000000 |
| 2.660000000000000 | 0.003230000000000 |
| 2.662500000000000 | 0.003050000000000 |
| 2.665000000000000 | 0.003250000000000 |
| 2.667500000000000 | 0.003240000000000 |
| 2.670000000000000 | 0.003190000000000 |
| 2.672500000000000 | 0.003200000000000 |
| 2.675000000000000 | 0.003220000000000 |
| 2.677500000000000 | 0.003360000000000 |
| 2.680000000000000 | 0.003370000000000 |
| 2.682500000000000 | 0.003540000000000 |
| 2.685000000000000 | 0.003680000000000 |
| 2.687500000000000 | 0.003750000000000 |
| 2.690000000000000 | 0.003610000000000 |
| 2.692500000000000 | 0.003530000000000 |
| 2.695000000000000 | 0.003610000000000 |
| 2.697500000000000 | 0.003570000000000 |
| 2.700000000000000 | 0.003610000000000 |
| 2.702500000000000 | 0.003480000000000 |
| 2.705000000000000 | 0.003790000000000 |
| 2.707500000000000 | 0.003760000000000 |
| 2.710000000000000 | 0.003700000000000 |
| 2.712500000000000 | 0.003760000000000 |
| 2.715000000000000 | 0.003870000000000 |
| 2.717500000000000 | 0.003800000000000 |
| 2.720000000000000 | 0.003800000000000 |
| 2.722500000000000 | 0.003770000000000 |
| 2.725000000000000 | 0.003660000000000 |
| 2.727500000000000 | 0.003610000000000 |
| 2.730000000000000 | 0.003470000000000 |

| | |
|---|---|
| 2.732500000000000 | 0.003390000000000 |
| 2.735000000000000 | 0.003270000000000 |
| 2.737500000000000 | 0.003120000000000 |
| 2.740000000000000 | 0.003180000000000 |
| 2.742500000000000 | 0.003040000000000 |
| 2.745000000000000 | 0.003150000000000 |
| 2.747500000000000 | 0.003090000000000 |
| 2.750000000000000 | 0.003130000000000 |
| 2.752500000000000 | 0.003210000000000 |
| 2.755000000000000 | 0.003190000000000 |
| 2.757500000000000 | 0.003160000000000 |
| 2.760000000000000 | 0.003270000000000 |
| 2.762500000000000 | 0.003240000000000 |
| 2.765000000000000 | 0.003240000000000 |
| 2.767500000000000 | 0.003150000000000 |
| 2.770000000000000 | 0.003130000000000 |
| 2.772500000000000 | 0.003280000000000 |
| 2.775000000000000 | 0.003310000000000 |
| 2.777500000000000 | 0.003230000000000 |
| 2.780000000000000 | 0.003180000000000 |
| 2.782500000000000 | 0.003160000000000 |
| 2.785000000000000 | 0.003190000000000 |
| 2.787500000000000 | 0.003240000000000 |
| 2.790000000000000 | 0.003210000000000 |
| 2.792500000000000 | 0.003260000000000 |
| 2.795000000000000 | 0.003410000000000 |
| 2.797500000000000 | 0.003400000000000 |
| 2.800000000000000 | 0.003500000000000 |
| 2.802500000000000 | 0.003740000000000 |
| 2.805000000000000 | 0.003780000000000 |
| 2.807500000000000 | 0.003730000000000 |
| 2.810000000000000 | 0.003650000000000 |
| 2.812500000000000 | 0.003660000000000 |
| 2.815000000000000 | 0.003630000000000 |
| 2.817500000000000 | 0.003450000000000 |
| 2.820000000000000 | 0.003380000000000 |
| 2.822500000000000 | 0.003260000000000 |
| 2.825000000000000 | 0.003150000000000 |

| | |
|---|---|
| 2.827500000000000 | 0.003120000000000 |
| 2.830000000000000 | 0.003160000000000 |
| 2.832500000000000 | 0.003110000000000 |
| 2.835000000000000 | 0.003170000000000 |
| 2.837500000000000 | 0.003270000000000 |
| 2.840000000000000 | 0.003340000000000 |
| 2.842500000000000 | 0.003420000000000 |
| 2.845000000000000 | 0.003380000000000 |
| 2.847500000000000 | 0.003500000000000 |
| 2.850000000000000 | 0.003400000000000 |
| 2.852500000000000 | 0.003420000000000 |
| 2.855000000000000 | 0.003440000000000 |
| 2.857500000000000 | 0.003330000000000 |
| 2.860000000000000 | 0.003190000000000 |
| 2.862500000000000 | 0.003250000000000 |
| 2.865000000000000 | 0.003370000000000 |
| 2.867500000000000 | 0.003310000000000 |
| 2.870000000000000 | 0.003300000000000 |
| 2.872500000000000 | 0.003230000000000 |
| 2.875000000000000 | 0.003230000000000 |
| 2.877500000000000 | 0.003470000000000 |
| 2.880000000000000 | 0.003480000000000 |
| 2.882500000000000 | 0.003450000000000 |
| 2.885000000000000 | 0.003500000000000 |
| 2.887500000000000 | 0.003460000000000 |
| 2.890000000000000 | 0.003490000000000 |
| 2.892500000000000 | 0.003340000000000 |
| 2.895000000000000 | 0.003290000000000 |
| 2.897500000000000 | 0.003140000000000 |
| 2.900000000000000 | 0.003220000000000 |
| 2.902500000000000 | 0.003210000000000 |
| 2.905000000000000 | 0.003320000000000 |
| 2.907500000000000 | 0.003280000000000 |
| 2.910000000000000 | 0.003270000000000 |
| 2.912500000000000 | 0.003300000000000 |
| 2.915000000000000 | 0.003490000000000 |
| 2.917500000000000 | 0.003410000000000 |
| 2.920000000000000 | 0.003510000000000 |

| | |
|---|---|
| 2.922500000000000 | 0.003500000000000 |
| 2.925000000000000 | 0.003500000000000 |
| 2.927500000000000 | 0.003330000000000 |
| 2.930000000000000 | 0.003430000000000 |
| 2.932500000000000 | 0.003510000000000 |
| 2.935000000000000 | 0.003310000000000 |
| 2.937500000000000 | 0.003180000000000 |
| 2.940000000000000 | 0.003090000000000 |
| 2.942500000000000 | 0.002950000000000 |
| 2.945000000000000 | 0.002950000000000 |
| 2.947500000000000 | 0.003060000000000 |
| 2.950000000000000 | 0.002930000000000 |
| 2.952500000000000 | 0.002920000000000 |
| 2.955000000000000 | 0.003160000000000 |
| 2.957500000000000 | 0.003230000000000 |
| 2.960000000000000 | 0.003080000000000 |
| 2.962500000000000 | 0.003180000000000 |
| 2.965000000000000 | 0.003160000000000 |
| 2.967500000000000 | 0.003350000000000 |
| 2.970000000000000 | 0.003300000000000 |
| 2.972500000000000 | 0.003330000000000 |
| 2.975000000000000 | 0.003480000000000 |
| 2.977500000000000 | 0.003430000000000 |
| 2.980000000000000 | 0.003450000000000 |
| 2.982500000000000 | 0.003260000000000 |
| 2.985000000000000 | 0.003070000000000 |
| 2.987500000000000 | 0.003150000000000 |
| 2.990000000000000 | 0.003180000000000 |
| 2.992500000000000 | 0.003210000000000 |
| 2.995000000000000 | 0.003200000000000 |
| 2.997500000000000 | 0.003220000000000 |
| 3.000000000000000 | 0.003360000000000 |
| 3.005000000000000 | 0.003440000000000 |
| 3.010000000000000 | 0.003460000000000 |
| 3.015000000000000 | 0.003520000000000 |
| 3.020000000000000 | 0.003490000000000 |
| 3.025000000000000 | 0.003320000000000 |
| 3.030000000000000 | 0.003130000000000 |

| | |
|---|---|
| 3.035000000000000 | 0.003090000000000 |
| 3.040000000000000 | 0.003230000000000 |
| 3.045000000000000 | 0.003310000000000 |
| 3.050000000000000 | 0.003330000000000 |
| 3.055000000000000 | 0.003140000000000 |
| 3.060000000000000 | 0.002950000000000 |
| 3.065000000000000 | 0.003050000000000 |
| 3.070000000000000 | 0.003090000000000 |
| 3.075000000000000 | 0.003100000000000 |
| 3.080000000000000 | 0.003090000000000 |
| 3.085000000000000 | 0.002940000000000 |
| 3.090000000000000 | 0.003180000000000 |
| 3.095000000000000 | 0.003180000000000 |
| 3.100000000000000 | 0.003080000000000 |
| 3.105000000000000 | 0.003060000000000 |
| 3.110000000000000 | 0.003130000000000 |
| 3.115000000000000 | 0.003220000000000 |
| 3.120000000000000 | 0.003260000000000 |
| 3.125000000000000 | 0.003340000000000 |
| 3.130000000000000 | 0.003380000000000 |
| 3.135000000000000 | 0.003450000000000 |
| 3.140000000000000 | 0.003380000000000 |
| 3.145000000000000 | 0.003350000000000 |
| 3.150000000000000 | 0.003160000000000 |
| 3.155000000000000 | 0.002950000000000 |
| 3.160000000000000 | 0.003010000000000 |
| 3.165000000000000 | 0.003040000000000 |
| 3.170000000000000 | 0.003140000000000 |
| 3.175000000000000 | 0.003190000000000 |
| 3.180000000000000 | 0.003160000000000 |
| 3.185000000000000 | 0.003080000000000 |
| 3.190000000000000 | 0.002950000000000 |
| 3.195000000000000 | 0.003090000000000 |
| 3.200000000000000 | 0.003020000000000 |
| 3.205000000000000 | 0.003000000000000 |
| 3.210000000000000 | 0.003020000000000 |
| 3.215000000000000 | 0.003200000000000 |
| 3.220000000000000 | 0.003130000000000 |

| | |
|---|---|
| 3.225000000000000 | 0.003210000000000 |
| 3.230000000000000 | 0.003130000000000 |
| 3.235000000000000 | 0.003240000000000 |
| 3.240000000000000 | 0.003040000000000 |
| 3.245000000000000 | 0.003040000000000 |
| 3.250000000000000 | 0.003030000000000 |
| 3.255000000000000 | 0.003170000000000 |
| 3.260000000000000 | 0.003160000000000 |
| 3.265000000000000 | 0.003320000000000 |
| 3.270000000000000 | 0.003270000000000 |
| 3.275000000000000 | 0.003330000000000 |
| 3.280000000000000 | 0.003260000000000 |
| 3.285000000000000 | 0.003400000000000 |
| 3.290000000000000 | 0.003650000000000 |
| 3.295000000000000 | 0.003740000000000 |
| 3.300000000000000 | 0.003710000000000 |
| 3.305000000000000 | 0.003420000000000 |
| 3.310000000000000 | 0.003370000000000 |
| 3.315000000000000 | 0.003220000000000 |
| 3.320000000000000 | 0.003100000000000 |
| 3.325000000000000 | 0.003200000000000 |
| 3.330000000000000 | 0.003210000000000 |
| 3.335000000000000 | 0.003250000000000 |
| 3.340000000000000 | 0.003410000000000 |
| 3.345000000000000 | 0.003310000000000 |
| 3.350000000000000 | 0.003180000000000 |
| 3.355000000000000 | 0.003220000000000 |
| 3.360000000000000 | 0.003170000000000 |
| 3.365000000000000 | 0.003110000000000 |
| 3.370000000000000 | 0.003150000000000 |
| 3.375000000000000 | 0.003350000000000 |
| 3.380000000000000 | 0.003350000000000 |
| 3.385000000000000 | 0.003250000000000 |
| 3.390000000000000 | 0.003160000000000 |
| 3.395000000000000 | 0.003080000000000 |
| 3.400000000000000 | 0.003130000000000 |
| 3.405000000000000 | 0.003140000000000 |
| 3.410000000000000 | 0.003080000000000 |

| | |
|---|---|
| 3.415000000000000 | 0.002990000000000 |
| 3.420000000000000 | 0.003100000000000 |
| 3.425000000000000 | 0.003200000000000 |
| 3.430000000000000 | 0.003110000000000 |
| 3.435000000000000 | 0.002900000000000 |
| 3.440000000000000 | 0.002900000000000 |
| 3.445000000000000 | 0.003040000000000 |
| 3.450000000000000 | 0.003120000000000 |
| 3.455000000000000 | 0.003030000000000 |
| 3.460000000000000 | 0.003070000000000 |
| 3.465000000000000 | 0.003230000000000 |
| 3.470000000000000 | 0.003090000000000 |
| 3.475000000000000 | 0.002920000000000 |
| 3.480000000000000 | 0.002890000000000 |
| 3.485000000000000 | 0.003000000000000 |
| 3.490000000000000 | 0.002980000000000 |
| 3.495000000000000 | 0.003060000000000 |
| 3.500000000000000 | 0.002930000000000 |
| 3.505000000000000 | 0.002950000000000 |
| 3.510000000000000 | 0.002970000000000 |
| 3.515000000000000 | 0.003040000000000 |
| 3.520000000000000 | 0.003110000000000 |
| 3.525000000000000 | 0.003180000000000 |
| 3.530000000000000 | 0.003140000000000 |
| 3.535000000000000 | 0.003010000000000 |
| 3.540000000000000 | 0.002920000000000 |
| 3.545000000000000 | 0.003010000000000 |
| 3.550000000000000 | 0.003110000000000 |
| 3.555000000000000 | 0.003120000000000 |
| 3.560000000000000 | 0.003140000000000 |
| 3.565000000000000 | 0.003050000000000 |
| 3.570000000000000 | 0.002890000000000 |
| 3.575000000000000 | 0.002940000000000 |
| 3.580000000000000 | 0.003180000000000 |
| 3.585000000000000 | 0.003160000000000 |
| 3.590000000000000 | 0.003150000000000 |
| 3.595000000000000 | 0.002990000000000 |
| 3.600000000000000 | 0.003000000000000 |

| | |
|---|---|
| 3.605000000000000 | 0.003090000000000 |
| 3.610000000000000 | 0.002930000000000 |
| 3.615000000000000 | 0.003040000000000 |
| 3.620000000000000 | 0.003200000000000 |
| 3.625000000000000 | 0.003330000000000 |
| 3.630000000000000 | 0.003410000000000 |
| 3.635000000000000 | 0.003290000000000 |
| 3.640000000000000 | 0.003050000000000 |
| 3.645000000000000 | 0.002980000000000 |
| 3.650000000000000 | 0.002960000000000 |
| 3.655000000000000 | 0.003140000000000 |
| 3.660000000000000 | 0.003180000000000 |
| 3.665000000000000 | 0.003400000000000 |
| 3.670000000000000 | 0.003450000000000 |
| 3.675000000000000 | 0.003210000000000 |
| 3.680000000000000 | 0.003050000000000 |
| 3.685000000000000 | 0.002980000000000 |
| 3.690000000000000 | 0.002920000000000 |
| 3.695000000000000 | 0.003020000000000 |
| 3.700000000000000 | 0.003050000000000 |
| 3.705000000000000 | 0.003100000000000 |
| 3.710000000000000 | 0.003280000000000 |
| 3.715000000000000 | 0.003290000000000 |
| 3.720000000000000 | 0.002970000000000 |
| 3.725000000000000 | 0.002830000000000 |
| 3.730000000000000 | 0.003000000000000 |
| 3.735000000000000 | 0.002930000000000 |
| 3.740000000000000 | 0.002960000000000 |
| 3.745000000000000 | 0.003200000000000 |
| 3.750000000000000 | 0.003080000000000 |
| 3.755000000000000 | 0.002980000000000 |
| 3.760000000000000 | 0.002930000000000 |
| 3.765000000000000 | 0.002890000000000 |
| 3.770000000000000 | 0.003090000000000 |
| 3.775000000000000 | 0.003080000000000 |
| 3.780000000000000 | 0.003050000000000 |
| 3.785000000000000 | 0.002970000000000 |
| 3.790000000000000 | 0.003060000000000 |

| | |
|---|---|
| 3.795000000000000 | 0.003210000000000 |
| 3.800000000000000 | 0.003000000000000 |
| 3.805000000000000 | 0.003100000000000 |
| 3.810000000000000 | 0.003050000000000 |
| 3.815000000000000 | 0.003000000000000 |
| 3.820000000000000 | 0.002990000000000 |
| 3.825000000000000 | 0.003220000000000 |
| 3.830000000000000 | 0.003260000000000 |
| 3.835000000000000 | 0.003050000000000 |
| 3.840000000000000 | 0.002860000000000 |
| 3.845000000000000 | 0.002880000000000 |
| 3.850000000000000 | 0.002900000000000 |
| 3.855000000000000 | 0.002890000000000 |
| 3.860000000000000 | 0.002890000000000 |
| 3.865000000000000 | 0.003170000000000 |
| 3.870000000000000 | 0.003160000000000 |
| 3.875000000000000 | 0.003250000000000 |
| 3.880000000000000 | 0.003010000000000 |
| 3.885000000000000 | 0.002950000000000 |
| 3.890000000000000 | 0.002890000000000 |
| 3.895000000000000 | 0.003060000000000 |
| 3.900000000000000 | 0.002920000000000 |
| 3.905000000000000 | 0.002920000000000 |
| 3.910000000000000 | 0.002870000000000 |
| 3.915000000000000 | 0.003070000000000 |
| 3.920000000000000 | 0.003080000000000 |
| 3.925000000000000 | 0.002910000000000 |
| 3.930000000000000 | 0.002890000000000 |
| 3.935000000000000 | 0.002830000000000 |
| 3.940000000000000 | 0.002970000000000 |
| 3.945000000000000 | 0.003040000000000 |
| 3.950000000000000 | 0.003020000000000 |
| 3.955000000000000 | 0.002900000000000 |
| 3.960000000000000 | 0.002890000000000 |
| 3.965000000000000 | 0.002940000000000 |
| 3.970000000000000 | 0.003010000000000 |
| 3.975000000000000 | 0.003010000000000 |
| 3.980000000000000 | 0.003060000000000 |

| | |
|---|---|
| 3.985000000000000 | 0.003060000000000 |
| 3.990000000000000 | 0.003240000000000 |
| 3.995000000000000 | 0.003300000000000 |
| 4.000000000000000 | 0.003230000000000 |
| 4.005000000000000 | 0.003160000000000 |
| 4.010000000000000 | 0.002960000000000 |
| 4.015000000000000 | 0.002950000000000 |
| 4.020000000000000 | 0.003070000000000 |
| 4.025000000000000 | 0.003020000000000 |
| 4.030000000000000 | 0.003120000000000 |
| 4.035000000000000 | 0.003170000000000 |
| 4.040000000000000 | 0.003200000000000 |
| 4.045000000000000 | 0.003160000000000 |
| 4.050000000000000 | 0.003060000000000 |
| 4.055000000000000 | 0.003060000000000 |
| 4.060000000000000 | 0.003100000000000 |
| 4.065000000000000 | 0.003010000000000 |
| 4.070000000000000 | 0.003080000000000 |
| 4.075000000000000 | 0.003020000000000 |
| 4.080000000000000 | 0.003010000000000 |
| 4.085000000000000 | 0.003030000000000 |
| 4.090000000000000 | 0.003070000000000 |
| 4.095000000000000 | 0.003070000000000 |
| 4.100000000000000 | 0.002870000000000 |
| 4.105000000000000 | 0.002880000000000 |
| 4.110000000000000 | 0.002870000000000 |
| 4.115000000000000 | 0.002910000000000 |
| 4.120000000000000 | 0.002940000000000 |
| 4.125000000000000 | 0.003000000000000 |
| 4.130000000000000 | 0.002820000000000 |
| 4.135000000000000 | 0.002980000000000 |
| 4.140000000000000 | 0.002920000000000 |
| 4.145000000000000 | 0.002980000000000 |
| 4.150000000000000 | 0.003110000000000 |
| 4.155000000000000 | 0.003160000000000 |
| 4.160000000000000 | 0.003110000000000 |
| 4.165000000000000 | 0.003000000000000 |
| 4.170000000000000 | 0.003060000000000 |

| | |
|---|---|
| 4.175000000000000 | 0.002990000000000 |
| 4.180000000000000 | 0.002870000000000 |
| 4.185000000000000 | 0.002860000000000 |
| 4.190000000000000 | 0.002950000000000 |
| 4.195000000000000 | 0.003040000000000 |
| 4.200000000000000 | 0.003110000000000 |
| 4.205000000000000 | 0.003040000000000 |
| 4.210000000000000 | 0.003000000000000 |
| 4.215000000000000 | 0.002850000000000 |
| 4.220000000000000 | 0.003000000000000 |
| 4.225000000000000 | 0.002980000000000 |
| 4.230000000000000 | 0.002910000000000 |
| 4.235000000000000 | 0.003030000000000 |
| 4.240000000000000 | 0.003030000000000 |
| 4.245000000000000 | 0.003060000000000 |
| 4.250000000000000 | 0.002990000000000 |
| 4.255000000000000 | 0.003100000000000 |
| 4.260000000000000 | 0.002950000000000 |
| 4.265000000000000 | 0.002960000000000 |
| 4.270000000000000 | 0.002880000000000 |
| 4.275000000000000 | 0.003060000000000 |
| 4.280000000000000 | 0.002900000000000 |
| 4.285000000000000 | 0.002950000000000 |
| 4.290000000000000 | 0.003020000000000 |
| 4.295000000000000 | 0.003030000000000 |
| 4.300000000000000 | 0.003040000000000 |
| 4.305000000000000 | 0.002880000000000 |
| 4.310000000000000 | 0.002860000000000 |
| 4.315000000000000 | 0.002850000000000 |
| 4.320000000000000 | 0.002840000000000 |
| 4.325000000000000 | 0.002830000000000 |
| 4.330000000000000 | 0.003000000000000 |
| 4.335000000000000 | 0.002850000000000 |
| 4.340000000000000 | 0.002730000000000 |
| 4.345000000000000 | 0.002870000000000 |
| 4.350000000000000 | 0.002840000000000 |
| 4.355000000000000 | 0.002890000000000 |
| 4.360000000000000 | 0.003030000000000 |

| | |
|---|---|
| 4.365000000000000 | 0.003070000000000 |
| 4.370000000000000 | 0.002950000000000 |
| 4.375000000000000 | 0.002890000000000 |
| 4.380000000000000 | 0.002980000000000 |
| 4.385000000000000 | 0.002760000000000 |
| 4.390000000000000 | 0.002830000000000 |
| 4.395000000000000 | 0.002940000000000 |
| 4.400000000000000 | 0.002990000000000 |
| 4.405000000000000 | 0.003080000000000 |
| 4.410000000000000 | 0.003030000000000 |
| 4.415000000000000 | 0.003130000000000 |
| 4.420000000000000 | 0.002970000000000 |
| 4.425000000000000 | 0.002940000000000 |
| 4.430000000000000 | 0.002880000000000 |
| 4.435000000000000 | 0.002920000000000 |
| 4.440000000000000 | 0.002880000000000 |
| 4.445000000000000 | 0.002900000000000 |
| 4.450000000000000 | 0.003030000000000 |
| 4.455000000000000 | 0.002950000000000 |
| 4.460000000000000 | 0.002850000000000 |
| 4.465000000000000 | 0.002850000000000 |
| 4.470000000000000 | 0.002870000000000 |
| 4.475000000000000 | 0.002860000000000 |
| 4.480000000000000 | 0.002950000000000 |
| 4.485000000000000 | 0.002920000000000 |
| 4.490000000000000 | 0.003100000000000 |
| 4.495000000000000 | 0.003100000000000 |
| 4.500000000000000 | 0.003120000000000 |
| 4.505000000000000 | 0.003070000000000 |
| 4.510000000000000 | 0.002930000000000 |
| 4.515000000000000 | 0.002990000000000 |
| 4.520000000000000 | 0.002790000000000 |
| 4.525000000000000 | 0.003010000000000 |
| 4.530000000000000 | 0.003040000000000 |
| 4.535000000000000 | 0.003090000000000 |
| 4.540000000000000 | 0.002870000000000 |
| 4.545000000000000 | 0.002810000000000 |
| 4.550000000000000 | 0.002880000000000 |

| | |
|---|---|
| 4.555000000000000 | 0.002900000000000 |
| 4.560000000000000 | 0.003020000000000 |
| 4.565000000000000 | 0.002930000000000 |
| 4.570000000000000 | 0.002980000000000 |
| 4.575000000000000 | 0.003160000000000 |
| 4.580000000000000 | 0.003110000000000 |
| 4.585000000000000 | 0.003100000000000 |
| 4.590000000000000 | 0.003000000000000 |
| 4.595000000000000 | 0.003000000000000 |
| 4.600000000000000 | 0.002980000000000 |
| 4.605000000000000 | 0.003090000000000 |
| 4.610000000000000 | 0.003140000000000 |
| 4.615000000000000 | 0.003050000000000 |
| 4.620000000000000 | 0.003040000000000 |
| 4.625000000000000 | 0.003010000000000 |
| 4.630000000000000 | 0.003020000000000 |
| 4.635000000000000 | 0.002950000000000 |
| 4.640000000000000 | 0.002950000000000 |
| 4.645000000000000 | 0.002930000000000 |
| 4.650000000000000 | 0.003090000000000 |
| 4.655000000000000 | 0.003130000000000 |
| 4.660000000000000 | 0.002880000000000 |
| 4.665000000000000 | 0.002970000000000 |
| 4.670000000000000 | 0.002860000000000 |
| 4.675000000000000 | 0.002920000000000 |
| 4.680000000000000 | 0.002860000000000 |
| 4.685000000000000 | 0.003060000000000 |
| 4.690000000000000 | 0.003170000000000 |
| 4.695000000000000 | 0.003080000000000 |
| 4.700000000000000 | 0.003130000000000 |
| 4.705000000000000 | 0.002970000000000 |
| 4.710000000000000 | 0.002920000000000 |
| 4.715000000000000 | 0.003030000000000 |
| 4.720000000000000 | 0.002950000000000 |
| 4.725000000000000 | 0.003130000000000 |
| 4.730000000000000 | 0.003140000000000 |
| 4.735000000000000 | 0.003090000000000 |
| 4.740000000000000 | 0.002910000000000 |

| | |
|---|---|
| 4.745000000000000 | 0.002960000000000 |
| 4.750000000000000 | 0.003040000000000 |
| 4.755000000000000 | 0.002860000000000 |
| 4.760000000000000 | 0.002920000000000 |
| 4.765000000000000 | 0.003000000000000 |
| 4.770000000000000 | 0.003200000000000 |
| 4.775000000000000 | 0.003170000000000 |
| 4.780000000000000 | 0.003000000000000 |
| 4.785000000000000 | 0.002990000000000 |
| 4.790000000000000 | 0.002930000000000 |
| 4.795000000000000 | 0.002960000000000 |
| 4.800000000000000 | 0.002990000000000 |
| 4.805000000000000 | 0.002990000000000 |
| 4.810000000000000 | 0.003150000000000 |
| 4.815000000000000 | 0.003150000000000 |
| 4.820000000000000 | 0.003030000000000 |
| 4.825000000000000 | 0.002830000000000 |
| 4.830000000000000 | 0.002850000000000 |
| 4.835000000000000 | 0.002960000000000 |
| 4.840000000000000 | 0.003040000000000 |
| 4.845000000000000 | 0.003240000000000 |
| 4.850000000000000 | 0.003300000000000 |
| 4.855000000000000 | 0.003080000000000 |
| 4.860000000000000 | 0.003060000000000 |
| 4.865000000000000 | 0.002980000000000 |
| 4.870000000000000 | 0.002880000000000 |
| 4.875000000000000 | 0.002860000000000 |
| 4.880000000000000 | 0.002910000000000 |
| 4.885000000000000 | 0.003180000000000 |
| 4.890000000000000 | 0.003270000000000 |
| 4.895000000000000 | 0.003300000000000 |
| 4.900000000000000 | 0.003150000000000 |
| 4.905000000000000 | 0.003030000000000 |
| 4.910000000000000 | 0.002850000000000 |
| 4.915000000000000 | 0.002850000000000 |
| 4.920000000000000 | 0.002960000000000 |
| 4.925000000000000 | 0.002770000000000 |
| 4.930000000000000 | 0.002910000000000 |

| | |
|---|---|
| 4.935000000000000 | 0.003030000000000 |
| 4.940000000000000 | 0.003070000000000 |
| 4.945000000000000 | 0.003030000000000 |
| 4.950000000000000 | 0.003050000000000 |
| 4.955000000000000 | 0.002840000000000 |
| 4.960000000000000 | 0.002860000000000 |
| 4.965000000000000 | 0.002910000000000 |
| 4.970000000000000 | 0.002930000000000 |
| 4.975000000000000 | 0.002940000000000 |
| 4.980000000000000 | 0.003170000000000 |
| 4.985000000000000 | 0.002950000000000 |
| 4.990000000000000 | 0.002740000000000 |
| 4.995000000000000 | 0.002850000000000 |
| 5.000000000000000 | 0.002790000000000 |
| 5.005000000000000 | 0.002950000000000 |
| 5.010000000000000 | 0.002940000000000 |
| 5.015000000000000 | 0.002870000000000 |
| 5.020000000000000 | 0.002740000000000 |
| 5.025000000000000 | 0.002740000000000 |
| 5.030000000000000 | 0.002770000000000 |
| 5.035000000000000 | 0.002790000000000 |
| 5.040000000000000 | 0.003060000000000 |
| 5.045000000000000 | 0.003170000000000 |
| 5.050000000000000 | 0.003040000000000 |
| 5.055000000000000 | 0.003060000000000 |
| 5.060000000000000 | 0.002850000000000 |
| 5.065000000000000 | 0.002900000000000 |
| 5.070000000000000 | 0.002930000000000 |
| 5.075000000000000 | 0.002690000000000 |
| 5.080000000000000 | 0.002780000000000 |
| 5.085000000000000 | 0.002810000000000 |
| 5.090000000000000 | 0.002660000000000 |
| 5.095000000000000 | 0.002880000000000 |
| 5.100000000000000 | 0.003000000000000 |
| 5.105000000000000 | 0.003040000000000 |
| 5.110000000000000 | 0.002920000000000 |
| 5.115000000000000 | 0.002860000000000 |
| 5.120000000000000 | 0.002800000000000 |

| | |
|---|---|
| 5.125000000000000 | 0.002720000000000 |
| 5.130000000000000 | 0.002680000000000 |
| 5.135000000000000 | 0.002650000000000 |
| 5.140000000000000 | 0.002940000000000 |
| 5.145000000000000 | 0.002900000000000 |
| 5.150000000000000 | 0.002890000000000 |
| 5.155000000000000 | 0.002910000000000 |
| 5.160000000000000 | 0.002820000000000 |
| 5.165000000000000 | 0.002950000000000 |
| 5.170000000000000 | 0.003020000000000 |
| 5.175000000000000 | 0.003210000000000 |
| 5.180000000000000 | 0.003110000000000 |
| 5.185000000000000 | 0.003020000000000 |
| 5.190000000000000 | 0.003000000000000 |
| 5.195000000000000 | 0.002820000000000 |
| 5.200000000000000 | 0.002860000000000 |
| 5.205000000000000 | 0.003000000000000 |
| 5.210000000000000 | 0.002960000000000 |
| 5.215000000000000 | 0.002870000000000 |
| 5.220000000000000 | 0.002840000000000 |
| 5.225000000000000 | 0.002930000000000 |
| 5.230000000000000 | 0.002980000000000 |
| 5.235000000000000 | 0.002770000000000 |
| 5.240000000000000 | 0.002920000000000 |
| 5.245000000000000 | 0.002970000000000 |
| 5.250000000000000 | 0.003060000000000 |
| 5.255000000000000 | 0.003090000000000 |
| 5.260000000000000 | 0.003010000000000 |
| 5.265000000000000 | 0.002980000000000 |
| 5.270000000000000 | 0.002800000000000 |
| 5.275000000000000 | 0.002890000000000 |
| 5.280000000000000 | 0.002890000000000 |
| 5.285000000000000 | 0.002800000000000 |
| 5.290000000000000 | 0.002920000000000 |
| 5.295000000000000 | 0.002970000000000 |
| 5.300000000000000 | 0.002910000000000 |
| 5.305000000000000 | 0.002790000000000 |
| 5.310000000000000 | 0.002790000000000 |

5.315000000000000 0.002840000000000

5.320000000000000 0.002910000000000];
