## Supplementary material for "Adaptive peak tracking as explanation of sparse fossil data across fluctuating ancient environments": MATLAB Code for all simulations

**Code for figures**

in

**across fluctuating ancient environments**

Rolf Ergon

University of South-Eastern Norway

June 2, 2025

### Figure 1

```
clear

w=100;     % Moving window size

%% Load O18 data, order and plot

load O18.mat

x=5320-x;

x=flipud(x);

u0=flipud(y);

% Plot raw and filtered data back to -2.3 million years

figure(1)

subplot(2,1,1)

xplot=0.001*x-5.3-0.02;       % -0.02 for zero point adjustment

plot(xplot,u0,'g'), hold on

u=movmean(u0,w);

x=x(737:2115);

x=0.001*x;

u=u(737:2115);

tplot=x-5.3-0.02;           % -0.2 for zero point adjustment

plot(tplot,u,'m','LineWidth',1.0), hold off, grid

ylabel('d^1^8O')

axis([-2.3 0.05 3.2 5.1])

text(-2.3,5.25,'A','FontSize',12)

% Data and plot as in Liow et al. (2024), Fig. S2, panel A

tlabel=[-2.1895 -1.9660 -1.8505 -0.9265 -0.6485 -0.5480 -0.5055 -0.4510 -0.3990  0 0];

ylabell=[11.393 11.423 11.455 11.615 11.66 11.555 11.655 11.75 11.7 11.67 11.5];

subplot(2,1,2)

err=[0.05 0.05 0.13 0.05 0.03 0.045 0.035 0.085 0.12 0.1 0.1];

errorbar(tlabel,ylabell,err,'bo'), hold on
```

```
plot(tlabel,ylabell,'b')
plot(tlabel,ylabell,'b.'), hold off, grid
ylabel('log AZ area mean')
axis([-2.3 0.05 11.29 11.84])
text(-2.3,11.885,'B','FontSize',12)
ylabel('log AZ area mean')
xlabel('Million years')
```

### Figure 3

```
clear
%% Load and order data
load O18.mat
window=100;
x=5320-x;
x=flipud(x);
u=flipud(y);
T=length(u);
N=400;
w2=50;
Gaa=2;
varv=0.5*Gaa;
Gab=0;
Gbb=2;
Gbb=0.0000001;      % Use without plasticity
Paa=Gaa+varv;
Pab=Gab;
Pbb=Gbb;
G=[Gaa Gab ; Gab Gbb ];
P=[Paa Pab ; Pab Pbb ];
%% Generate u and theta sequences
u=u-2.9*ones(1,length(u));
ufilt=movmean(u,window);
theta=u;
thetafilt=movmean(theta,window);
%% Individual population traits around abar, bbar and cbar
for t=1:T
   a(:,t)=sqrt(Gaa)*randn(N,1);
```

```
    b(:,t)=sqrt(Gbb)*randn(N,1);
    v(:,t)=sqrt(varv)*randn(N,1);
end
%% Simulation of true system
abar=0*ones(1,T);
bbar=0.8*ones(1,T);
bbar=0.0*ones(1,T);        % Use without plasticity
y=zeros(N,T);
ybar=thetafilt(1)*ones(1,T);
ybar=zeros(1,T);
Wbar=ones(1,T);
for t=1:T-1
    W(:,t)=1*exp(-(abar(t)+a(:,t)+v(:,t)+(bbar(t)+b(:,t)).*u(t)-theta(t)).^2/(2*w2));
    Wbar(t)=mean(W(:,t));
    covabW=cov([a(:,t)+v(:,t) b(:,t) W(:,t)]);
    y(:,t)=abar(t)+a(:,t)+v(:,t)+(bbar(t)+b(:,t)).*u(t);
    xbar(:,t)=[abar(t) bbar(t)]';
    xbar(:,t+1)=xbar(:,t)+G*pinv(P)*[covabW(1,3) covabW(2,3)]'/Wbar(t);
    xbarny=xbar(:,t+1);
    abar(t+1)=xbarny(1);
    bbar(t+1)=xbarny(2);
    ybar(t+1)=abar(t+1)+bbar(t+1)*u(t+1);
end
ybarfilt=movmean(ybar,window);
figure(3)
subplot(5,2,2)
plot(u,'g'), hold on, grid
plot(ybar,'b')
plot(ufilt,'m','LineWidth',1.0), hold off
axis([0 2120 -0.2 1.8])
% ylabel('u and mean(y)')
title('With plasticity')
title('Without plasticity')        % Use without plasticity
text(0,2.4,'F','FontSize',12)
subplot(5,2,4)
plot(bbar,'--b'), hold on, grid
plot(abar,'b'), hold off
axis([0 2120 -0.2 1.8])
```

```
ylabel('mean(a) and mean(b)')

text(0,2.4,'G','FontSize',12)

subplot(5,2,6)

plot(Wbar), grid

axis([0 2120 0.6 1])

% ylabel('mean(W)')

text(0,1.1,'H','FontSize',12)

subplot(5,2,8)

plot(thetafilt,'b'), hold on, grid

plot(ybarfilt,'m'), hold off

axis([0 2120 -0.2 1.5])

xlabel('time step')

ylabel('(th and mean(y))_f_i_l_t')

text(0,2.0,'I','FontSize',12)

subplot(5,2,10)

plot(ufilt,ybarfilt,'b'),hold on, grid

i=0;

for j=1:7

   i=i+200;

   plot(ufilt(i),ybarfilt(i),'bo')

end

hold off

xlabel('Environment u_f_i_l_t')

ylabel('mean(y_f_i_l_t)')

axis([0 1.5 -0.1 1.5])

text(0,2.0,'J','FontSize',12)
```

### Figure 4

```
clear

%% Load and order data

load O18.mat

window=100;

x=5320-x;

x=flipud(x);

u0=flipud(y);

u=u0;

% ufilt=movmean(u,window);
```

```
T=length(u);
N=400;
w2=50;
Gaa=2;
varv=0.5*Gaa;
Gab=0;
Gbb=2;
% Gbb=0.0000001;      % Use without plasticity
Paa=Gaa+varv;
Pab=Gab;
Pbb=Gbb;
G=[Gaa Gab ; Gab Gbb ];
P=[Paa Pab ; Pab Pbb ];
%% Generate u and theta sequences
u=u-2.9*ones(1,length(u));
ufilt=movmean(u,window);
theta=u+4*u.^2;
thetafilt=movmean(theta,window);
%% Individual population traits around abar, bbar and cbar
for t=1:T
   a(:,t)=sqrt(Gaa)*randn(N,1);
   b(:,t)=sqrt(Gbb)*randn(N,1);
   v(:,t)=sqrt(varv)*randn(N,1);
end
%% Simulation of true system
abar=0*ones(1,T);
bbar=0.8*ones(1,T);
% bbar=0.0*ones(1,T);      % Use without plasticity
y=zeros(N,T);
% ybar=thetafilt(1)*ones(1,T);
ybar=zeros(1,T);
Wbar=ones(1,T);
for t=1:T-1
   W(:,t)=1*exp(-(abar(t)+a(:,t)+v(:,t)+(bbar(t)+b(:,t)).*u(t)-theta(t)).^2/(2*w2));
   Wbar(t)=mean(W(:,t));
   covabW=cov([a(:,t)+v(:,t) b(:,t) W(:,t)]);
   y(:,t)=abar(t)+a(:,t)+v(:,t)+(bbar(t)+b(:,t)).*u(t);
   xbar(:,t)=[abar(t) bbar(t)]';
```

```
   xbar(:,t+1)=xbar(:,t)+G*pinv(P)*[covabW(1,3) covabW(2,3)]'/Wbar(t);
   xbarny=xbar(:,t+1);
   abar(t+1)=xbarny(1);
   bbar(t+1)=xbarny(2);
   ybar(t+1)=abar(t+1)+bbar(t+1)*u(t+1);
end
ybarfilt=movmean(ybar,window);
figure(4)
subplot(4,2,2)
plot(bbar,'--b'), hold on, grid
plot(abar,'b'), hold off
axis([0 2120 -5 12])
% ylabel('mean(a) and mean(b)')
text(0,15,'E','FontSize',12)
title('With plasticity')
subplot(4,2,4)
plot(Wbar), grid
axis([0 2120 0.5 1])
% ylabel('mean(W)')
text(0,1.1,'F','FontSize',12)
subplot(4,2,6)
plot(thetafilt,'b'), hold on, grid
plot(ybarfilt,'m'), hold off
axis([0 2120 -1 10])
xlabel('time step')
% ylabel('(th and mean(y))_f_i_l_t')
text(0,12,'G','FontSize',12)
subplot(4,2,8)
plot(ufilt,ybarfilt,'b'),hold on, grid
i=0;
for j=1:7
   i=i+200;
   plot(ufilt(i),ybarfilt(i),'bo')
end
hold off
xlabel('Environment u_f_i_l_t')
% ylabel('mean(y_f_i_l_t)')
axis([0 1.5 -1 10])
```

```
text(0,12,'H','FontSize',12)
```

### Figure 5

```
clear
%% Load and order data
load O18.mat
window=100;
x=5320-x;
x=flipud(x);
u0=flipud(y);
u=u0;
ufilt=movmean(u,window);
for i=1:2115
   t(i)=i;
   limit(1,t)=1;
end
figure(5)
plot(u-2.9), hold on, grid
plot(t,limit), hold off
axis([0 2115 -0.5 2.5])
ylabel('theta')
xlabel('time step')
```

### Figure 6 left panels

```
clear
%% Load and order data
load O18.mat
window=100;
x=5320-x;
x=flipud(x);
u0=flipud(y);
u=u0;
ufilt=movmean(u,window);
T=length(u);
N=1000;
w2=50;
Gaa=2;
```

```
varv=Gaa;

Gab=0;

Gbb=2;

Paa=Gaa+varv;

Pab=Gab;

Pbb=Gbb;

G=[Gaa Gab ; Gab Gbb ];

P=[Paa Pab ; Pab Pbb ];

%% Generate u and theta sequences

du=u-2.9*ones(1,length(u));

theta=du;

thetafilt=movmean(theta,window);

%% Individual population traits around abar, bbar and cbar

for t=1:T

   a(:,t)=sqrt(Gaa)*randn(N,1);

   b(:,t)=sqrt(Gbb)*randn(N,1);

   v(:,t)=sqrt(varv)*randn(N,1);

end

%% Simulation of true system

abar=0*ones(1,T);

bbar=1*ones(1,T);

y=zeros(N,T);

ybar=thetafilt(1)*ones(1,T);

Wbar=ones(1,T);

for t=1:T-1

   y(:,t)=abar(t)+a(:,t)+v(:,t)+(bbar(t)+b(:,t)).*du(t);

   for i=1:N

      if y(i,t)>1

         y(i,t)=1;

      end

   end

   vary(t)=var(y(:,t));

   W(:,t)=1*exp(-(y(:,t)-theta(t)).^2/(2*w2));

   Wtest(t)=exp(-(mean(y(:,t))-theta(t)).^2/(2*w2));

   Wbar(t)=mean(W(:,t));

   covabW=cov([a(:,t)+v(:,t) b(:,t) W(:,t)]);

   vary(t)=var(y(:,t));

   covyWmatrix=cov([y(:,t) W(:,t)]);
```

```
    covyW(t)=covyWmatrix(1,2);
    xbar(:,t)=[abar(t) bbar(t)]';
    xbar(:,t+1)=xbar(:,t)+G*pinv(P)*[covabW(1,3) covabW(2,3)]'/Wbar(t);
    xbarny=xbar(:,t+1);
    abar(t+1)=xbarny(1);
    bbar(t+1)=xbarny(2);
end
ybar=mean(y);
ybarfilt=movmean(ybar,window);
figure(6)
subplot(4,2,1)
plot(bbar,'--b'), hold on, grid
plot(abar,'b'), hold off
axis([0 2120 0 5])
ylabel('mean(a) and mean(b)')
title('Constraint example')
text(0,6,'A','FontSize',12)
subplot(4,2,3)
plot(Wbar), grid
axis([0 2120 0.98 1])
ylabel('mean(W)')
text(0,1.004,'B','FontSize',12)
subplot(4,2,5)
plot(thetafilt,'b'), hold on, grid
plot(ybarfilt,'m'), hold off
axis([0 2120 -0.1 1.7])
xlabel('time step')
ylabel('(th and mean(y))_f_i_l_t')
text(0,2.05,'C','FontSize',12)
subplot(4,2,7)
plot(ufilt,ybarfilt),hold on, grid
k=0;
for j=1:7
   k=k+200;
    plot(ufilt(k),ybarfilt(k),'bo')
end
hold off
xlabel('Environment u_f_i_l_t')
```

```
ylabel('mean(y_f_i_l_t)')

axis([2.9 4.4 0.5 1.1])

text(2.9,1.2,'D','FontSize',12)
```

### Figure 6 right panels

```
clear

%% Load and order data

load O18.mat

window=100;

x=5320-x;

x=flipud(x);

u0=flipud(y);

u=u0;

ufilt=movmean(u,window);

T=length(u);

N=1000;

w2=50;

Gaa=2;

varv=Gaa;

Gab=0;

Gbb=2;

Paa=Gaa+varv;

Pab=Gab;

Pbb=Gbb;

G=[Gaa Gab ; Gab Gbb ];

P=[Paa Pab ; Pab Pbb ];

%% Generate u and theta sequences

du=u-2.9*ones(1,length(u));

theta0=du;

for t=1:T

   theta(t)=theta0(t)+2*t/T;

end

thetafilt=movmean(theta,window);

%% Individual population traits around abar, bbar and cbar

for t=1:T

   a(:,t)=sqrt(Gaa)*randn(N,1);

   b(:,t)=sqrt(Gbb)*randn(N,1);
```

```
    v(:,t)=sqrt(varv)*randn(N,1);
end
%% Simulation of true system
abar=0*ones(1,T);
bbar=1.0*ones(1,T);
y=zeros(N,T);
ybar=thetafilt(1)*ones(1,T);
Wbar=ones(1,T);
for t=1:T-1
    y(:,t)=abar(t)+a(:,t)+v(:,t)+(bbar(t)+b(:,t)).*du(t);
    W(:,t)=1*exp(-(y(:,t)-theta(t)).^2/(2*w2));
    Wtest(t)=exp(-(mean(y(:,t))-theta(t)).^2/(2*w2));
    Wbar(t)=mean(W(:,t));
    covabW=cov([a(:,t)+v(:,t) b(:,t) W(:,t)]);
    vary(t)=var(y(:,t));
    covyWmatrix=cov([y(:,t) W(:,t)]);
    covyW(t)=covyWmatrix(1,2);
    xbar(:,t)=[abar(t) bbar(t)]';
    xbar(:,t+1)=xbar(:,t)+G*pinv(P)*[covabW(1,3) covabW(2,3)]'/Wbar(t);
    xbarny=xbar(:,t+1);
    abar(t+1)=xbarny(1);
    bbar(t+1)=xbarny(2);
end
ybar=mean(y);
ybarfilt=movmean(ybar,window);
figure(6)
subplot(4,2,2)
plot(bbar,'--b'), hold on, grid
plot(abar,'b'), hold off
axis([0 2115 -0.2 2])
title('Added theta term example')
text(0,2.5,'E','FontSize',12)
subplot(4,2,4)
plot(Wbar), grid
axis([0 2115 0.88 1])
text(0,1.025,'F','FontSize',12)
subplot(4,2,6)
plot(thetafilt,'b'), hold on, grid
```

```
plot(ybarfilt,'m'), hold off

axis([0 2115 -0.3 3.5])

xlabel('time step')

text(0,4.2,'G','FontSize',12)

subplot(4,2,8)

plot(ufilt,ybarfilt,'b'),hold on, grid

k=0;

for j=1:7

  k=k+200;

   plot(ufilt(k),ybarfilt(k),'bo')

end

hold off

xlabel('Environment u_f_i_l_t')

axis([2.9 4.4 -0.2 3.5])

text(2.9,4.2,'H','FontSize',12)
```

### Figure 8

```
clear

%% Load O18 data, order and plot

load O18.mat

x=5320-x;

x=flipud(x);

u=flipud(y);

x=0.001*x;            % Million year scale

x=x(737:2115);

u=u(737:2115);

tplot=x-5.3-0.02;       % -0.2 for zero point correction

figure(8)

subplot(3,1,1)

plot(tplot,u,'g'), hold on

ylabel('d^1^8O')

axis([-2.35 -0.3 3.1 5.1])

text(-2.35,5.35,'A','FontSize',12)

% Sample data from Liow et al. (2024)

tlabel=[-2.1895 -1.9660 -1.8505 -0.9265 -0.6485 -0.5480 -0.5055 -0.4510 -0.3990  ];

ylabell=[11.393 11.423 11.455 11.615 11.66 11.555 11.655 11.75 11.70 ];
```

```
ulabel= [mean(u(13:94))
    mean(u(122:163))
    mean(u(179:199))
    mean(u(611:620))
    mean(u(741:769))
    mean(u(816:846))
    mean(u(846:901))
    mean(u(901:955))
    mean(u(955:1005))
    ]';
plot(tplot(13:94),u(13:94),'m')
plot(tplot(122:163),u(122:163),'b')
plot(tplot(179:199),u(179:199),'m')
plot(tplot(611:621),u(611:621),'b')
plot(tplot(740:769),u(740:769),'m')
plot(tplot(816:846),u(816:846),'b')
plot(tplot(846:901),u(846:901),'m')
plot(tplot(901:955),u(901:955),'b')
plot(tplot(955:1005),u(955:1005),'m')
plot(tlabel,ulabel,'ko')
plot(tlabel,ulabel,'k.')
hold off, grid
subplot(3,1,3)
err=[0.05 0.05 0.13 0.05 0.03 0.045 0.035 0.085 0.12 ];
errorbar(ulabel,ylabell,err,'bo'), hold on, grid
errorbar(ulabel,ylabell,err,'b.')
% plot(ulabel(9:9),ylabell(9:9),'m*')
xlabel('Environment u_m_e_a_n')
ylabel('log AZ area mean')
X=[ones(9,1) ulabel'-3.6*ones(9,1)];
bls=inv(X'*X)*X'*ylabell'
a=bls(1);
b=bls(2);
yplot=bls(1)+bls(2)*(ulabel'-3.6*ones(9,1));
plot(ulabel,yplot,'m'), hold off
axis([3.58 4.74 11.28 11.87])
text(3.58,11.95,'C','FontSize',12)
for i=1:9
```

```
   yhat(i)=a+b*(ulabel(i)-3.6);
end
subplot(3,1,2)
errorbar(tlabel,ylabell,err), hold on
plot(tlabel,ylabell,'bo')
plot(tlabel,ylabell,'b.')
plot(tlabel,ylabell,'b')
plot(tlabel,yhat,'m')
plot(tlabel,yhat,'mo')
plot(tlabel,yhat,'m.'), hold off, grid
axis([-2.33 -0.3 11.28 11.87])
xlabel('Million years')
ylabel('log AZ area mean')
text(-2.33,11.95,'B','FontSize',12)
b
MSE=sum((ylabell-yhat).^2)/9
```

### Figure 9

```
clear
w=100
%% Load O18 data, order and plot
load O18.mat
x=5320-x;
x=flipud(x);
u=flipud(y);
x=0.001*x;              % Million year scale
ufilt=movmean(u,w);
x=x(737:2115);
u=u(737:2115);
ufilt=ufilt(737:2115);
tplot=x-5.3-0.02;       % -0.2 for zero point correction
figure(9)
subplot(3,1,1)
plot(tplot,u,'g'), hold on
plot(tplot,ufilt,'m','LineWidth',1.0), grid
ylabel('d^1^8O')
axis([-2.35 -0.3 3.1 5.1])
```

```
text(-2.35,5.31,'A','FontSize',12)

% Sample data from Liow et al. (2024)

tlabel=[-2.1895 -1.9660 -1.8505 -0.9265 -0.6485 -0.5480 -0.5055 -0.4510 -0.3990 ];

ylabell=[11.393 11.423 11.455 11.615 11.66 11.555 11.655 11.75 11.7 ];

ulabel=[mean(ufilt(54))

    mean(ufilt(143))

    mean(ufilt(189))

    mean(ufilt(616))

    mean(ufilt(755))

    mean(ufilt(831))

    mean(ufilt(874))

    mean(ufilt(928))

    mean(ufilt(980))

    ]';

plot(tlabel,ulabel,'ko')

plot(tlabel,ulabel,'k.')

hold off, grid

subplot(3,1,3)

err=[0.05 0.05 0.13 0.05 0.03 0.045 0.035 0.085 0.12 ];

errorbar(ulabel,ylabell,err,'bo'), hold on, grid

errorbar(ulabel,ylabell,err,'b.')

% plot(ulabel(9:9),ylabell(9:9),'m*')

xlabel('Environment u_f_i_l_t')

ylabel('log AZ area mean')

% X=[ones(9,1) ulabel'-3.6*ones(9,1)];

X=[ones(9,1) ulabel'-3.6*ones(9,1)];

bls=inv(X'*X)*X'*ylabell'

a=bls(1);

b=bls(2);

% yplot=bls(1)+bls(2)*(ulabel'-3.6*ones(10,1));

yplot=bls(1)+bls(2)*(ulabel'-3.6*ones(9,1));

plot(ulabel,yplot,'m'), hold off

axis([3.6 4.25 11.28 11.87])

text(3.6,11.95,'C','FontSize',12)

for i=1:9

   yhat(i)=a+b*(ulabel(i)-3.6);

end
```

```
for t=1:1379
   yhatplot(t)=a+b*(ufilt(t)-3.6);
   % yhat0(t)=a+b*(ufilt(t)-3.6);
end
subplot(3,1,2)
errorbar(tlabel,ylabell,err), hold on
plot(tlabel,ylabell,'bo')
plot(tlabel,ylabell,'b.')
plot(tlabel,ylabell,'b')
plot(tplot,yhatplot,'m')
plot(tlabel,yhat,'mo')
plot(tlabel,yhat,'m.'), hold off, grid
axis([-2.33 -0.3 11.28 11.87])
xlabel('Million years')
ylabel('log AZ area mean')
text(-2.33,11.95,'B','FontSize',12)
b
MSE=sum((ylabell-yhat).^2)/9
```

### Figure 10

```
clear
w=100
%% Load O18 data, order and plot
load O18.mat
x=5320-x;
x=flipud(x);
u=flipud(y);
x=0.001*x;              % Million year scale
ufilt=movmean(u,w);
x=x(737:2115);
u=u(737:2115);
ufilt=ufilt(737:2115);
tplot=x-5.3-0.02;        % -0.2 for zero point correction
% Sample data from Liow et al. (2024)
tlabel=[-2.1895 -1.9660 -1.8505 -0.9265 -0.6485 -0.5480 -0.5055 -0.4510 -0.3990];
ylabell=[11.393 11.423 11.455 11.615 11.66 11.555 11.655 11.75 11.70 ];
ylabell0=ylabell;
```

```
    ulabel=[mean(ufilt(54))
    mean(ufilt(143))
    mean(ufilt(189))
    mean(ufilt(616))
    mean(ufilt(755))
    mean(ufilt(831))
    mean(ufilt(874))
    mean(ufilt(928))
    mean(ufilt(980))
    ]';
X=[ones(9,1) ulabel'-3.6*ones(9,1)];
bls=inv(X'*X)*X'*ylabell'
a=bls(1);
b=bls(2);
% ufilt=[ufilt(1:1378) ;  mean(u(1379:1379))*ones(1,1)];
for t=1:1379
   yhatplot(t)=a+b*(ufilt(t)-3.6);
end
for i=1:9
   yhat(i)=a+b*(ulabel(i)-3.6);
end
b
MSE=sum((ylabell-yhat).^2)/9
figure(10)
err=[0.05 0.05 0.13 0.05 0.03 0.045 0.035 0.085 0.12];
errorbar(tlabel,ylabell,err,'b'), hold on
plot(tlabel,ylabell,'bo')
plot(tlabel,ylabell,'b.')
plot(tlabel,ylabell,'b')
plot(tplot,yhatplot,'m','LineWidth',1.0)
% axis([-2.33 0.05 11.28 11.87])
axis([-2.33 -0.3 11.28 11.87])
xlabel('Million years')
ylabel('log AZ area mean')
%% Validering
U=[ulabel(2:9)
   [ulabel(1) ulabel(3:9)]
```

```
   [ulabel(1:2) ulabel(4:9)]
   [ulabel(1:3) ulabel(5:9)]
   [ulabel(1:4) ulabel(6:9)]
   [ulabel(1:5) ulabel(7:9)]
   [ulabel(1:6) ulabel(8:9)]
   [ulabel(1:7) ulabel(9:9)]
   ulabel(1:8)
   ];
Y=[ylabell(2:9)
   [ylabell(1) ylabell(3:9)]
   [ylabell(1:2) ylabell(4:9)]
   [ylabell(1:3) ylabell(5:9)]
   [ylabell(1:4) ylabell(6:9)]
   [ylabell(1:5) ylabell(7:9)]
   [ylabell(1:6) ylabell(8:9)]
   [ylabell(1:7) ylabell(9:9)]
         ylabell(1:8)];
Utest=[ulabel(1)
   [ulabel(2) ]
   [ulabel(3) ]
   [ulabel(4) ]
   [ulabel(5) ]
   [ulabel(6) ]
   [ulabel(7) ]
   [ulabel(8) ]
   [ulabel(9) ]
   ];
Ytest=[ylabell(1)
   [ylabell(2) ]
   [ylabell(3) ]
   [ylabell(4) ]
   [ylabell(5) ]
   [ylabell(6) ]
   [ylabell(7) ]
   [ylabell(8) ]
   [ylabell(9) ]
   ];
```

```
for k=1:9
   ulabelcv=U(k,:);
   ylabelcv=Y(k,:);
   utest(k)=Utest(k);
   ytest(k)=Ytest(k);
X=[ones(8,1) ulabelcv'-3.6*ones(8,1)];
bls=inv(X'*X)*X'*ylabelcv';
a=bls(1);
b=bls(2);
yhattest(k)=a+b*(utest(k)-3.6);
MSE(k)=(ytest(k)-yhattest(k)).^2;
bb(k)=b;
end
bcv=sum(bb)/9
MSEcv=sum(MSE)/9
plot(tlabel,yhattest,'m.')
plot(tlabel,yhattest,'mo')
plot(tlabel,yhattest,'--m'), hold off, grid
```

### Figure 11

```
w=100;    % Moving window size
%% Load O18 data, order and plot
load O18.mat
x=5320-x;
x=flipud(x);
u0=flipud(y);
xplot=0.001*x-5.3;
% u=movmean(u0,w);
x=x(737:2115);
x=0.001*x;
u=u0(737:2115);
ufilt=movmean(u,w);
tplot=x-5.3;
% % Data and plot as in Liow et al. (2024), Fig. S2, panel A
tlabel=[-2.1895 -1.9660 -1.8505 -0.9265 -0.6485 -0.5480 -0.5055 -0.4510 -0.3990];
% ulabel=[u(54) u(143) u(189) u(616) u(755) u(831) u(874) u(928) u(980) u(1354) u(1364) ];
ylabell0=[11.393 11.423 11.455 11.615 11.66 11.555 11.655 11.75 11.7 ];
```

```
ulabel=[mean(ufilt(54))
    mean(ufilt(143))
    mean(ufilt(189))
    mean(ufilt(616))
    mean(ufilt(755))
    mean(ufilt(831))
    mean(ufilt(874))
    mean(ufilt(928))
    mean(ufilt(980))
    ]';
X=[ones(9,1) ulabel'-3.6*ones(9,1)];
bls=inv(X'*X)*X'*ylabell0';
a=bls(1);
b=bls(2);
yplot0=bls(1)+bls(2)*(ulabel'-3.6*ones(9,1));
figure(11)
subplot(2,1,1)
n=10000
yplot=zeros(n,9);
for i=1:n
ylabell(i,:)=ylabell0+[0.05 0.05 0.13 0.05 0.03 0.045 0.035 0.085 0.12 ].*randn(1,9);
bls=inv(X'*X)*X'*ylabell(i,:)';
a(i)=bls(1);
b(i)=bls(2);
bhist(i)=b(i);
yplot(i,:)=a(i)+b(i)*(ulabel-3.6*ones(1,9));
if i<101
   plot(ulabel,yplot(i,:),'c'), hold on
end
axis([3.6 4.3 11.28 11.85])
plot(ulabel,yplot0,'b')
end
err=[0.05 0.05 0.13 0.05 0.03 0.045 0.035 0.085 0.12 ];
errorbar(ulabel,ylabell0,err,'bo')
plot(ulabel,ylabell0,'b.')
ylabel('log AZ area mean')
xlabel('Environment u_f_i_l_t')
```

```
hold off, grid

subplot(2,1,2)

histogram(bhist), grid

xlabel('Slopes of prediction functions')
```
